# Integrative mapping reveals molecular features underlying the mechanism of nucleocytoplasmic transport

**DOI:** 10.1101/2023.12.31.573409

**Authors:** Barak Raveh, Roi Eliasian, Shaked Rashkovits, Daniel Russel, Ryo Hayama, Samuel E. Sparks, Digvijay Singh, Roderick Lim, Elizabeth Villa, Michael P. Rout, David Cowburn, Andrej Sali

## Abstract

Nuclear Pore Complexes (NPCs) enable rapid, selective, and robust nucleocytoplasmic transport. To explain how transport emerges from the system components and their interactions, we used experimental data and theoretical information to construct an integrative Brownian dynamics model of transport through an NPC, coupled to a kinetic model of transport in the cell. The model recapitulates key aspects of transport for a wide range of molecular cargos, including pre-ribosomes and viral capsids. It quantifies how flexible phenylalanine-glycine (FG) repeat proteins raise an entropy barrier to passive diffusion and how this barrier is selectively lowered in facilitated diffusion by the many transient interactions of nuclear transport receptors with the FG repeats. Selective transport is enhanced by “fuzzy” multivalent interactions, redundant FG repeats, coupling to the energy-dependent RanGTP concentration gradient, and exponential dependence of transport kinetics on the transport barrier. Our model will facilitate rational modulation of the NPC and its artificial mimics.

---

The Nuclear Pore Complex (NPC) mediates selective macromolecular traffic between the nucleus and cytoplasm of the eukaryotic cell, a process known as nucleocytoplasmic transport^1–3^. Being the gateway to the nucleus, the NPC is involved in most key cellular processes^4–10^. Consequently, NPC aberrations are associated with a number of disease states, disrupting vital cellular activities and leading to pathologies, such as cancer onset^11^, viral infection^12^, aging^13^, and neurodegenerative diseases^14–16^. The NPC is composed of a ring-shaped scaffold consisting of hundreds of protein subunits (nucleoporins or Nups) and a central transporter consisting of dozens of intrinsically disordered phenylalanine-glycine (FG) rich repeat domains of FG Nups, anchored inside the scaffold ring and occupying its central channel^17^ (**Fig. 1a**). The FG repeats form a size-dependent barrier, permitting rapid passive diffusion of ions, metabolites, and proteins smaller than a few dozen kDa, while impeding diffusion of larger macromolecules^18–20^. Specific large macromolecules can nonetheless overcome this diffusion barrier by forming molecular complexes with nuclear transport receptors (NTRs) that interact directly with the FG repeats^21,22^.

**Fig. 1.**
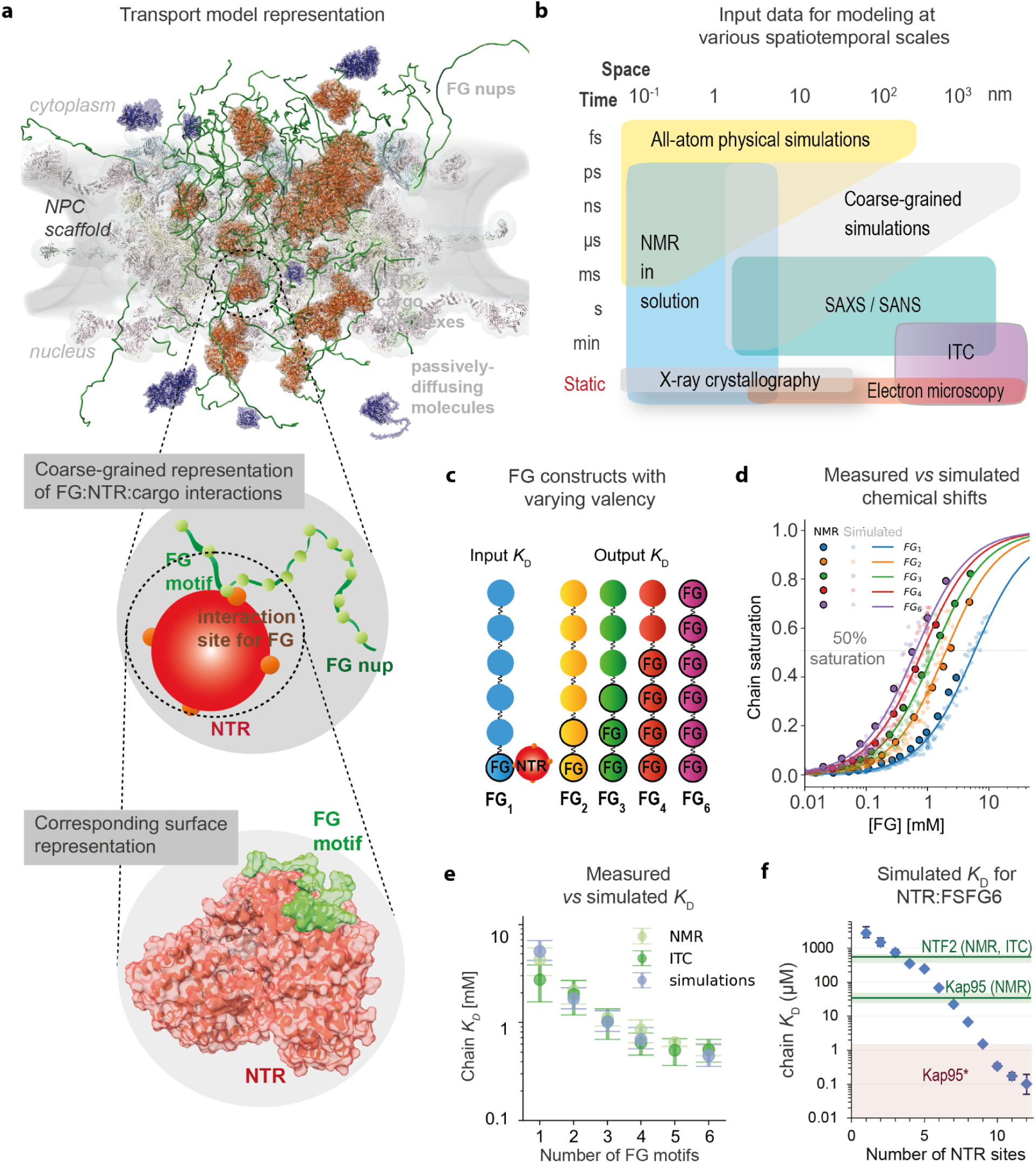
Integrative spatiotemporal modeling of nucleocytoplasmic transport. **a,** Representation of the transport model. **b,** Spatiotemporal scales of the input information for modeling. **c,** FG repeat constructs used here *in silico* and previously *in vitro*^102^ to assess the effect of their valency on the FG motif:NTR interactions. Each circle represents a segment of 20 amino acid residues. Segments labeled “FG” contain a single FSFG motif. The FSFG motifs in unlabeled segments were mutated to SSSG and therefore do not interact with NTRs. **d,** Simulated (here) and experimental^102^ titration curves of NTR chemical shifts (y-axis) as a function of the concentration of FG constructs with varying valencies (**c**; x-axis) at a fixed concentration of NTR molecules (NTF2, 10 μM). The model parameters were fitted to reproduce the data for FG_1_ and the resulting predictions were validated on FG_2_ – FG_6_ **e,** Per-chain *K*_D_ values for the interaction between an FG construct and NTF2 as a function of FG valency in the NMR or ITC experiments (light and dark green, respectively)^102^ and in the model (purple-blue). **f**, Per-chain *K_D_* values for the interaction between an FG construct with six FG motifs (5 μM) and NTRs (36 μM) as a function of NTR valency in a model for a 144 kDa NTR:cargo complex, represented as a sphere of the 35 Å radius. The horizontal green lines indicate the measured *K*_D_ values for the interaction between FG_6_ and either NTF2^102^ or Kap95^41^. The shaded pink region indicates a range of estimates of Kap95 or importin-ß affinity for various types of FG repeats ^108–111^, corresponding to higher valencies in the coarse-grained simulations.

Nucleocytoplasmic transport is complex because the transport system consists of many varied components whose interactions result in long cargo transit times on the time scale of milliseconds^17,23–27^. The mechanism underlying rapid and selective nucleocytoplasmic transport is the subject of a lively scientific debate^2,17,18,20,28–50^. The transport system and its parts have been modeled using varying representations, scales, and levels of granularity, including structural modeling of individual proteins at atomic resolution^51–53^, atomistic and coarse-grained simulations of individual components such as FG repeats and NTRs^43,54^, structural modeling of the whole NPC^17,24,25,55–58^, spatiotemporal simulations of NPC dynamics and transport at varying resolutions^2,19,20,38,39,48,59–66^, mathematical modeling of transport through an individual NPC^67,68^, and mathematical modeling of transport at the cell level^69–71^. Different explanations have been suggested regarding the dependence of transport on sequence, charge, hydrophobicity, and cohesiveness of the FG repeats^29,32,35,39,42,72,73^; the multivalency, copy number per cargo, concentration, and spatial distribution of the NTRs^27,49,74–76^; the cargo size^18–20,74,77^; the RanGTP gradient across the NE^69,70^; the flexibility of the NPC scaffold^59,60,78–80^; and the NPC’s response to mechanical stress^81,82^. Furthermore, the intrinsically disordered nature of the FG repeats allows them to be manipulated *in vitro* into different physical states, such as gels, phase-separated droplets, and brushes; these states are then interrogated for their ability to mimic aspects of nuclear transport (*eg*, refs.^30,83–85^). However, it remains unclear how accurately these presupposed states represent the NPC *in vivo*^3^. It also remains unclear how key properties of transport arise from the system components and their interactions, including how the NPC facilitates rapid and selective transport of macromolecular complexes as large as pre-ribosomal subunits, ribonucleoprotein (RNP) complexes, and viral particles^22,86–91^ robustly; the transport system is robust when perturbations of the transport system (*eg,* NPC architecture) and/or its environment (*eg*, nuclear envelope and temperature) have a limited effect on the speed and selectivity of transport. Because the NPC operates over a broad spatial and temporal scales, no single experimental method can currently yield a model that is sufficiently accurate, precise, complete, and explanatory^92^ (**Fig. 1b**). However, given the extensive structural information about the transport system components as well as the wealth of *in vivo* and *in situ* data about transport through the NPC, we hypothesized that it might be possible to use all this information, with no prior assumptions, in an integrative manner, to accurately depict and explain the process of nuclear transport.

The most accurate, precise, and complete model of a cellular process is generally computed by integrative modeling, which aims to use all relevant experimental and theoretical information across multiple spatial and temporal scales, while explicitly accounting for its sparseness, uncertainty, and ambiguity^93–98^. For example, integrative modeling has been successfully applied to compute structural models of large macromolecular assemblies that have been refractive to traditional single methods, including the NPC^17,24,25,94^. To expand the applicability of integrative modeling to even more complex and multi-scale systems, such as an entire cell, we recently developed Bayesian metamodeling^99^, an instance of integrative modeling that aims to integrate varied input models (*eg,* a set of spatiotemporal molecular trajectories and a kinetic model specified by a set of ordinary differential equations) into a metamodel consisting of updated and harmonized input models^99^.

Here, we first used integrative modeling to compute spatiotemporal trajectories of molecules diffusing through a single NPC (**Fig. 1-7**), followed by Bayesian metamodeling to couple and harmonize these trajectories with our recent kinetic model of Ran-dependent transport in an entire cell (**Fig. 8**)^71^. Importantly, our model is reductionist in the sense that nucleocytoplasmic transport arises only from the composition of the system and microscopic interactions between its components. Hence, the model avoids the pitfalls of phenomenological assumptions about key properties of the NPC and NTRs. The composition and interactions in the model were parameterized to fit a large variety of experimental data and theoretical information across a broad range of spatial and temporal scales (**Table 1**). The resulting integrative model of nucleocytoplasmic transport was then validated by its consistency with a number of observations that were not used in modeling. Strikingly, the model reveals key design features and makes quantitative predictions about nucleocytoplasmic transport, based on the system components and their interactions. For example, it rationalizes the observed central channel morphology (**Fig. 3**), explains how the NPC forms a size-dependent barrier to passively diffusing molecules (**Fig. 4**), how this barrier is overcome for diverse cargoes including megadalton complexes (**Fig. 5-8**), and how the speed and selectivity are influenced by the geometry and composition of the NPC itself (**Fig. 9**).

**Fig. 2.**
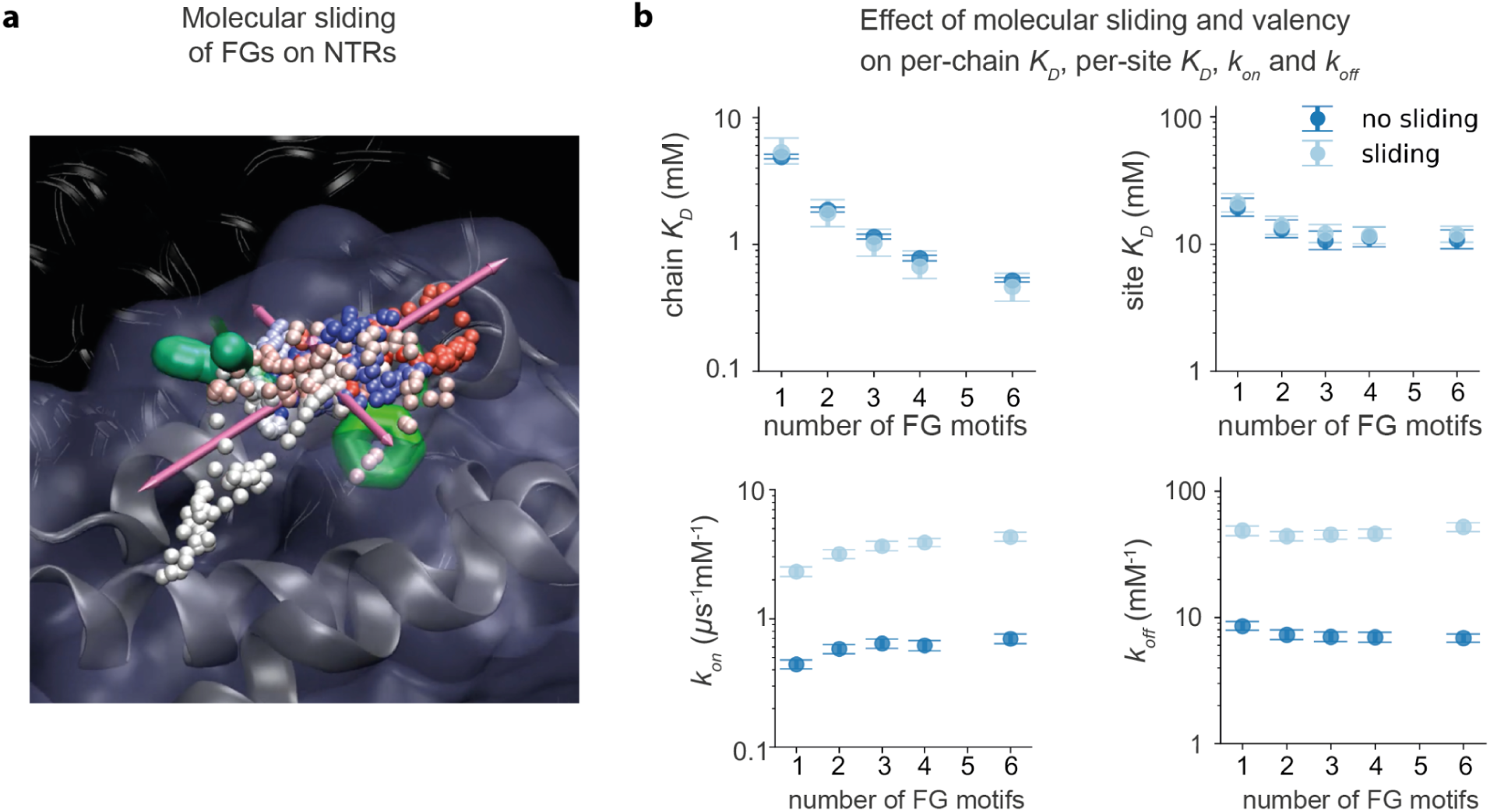
Effect of molecular sliding on FG:NTR interactions. **a**, Sliding of an FG motif interacting with an NTR, shown in a snapshot from an all-atom simulation on the Anton 2 supercomputer^117^ of a construct of six FG repeats interacting with Kap95. The pink spheres indicate the centers of mass of interacting FG motifs. The arrows indicate the principal components of the movement of these centers of mass, revealing an anisotropic sliding of the FG motifs on the interhelical groove formed between the HEAT repeats of Kap95, similar to our results for NTF2^43^. **b,** Per-chain *K*_D_, per-site *K*_D_, *k*_on_, and *k*_off_ values for the interactions between an NTR molecule (NTF2) and multivalent FG construct as a function of FG valency, with and without a sliding interaction potential. Even though the per-chain and per-site *K*_D_ values are nearly identical for the sliding and non-sliding potentials for any FG valency, the per-site exchange rates *k*_on_ and *k*_off_ are an order magnitude higher for the sliding potential than for the non-sliding potential.

**Fig. 3.**
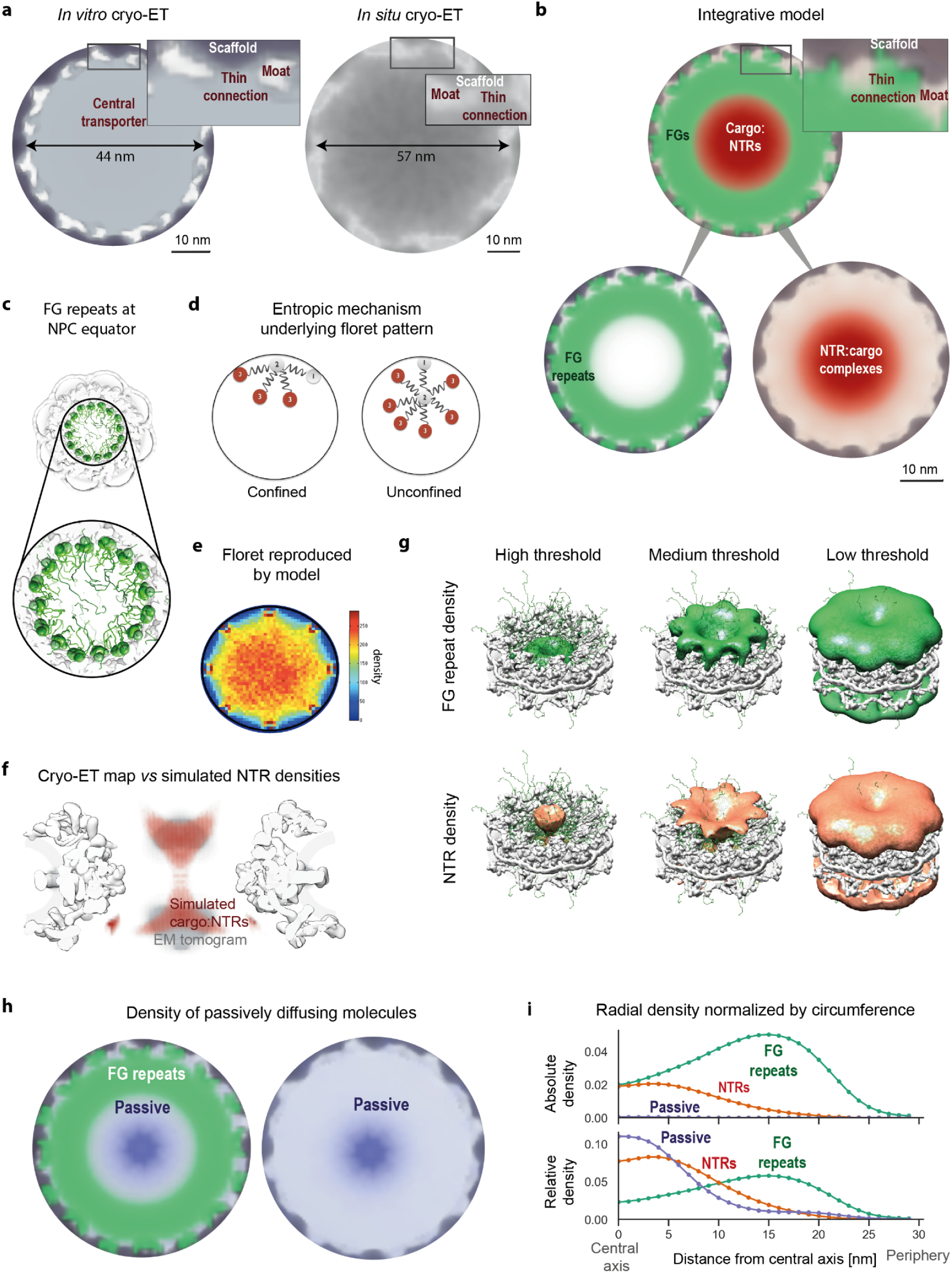
Morphology of the central transporter and spatial routes of transport. **a,** Density at the central channel (viewed from the cytoplasm) at the midpoint of the NE from the *in vitro* cryo-ET map of isolated NPCs^17^ and the current *in situ* cryo ET map, shown in gray. The scaffold density in the *in vitro* map is shown in black-gray. **b,** Same as in **a**, but for densities of the FG repeats (green) and NTRs (orange-red) near the equator of the central transporter, computed from simulated transport trajectories. The scaffold of the NPC from the cryo-ET map^17^ is shown in gray-black as in **a**, without the central transporter. **c,** The floret pattern for the density of a single layer of eight FG Nups with 15 FG repeats, each anchored to a cylinder with the radius of 20 nm, indicates that the floret in the cryo-ET map and NPC model is an inherent feature of disordered polymer chains anchored to the walls of a cylinder. **d,** Illustration of an entropic mechanism proposed to result in the floret: The number of polymer chain configurations is smaller when its stem is closer to the scaffold. **e,** Snapshot from a simulation showing FG repeat domains (green tube representation) projecting from their anchor domains (green surface) near the equator of the NPC scaffold (white surface)^17^. **f,** A side view of the NPC scaffold (white surface) with high-density lobes in the simulated NTRs (red) and in the cryo-ET map (gray)^17^. **g,** FG repeat and NTR densities contoured at three increasingly lower density thresholds, from left to right, inside the NPC scaffold^17^. **h,** Density of passively diffusing molecules (purple-blue) and FG repeats (green) at the equator of the central channel; because the density of passively diffusing molecules is low, the density threshold for the passively diffusing molecules is approximately 100 times lower than in **b**. **i,** Densities of FG repeats (green), cargo-carrying NTRs (red), and passively diffusing molecules (purple-blue) in the central transporter as a function of distance from the NPC central axis, computed from the corresponding radial densities divided by the radius, to account for the increasing circumference at the periphery. In the top panel, the densities of all three series sum to 1, whereas in the bottom panel, the densities of each series sum to 1. The overall density of passively diffusing molecules is significantly lower than that of NTRs (top); in addition, passively diffusing molecules tend to diffuse closer to the central axis of the NPC than the NTRs (bottom).

**Fig. 4.**
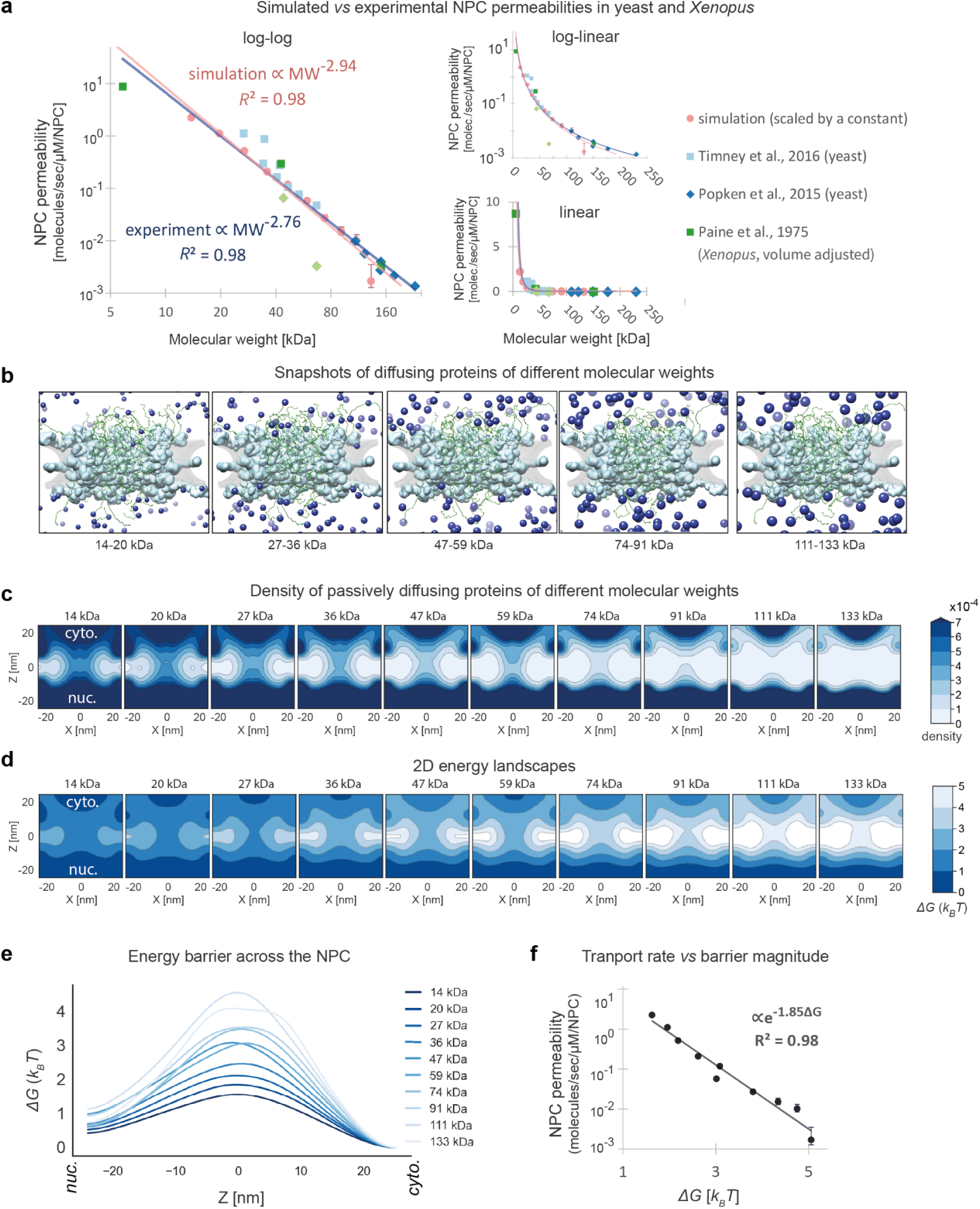
Size-dependent kinetics and thermodynamics of passive diffusion. **a,** Comparison of simulated NPC permeabilities (pink) and experimental permeabilities from *in vivo* measurements in yeast^19,20^ (blue shades) and Xenopus^88,167^ (green shades). The simulated permeability values are scaled by a single free parameter to fit the yeast data. The Xenopus data are scaled by adjusting for the significantly larger cytoplasm and nucleus volume in Xenopus (text). The pink and blue lines are power-law curves fitted to the simulated and experimental measurements in yeast, respectively. **b,** Snapshots showing passively diffusing molecules of different sizes (blue), matching the sizes for the two aligned panels in **a**. **c,** The density of passively diffusing molecules of varying molecular weights, in the *X*,*Z* plane at *Y*=0. **d,** Gibbs free energy landscape over a 2D cross-section of the NPC for passively diffusing molecules of different sizes, corresponding to the densities in **c**. **e,** Gibbs free energy for passively diffusing molecules of different sizes as a function of *Z* (integrated over the *X*,*Y* plane). **f,** NPC permeabilities as a function of the free energy barrier for passively diffusing molecules of different sizes in **e**, scaled by the same single free parameter as in **a**.

**Fig. 5.**
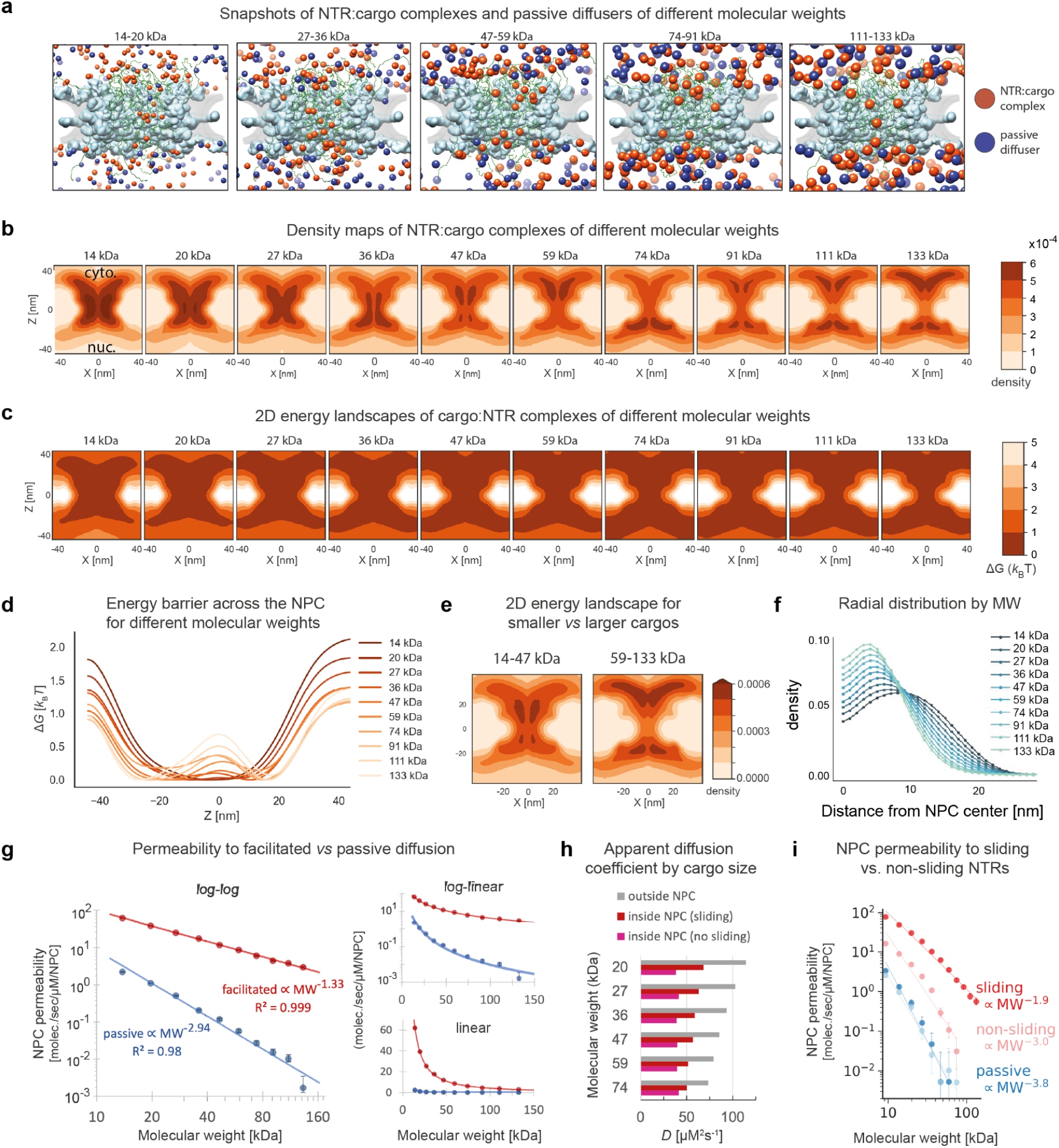
Size-dependent kinetics and thermodynamics of facilitated diffusion. **a,** A simulation snapshot showing both NTR:cargo complexes (orange-red) and passively diffusing molecules (blue) of different molecular weights. **b,** Simulated density over an *X/Z* cross-section of the NPC at *Y*=0 for NTR:cargo complexes of fixed FG valency (4 interaction sites), but different molecular weights, corresponding to those in **a**. **c**, Gibbs free energy landscape over a 2D cross-section of the NPC for NTR:cargo complexes of the different molecular weights, computed using the Boltzmann inversion of the probability densities in **b**. **d,** Gibbs free energy for NTR:cargo complexes translocating across the NPC for molecules of different weights, obtained by integrating across the *X* and *Y* dimensions of their densities followed by the Boltzmann inversion. These simulations do not include the RanGTP cycle; they quantify the permeability of the NPC in the absence of any energy exchange with the environment of the simulated system. **e,** The density of small *vs* large NTR:cargo complexes. **f,** Densities of NTR:cargo complexes of different molecular weights as a function of their distance from the NPC central axis, computed as in Fig. 2i. **g,** NPC permeability as a function of molecular weight for facilitated *vs* passive diffusion (red *vs* blue). **h,** Apparent diffusion coefficients of molecules of different molecular weights outside (gray) *vs* inside the central channel, with (red) and without sliding (magenta) of NTR:cargo complexes. **i**, NPC permeability as a function of molecular weight for sliding (red, facilitated diffusion; blue, passive diffusion) *vs* non-sliding (pink, facilitated diffusion; light blue, passive diffusion) of NTR:cargo complexes.

**Fig. 6.**
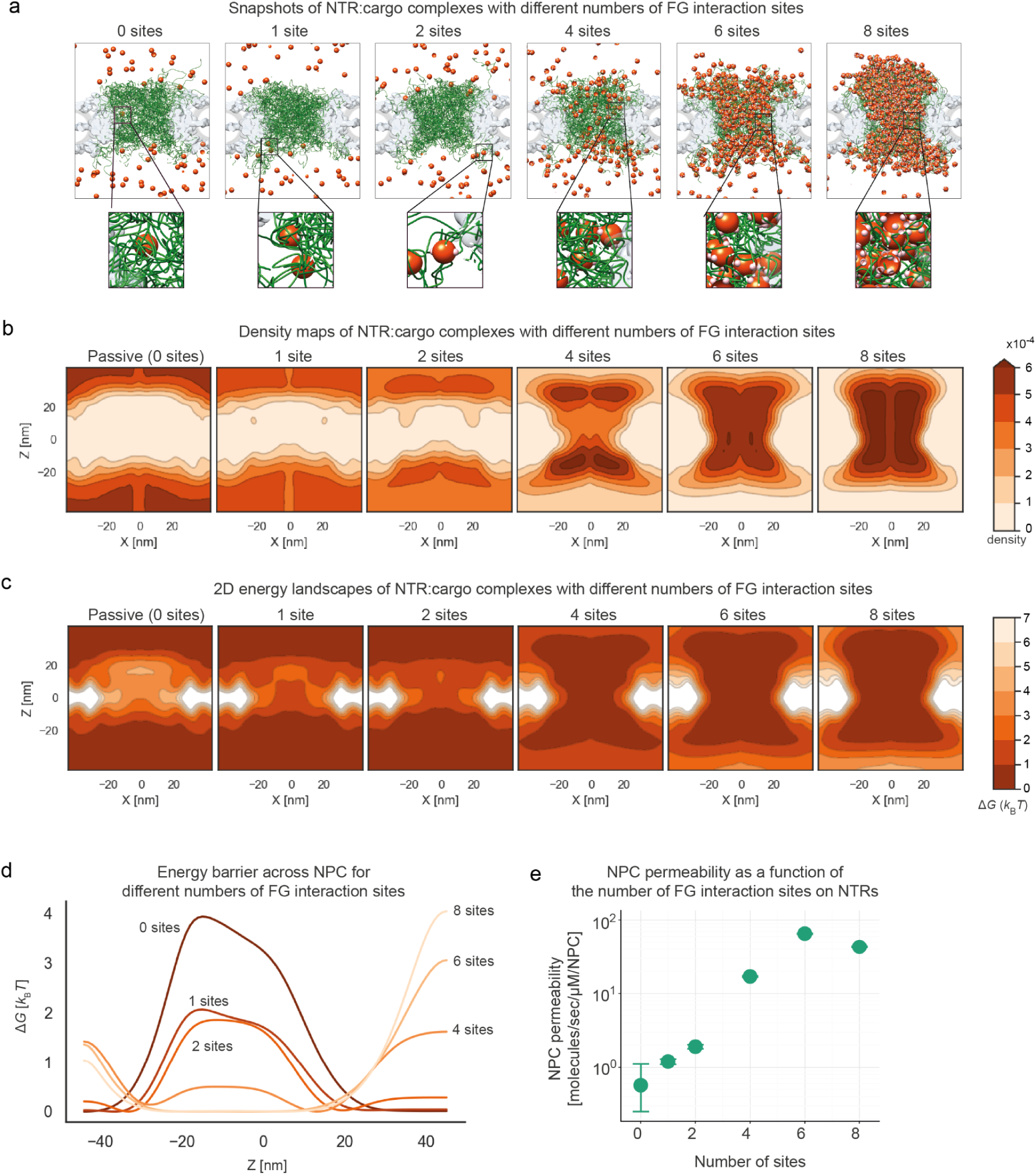
Effect of NTR valency on kinetics and thermodynamics of facilitated diffusion. **a**, A snapshot from a simulation showing NTR:cargo complexes (orange-red) with different NTR valencies (pink sites on NTRs) (*ie*, numbers of NTR interaction sites for FG motifs). **b**, Simulated densities for NTR:cargo complexes with different valencies. **c**, 2D Gibbs free energy landscapes (as in Fig. 4c) for NTR:cargo complexes with different valencies. **d**, Gibbs free energy of transport (as in Fig. 4d) for NTR:cargo complexes with different valencies. **e**, NPC permeability of the NPC to NTR:cargo complexes as a function of NTR valency.

**Fig. 7.**
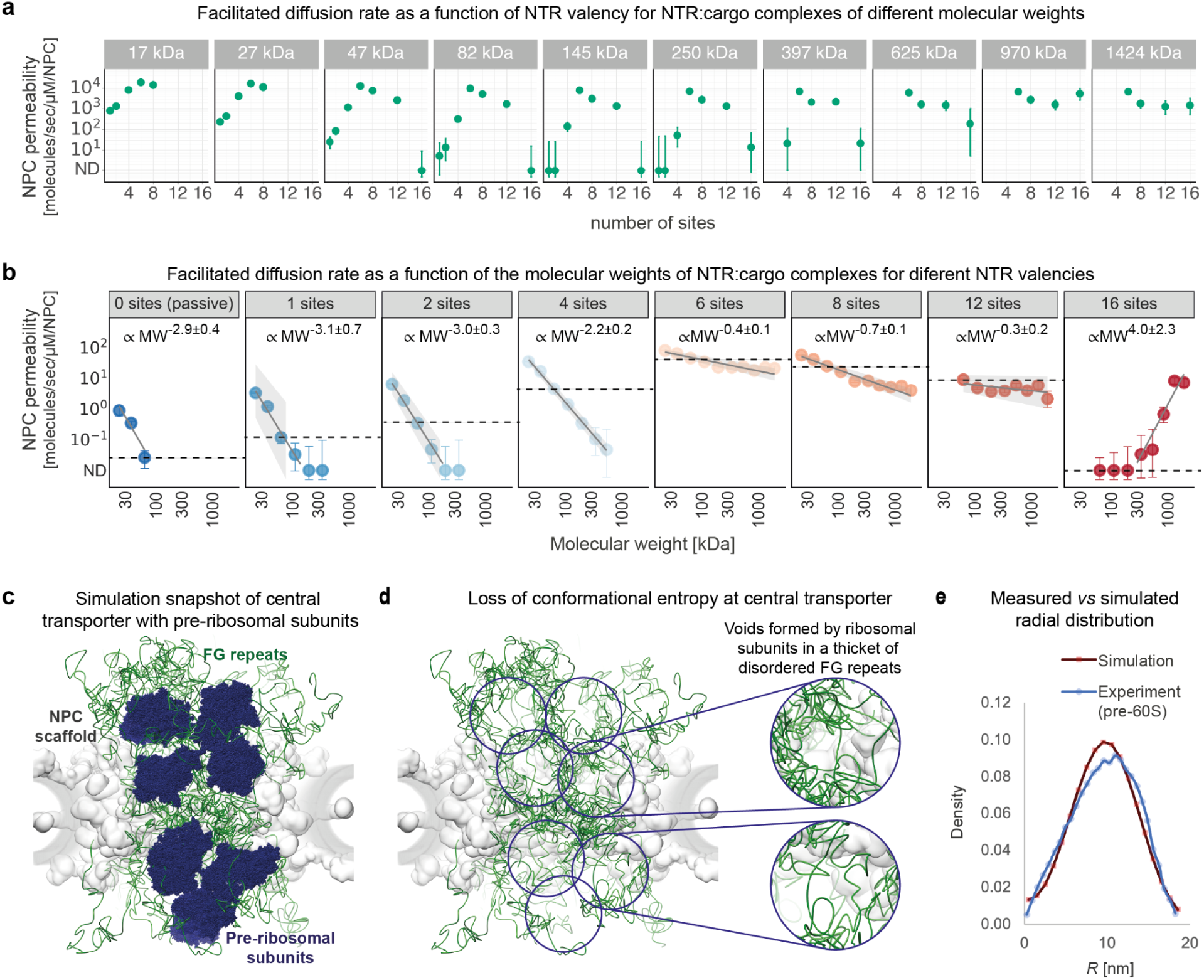
Facilitated diffusion of very large cargoes with different NTR valencies. **a**, NPC permeability as a function of molecular weight of the NTR:cargo complexes, shown for different numbers of FG-motif interaction sites on NTRs. ND (non-detectable) is the upper threshold of statistically reliable transport detection with 95% CI bars. **b**, Facilitated transport rate as a function of the number of FG motif interaction sites on NTRs, shown for different molecular weights of the NTR:cargo complexes. Gray line indicates fit to a power law (shaded area = 80% CI). Dashed black line indicates permeability for a 47 kDa NTR:cargo complex with the specified number of interaction sites. **c**, A simulation snapshot of an NPC transporting very large cargoes, equivalent in size to a late pre-40S ribosomal subunit (∼1.2 MDa). All-atom representations of pre-ribosomal subunits (PDB ID 6G18^168^) are shown at various cargo locations for illustrative purposes. **d**, Same as in **c**, without the pre-ribosomal subunits. **e**, Simulated (red line) *vs* measured (blue line) radial distribution of similarly-sized particles at the central channel (*d* = 15 nm simulated, *d* = 21 nm in experimental data on pre-60S particles^104^, radial distribution in simulations computed between *Z* = -12 nm and *Z* = 12 nm along the central axis).

**Fig. 8.**
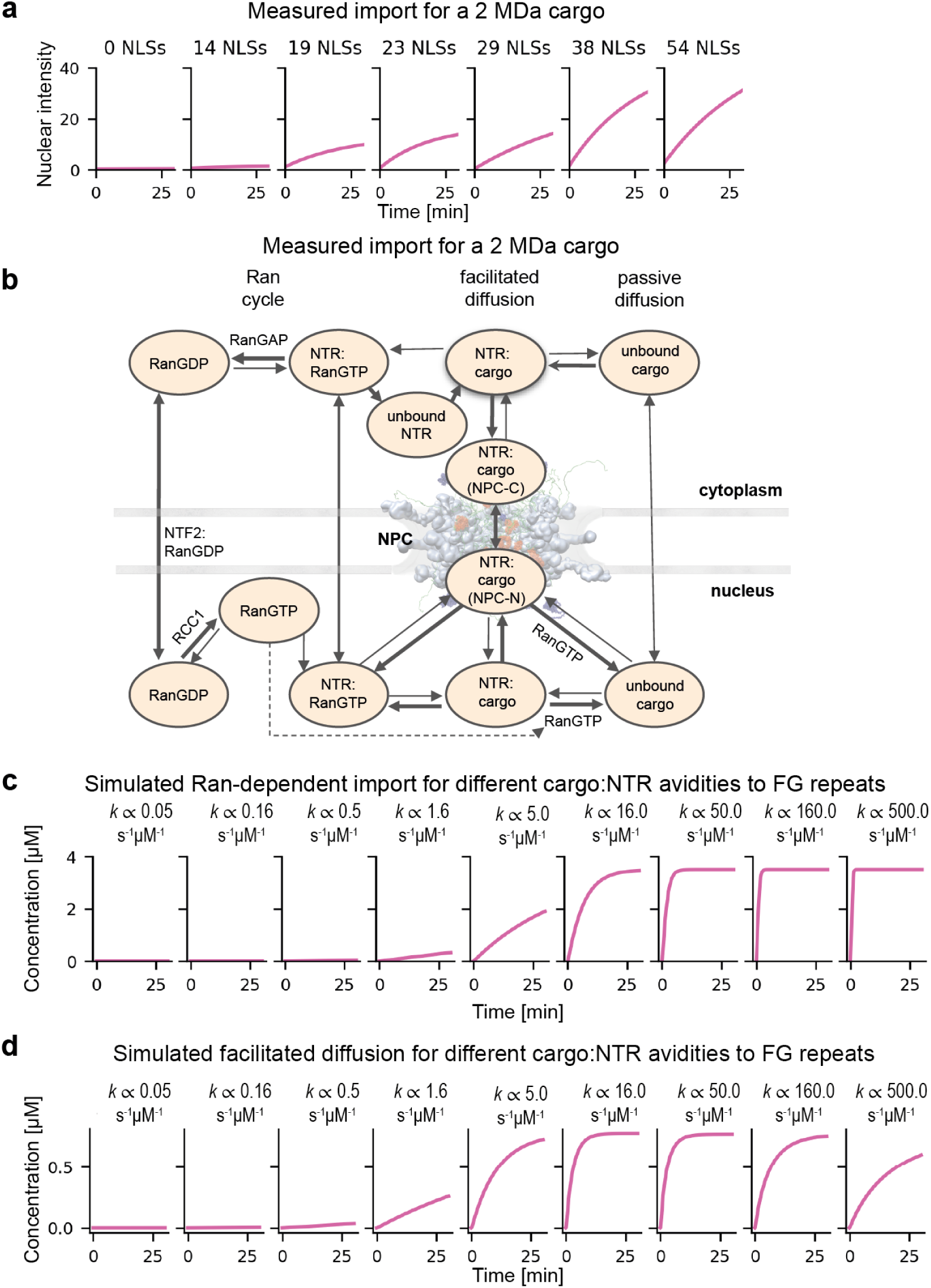
Ran-dependent transport. Experimental measurements and simulations of Ran-dependent transport and facilitated diffusion for very large cargo are compared, with the aid of a metamodel^99^ that couples our previous kinetic model of transport in the whole cell^71^ with the current spatiotemporal model of transport through a single NPC. **a**, Experimental measurements of Ran-dependent transport in permeabilized HeLa cells^74^, showing nuclear accumulation over time for very large cargo (MS2S37P; *R* = 17 nm, equivalent to a molecular weight of approximately 2 MDa), for varying numbers of Nuclear Localization Signals (NLSs) (left to right). **b**, A schematic representation of the system of ordinary differential equations describing Ran-dependent import kinetics at the whole-cell level based on a canonical description^71^. **c**, Simulations of Ran-dependent transport, showing nuclear accumulation over time for cargo of similar size, for varying magnitudes of avidity of the NTR:cargo complexes to the NPC (left to right). **d**, Same as in **c**, but for Ran-independent facilitated diffusion; here, RanGTP does not dissociate NTR:cargo complexes, which therefore dissociate at the same rate on both the nuclear and cytoplasmic sides of the NPC.

**Fig. 9.**
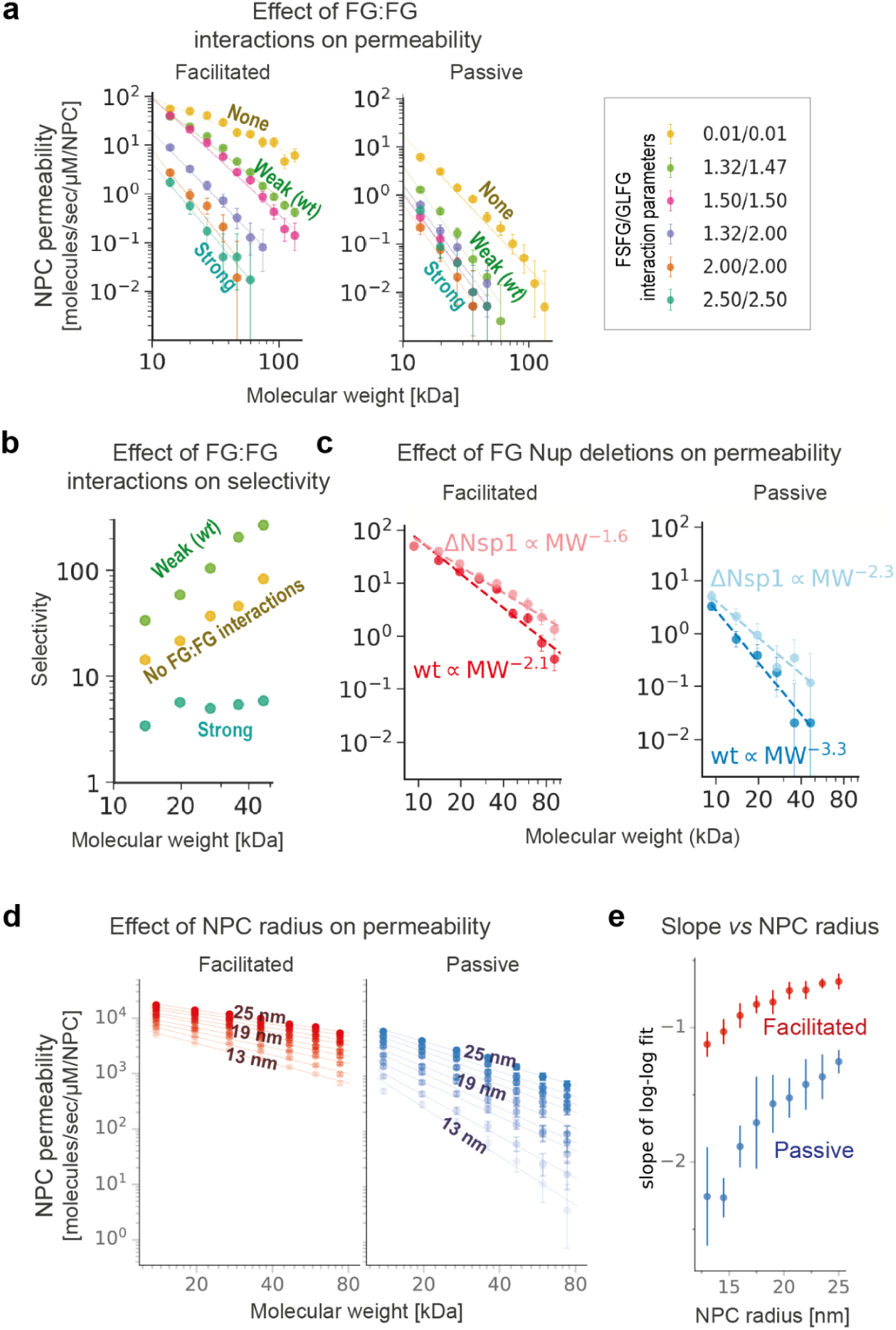
Effect of NPC perturbations on its permeability. **a,** NPC permeability as a function of molecular weight for different values of FG motif:FG motif interaction parameters. The legend indicates the cohesiveness parameter for FSFG-like and GLFG-like repeats, in units of kcal/mol/Å^2^. **b,** NPC selectivity as a function of molecular weight for different cohesiveness levels, computed by dividing the permeability for facilitated diffusion by the permeability for passive diffusion in **a**. **c,** NPC permeability as a function of molecular weight in the model of the wildtype NPC (dark points) *vs* the ΔNsp1 deletion mutant (light points) with 95% CI; lines show power-law fits (dashed lines). **d,** NPC permeability as a function of molecular weight for facilitated *vs* passive diffusion for NPCs with pore radii ranging from 13 nm to 25 nm, all else being equal, with 95% CI and power-law curve fits (lines), using a simplified representation of the NPC as a cylinder with 32 FG Nups. **e**, The slope of the fitted curves in **d** as a function of the central channel radius for facilitated (red) and passive (blue) diffusion, with 95% CI.

**Table 1.**
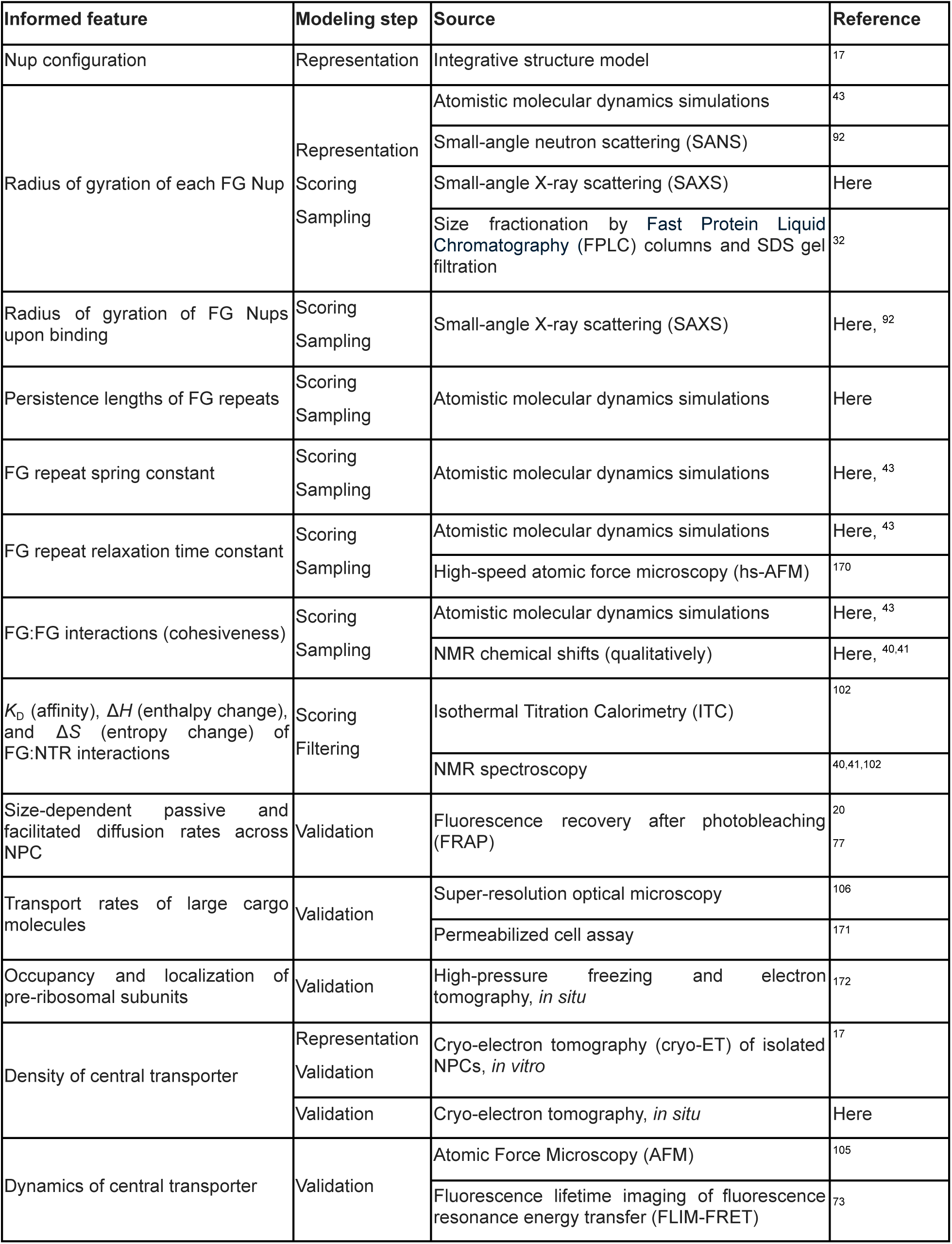
Input information for modeling. See **Supplementary Methods and Results** for details.

## Integrative spatiotemporal modeling of transport through the NPC

We begin by considering passive *versus* facilitated diffusion of macromolecular cargo through a single NPC, using integrative coarse-grained Brownian dynamics simulations (**Fig. 1a; Supplementary Video 1**). Brownian dynamics simulations have previously provided useful dynamic depictions of protein-protein interactions^100^, including in the crowded environment of the cell^101^ and the NPC^17,20,38,50^. The modeling process consisted of the following four steps (**Supplementary Methods**).

First, we gathered varied experimental and theoretical information on the structure and dynamics of the transport system and its various components (**Fig. 1b**, **Table 1**). This information included published datasets^17,32,43,92,102^ as well as new information from small-angle X-ray scattering (SAXS) profiles of FG Nups and atomistic molecular dynamics simulations of Kap95 in the presence of FSFG and GLFG repeats. We also relied on additional data for model validation, including other published datasets^19,20,73,74,103,104^ as well as new information from an *in situ* cryo-electron tomography (cryo-ET) map of the central transporter and atomic force microscopy (AFM) images of the NPC^105^. We avoided data pertaining to presupposed states of the FG repeats^3^.

Second, we created a coarse-grained Brownian dynamics representation of the transport system (**Fig. 1a**), including the nuclear envelope (NE), the scaffold of the yeast NPC embedded in the NE, the attachment sites of FG repeats to the scaffold, the dynamics of these FG repeats, NTR:cargo complexes with multiple interaction sites for FG repeats, and passively diffusing individual macromolecules that do not interact specifically with FG repeats. The integration of multiple types of information allowed us to coarse-grain both the representation of the system and its trajectories, thereby increasing modeling efficiency while retaining acceptable accuracy. Each nucleoporin was represented by a collection of beads, each bead representing approximately 20 residues. As a result, the number of simulated particles representing the FG repeats was reduced from over a million atoms (112,736 residues) to 5,488 beads.

Third, we computed a set of transport trajectories for each of the many combinations of parameter values (*eg*, interactions and diffusion constants), followed by systematically filtering these sets based on their fit to input information (*eg*, **Fig. 1c,d**, blue). Because transport is stochastic^27,106^, thousands of independent trajectories, each starting from a random configuration, were computed for each set of conditions, for an aggregate simulation time of several seconds. Several million CPU core hours on Google Exacycle and our own Linux high-performance computing clusters at UCSF and HUJI were used.

Finally, we validated each filtered set of trajectories by quantifying how well it reproduced experimental and theoretical information not used in model construction, pertaining to both the individual model components (*eg*, **Fig. 1c-f**; **Fig. 2**) and the entire transport system (*eg,* **Fig. 3a**, **Fig. 4a**, **Fig. 7e, Extended Data Fig. 2-4,6**). The next four sections present these validations.^3^

## The integrative model reproduces key biochemical characterizations of multivalent FG:NTR interactions not used during its construction

Given the established role of FG repeat – NTR:cargo interactions, we first ensured that our integrative model of nucleocytoplasmic transport accurately represents these essential components. Specifically, we simulated freely-diffusing FG repeats and NTR:cargo complexes, followed by assessing these simulations against key biochemical measurements, including those not utilized for model construction. The FG repeats were parameterized to match the known radius of gyration, end-to-end distance, and the relaxation times of these properties in a dilute solution (**Table 1**). Additionally, the interaction between FG repeats and different types of NTRs were parameterized as follows. First, the interaction between individual FG motifs and NTR molecule (NTF2) was parameterized to reproduce the corresponding dissociation constant in the dilute regime, measured by NMR titrations and Isothermal Titration Calorimetry (ITC) (**Fig. 1d**, blue; **Table 1**)^102^. Second, to validate the model of the interaction using data not used for model construction, we compared the simulated and measured avidity of multivalent FG repeat constructs with 2, 3, 4, or 6 FG motifs (**Fig. 1c-e**). The model reproduced not only the dissociation constants from NMR and ITC experiments (**Fig. 1e**), but also the raw NMR titration data quantifying the fraction of bound chains for a wide range of FG valencies and concentrations (**Fig. 1d**). In fact, the difference between the model and NMR data was smaller than that between the NMR and ITC data (**Fig. 1e**), despite a non-linear relation between the FG valencies and their NTR avidities. This non-linear relation cannot be explained by simple kinetic modeling^102^, illustrating the need for spatiotemporal modeling. Finally, measured Kap95 (importin subunit β-1) avidities span a range of values^41,107–111^; this range can be reproduced by our model simply by changing the number of interaction sites and molecular radius (**Fig. 1f**). In conclusion, the validation of the integrative model for both major families of NTRs that interact directly with the NPC^53,112–114^ indicates that the model sufficiently accurately represents the interactions between its key components, thereby increasing our confidence in other predictions below.

## The integrative model reproduces rapid exchange of interacting FG repeats and NTRs through a slide-and-exchange mechanism

FG repeats and NTRs exchange at an unusually rapid rate compared to other proteins interacting with similar affinity^40,41^. This rapid exchange can be explained by a “fuzzy” slide-and-exchange mechanism^43,92^, based on long molecular dynamics simulations and supporting NMR experiments. According to the slide-and-exchange mechanism, FG motifs slide anisotropically along NTR binding grooves, analogously to the sliding of transcription factors as they rapidly locate their binding sites on DNA^115^. As one FG motif slides into the NTR groove, it displaces the resident FG motif, which is significantly faster than the alternative of separate dissociation and association events^43,116^. However, sliding had been simulated for only one NTR (NTF2) interacting with FG repeats^43^. NTF2 represents one structurally related family of NTRs, the other family being the karyopherins, of which Kap95 is considered a canonical member. Therefore, to establish whether sliding can occur between other types of NTRs and FG repeats, we performed atomistic molecular dynamics simulations^117^ of FG repeats in the presence of Kap95. Crucially, despite the significant structural differences between Kap95 and NTF2, the simulations show that the FG motifs also slide through the grooves between adjacent Kap95 helices (**Supplementary Video 2**), as visualized by a principal component analysis of the FSFG motif center-of-mass sliding on Kap95 (**Fig. 2a**; **Supplementary Video 3**). Thus, the slide-and-exchange mechanism is substantiated by molecular dynamics simulations for representatives of both major families of NTRs^53,112–114^.

Next, we validated the contribution of the sliding of individual FG motifs and NTR molecules to their exchange rate at physiological concentrations^43^. The sliding of competing FG motifs on NTRs in the coarse-grained Brownian dynamics simulations resulted in on- and off-rates an order of magnitude higher in the sliding mode than in the non-sliding mode, despite having the same overall avidity (**Fig. 2b; Supplementary Results**). The accurate reproduction of the rapid exchange between FG repeats and NTRs^40,41^ in the context of multivalent interactions^102^ justifies using the sliding mode in our coarse-grained model. We discuss the effect of this sliding on facilitated diffusion in the context of the full NPC below.

## The integrative model reproduces the effect of anchoring on the polymer properties of FG repeats

Recent *in situ* FRET measurements indicated that the FG repeats of hNup98, the human ortholog of yeast Nup100, are significantly more extended in the NPC than in a dilute buffer^73^. Because these data were not available during the construction of the integrative model, we can use them to test how well the model captures the effect of the NPC milieu on the FG repeats. Although the underlying model was trained to reproduce the relatively low end-to-end distance (*R_E_*) of Nup100 measured in buffer (**Extended Data Figs. 1-2; Table 2**), it recapitulated the empirical increase in *R_E_* in the NPC, caused by the anchoring of multiple FG Nups to the NPC scaffold (**Extended Data Fig. 2b**). Thus, the integrative model correctly describes how the context of the NPC milieu modifies the intrinsic polymer properties of FG repeats, in agreement with the suggested transitioning from a poor-solvent to the theta-solvent regime *in situ*^73,118^.

**Table 2.**
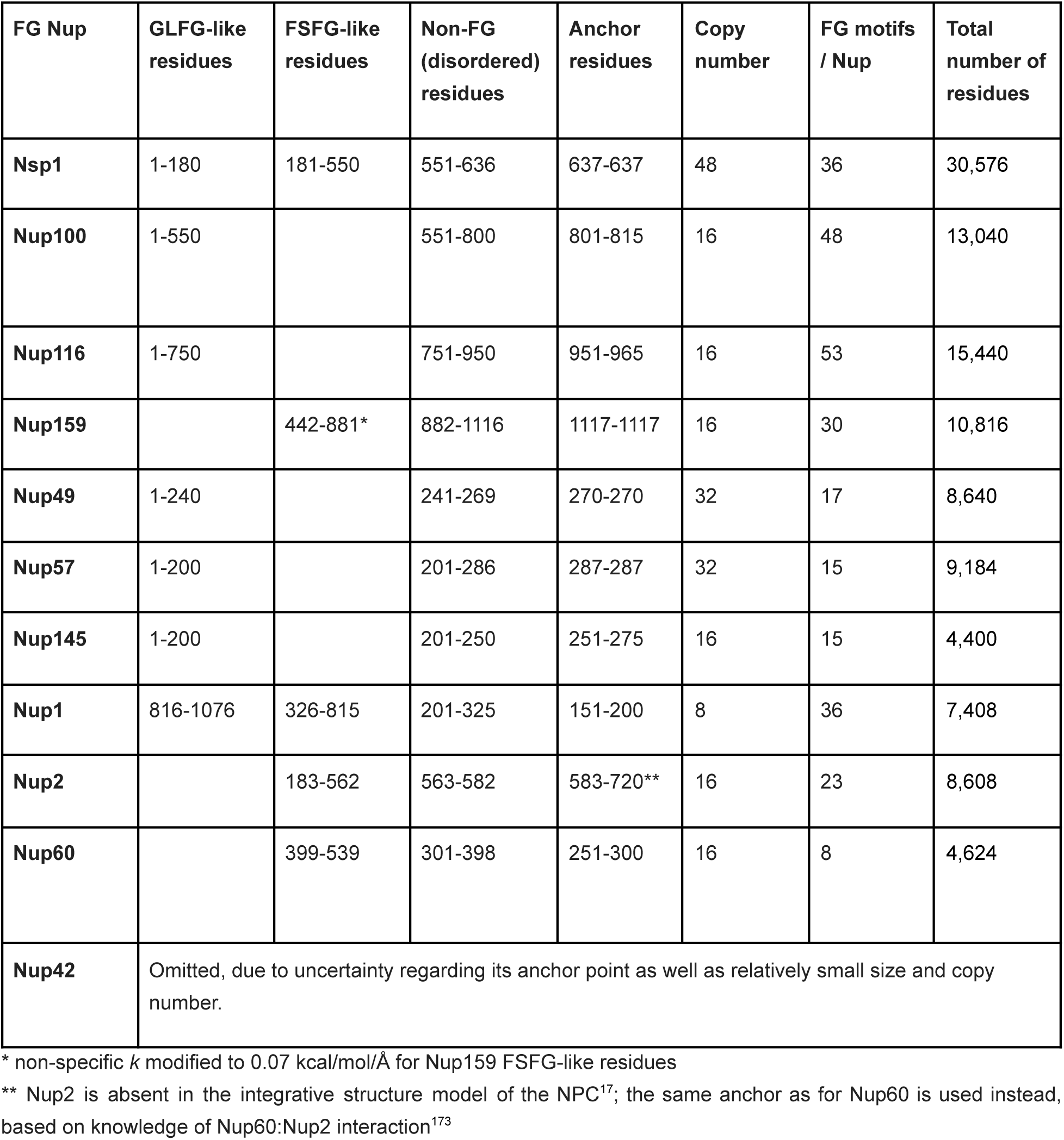
FG repeat domains of FG Nups, their flavors, and anchor points.

| FG Nup | GLFG-like residues | FSFG-like residues | Non-FG (disordered) residues | Anchor residues | Copy number | FG motifs / Nup | Total number of residues |
| --- | --- | --- | --- | --- | --- | --- | --- |
| **Nsp1** | 1-180 | 181-550 | 551-636 | 637-637 | 48 | 36 | 30,576 |
| **Nup100** | 1-550 |  | 551-800 | 801-815 | 16 | 48 | 13,040 |
| **Nup116** | 1-750 |  | 751-950 | 951-965 | 16 | 53 | 15,440 |
| **Nup159** |  | 442-881* | 882-1116 | 1117-1117 | 16 | 30 | 10,816 |
| **Nup49** | 1-240 |  | 241-269 | 270-270 | 32 | 17 | 8,640 |
| **Nup57** | 1-200 |  | 201-286 | 287-287 | 32 | 15 | 9,184 |
| **Nup145** | 1-200 |  | 201-250 | 251-275 | 16 | 15 | 4,400 |
| **Nup1** | 816-1076 | 326-815 | 201-325 | 151-200 | 8 | 36 | 7,408 |
| **Nup2** |  | 183-562 | 563-582 | 583-720** | 16 | 23 | 8,608 |
| **Nup60** |  | 399-539 | 301-398 | 251-300 | 16 | 8 | 4,624 |
| **Nup42** | Omitted, due to uncertainty regarding its anchor point as well as relatively small size and copy number. |  |  |  |  |  |  |

\* non-specific $k$ modified to 0.07 kcal/mol/Å for Nup159 FSFG-like residues
\*\* Nup2 is absent in the integrative structure model of the NPC<sup>17</sup>; the same anchor as for Nup60 is used instead, based on knowledge of Nup60:Nup2 interaction<sup>173</sup>

## The integrative model reproduces salient morphological features of the central transporter

We next assessed whether the integrative model reproduces the empirical density distribution of FG repeats, NTRs, and cargo molecules in the central channel of the NPC (*ie*, the central transporter)^17,76,105^. Originally termed the central plug^119^, the central transporter has been often omitted from electron microscopy (EM) reconstructions of the NPC, due to the heterogeneity of its composition and structure^55,58,120^. Nevertheless, the averaged cryo-ET subtomogram map of the isolated yeast NPC^17^ revealed three interrelated features of the central transporter (**Fig. 3a**, left): (i) a “moat” of low density surrounding the high-density region at the center of the central channel, (ii) “thin connections” of higher density that radially traverse the moat, and (iii) a central high-density region connected to the NPC scaffold *via* the thin connections. No model has yet reproduced these three features. To compare our integrative model and the EM maps of the central transporter, we averaged the densities of the FG repeats and NTRs in the ensemble of transport trajectories (**Fig. 3b**); in addition, the computed density, which was not originally symmetrically averaged, was averaged using C_8_ symmetry, as in the experimental maps. Strikingly, the resulting computed density map recapitulates the moat, thin connections, and the high-density region near the central axis in the cryo-ET map (**Fig. 3a,b**, insets), even though these features were not included in modeling.

As further validation, we also compared our simulated map with a new *in situ* cryo-ET map (**Fig. 3a**, right**; Extended Data Fig. 3)**. While the resolution of the *in situ* cryo-ET map is lower than that of the *in vitro* cryo-ET map (approximately 44 Å *vs* 28 Å), it is also consistent with the moat, thin connections, and central high-density features in our integrative model. Thin connections surrounded by a low-density moat are also clearly evident in other recent *in situ* cryo-ET and single particle isolated NPC maps of the central transporter^56,121^. These central transporter similarities among the *in vitro* cryo-ET map, *in situ* cryo-ET map, and map computed from the integrative model further increase our confidence in the integrative model applicability to functional NPCs.

## The thin connections are formed due to entropic confinement of FG repeats near the scaffold walls

With the four validations above in hand, we proceeded with an interpretation of the model. We started by interpreting the cryo-ET maps with the aid of the ensemble of transport trajectories. Naively, the low-density moat and thin connections could be interpreted as a peripheral cavity sectored by stable ultrastructures. However, neither the moat nor thin connections appear in any one simulation snapshot (**Fig. 3c**), only in the density map for the entire ensemble of transport trajectories (**Fig. 3b**). Thus, our integrative model indicates that the moat and thin connections in the cryo-ET maps should be interpreted respectively as low- and high-occupancy regions in a broadly distributed ensemble of disordered FG repeats, resulting from C_8_ averaging of thousands of NPC particles in cryo-ET map construction. As for the central high-density region, the integrative model suggests that it consists of both FG repeats and NTRs plus cargo molecules; the density of FG repeats is higher at the periphery than near the central axis, while the opposite is the case for the NTRs (**Fig. 3b**, bottom).

Because the moat and thin connections arise in the integrative map from the interactions between the model components of the transport system over time, analyzing the simulated trajectories reveals how they form. We hypothesized that the high-density thin connections emerge due to entropic confinement of the flexible FG repeats away from the NPC scaffold^28,29,56^. A theoretical underpinning for this hypothesis is provided by a smaller number of configurations of a flexible polymer close to its anchor point compared to a non-confined state closer to the center of the transporter (**Fig. 3d**). To test this hypothesis in a simpler setting, we simulated flexible polymers anchored to the walls of a cylinder, similarly to our previous miniature model of the NPC^20^. The simulations resulted in the C_8_ floret pattern seen in the actual central transporter, with low-density regions close to the FG repeat anchor points (**Fig. 3e**). The C_8_ floret pattern is also visible in previously published simulations in which FG repeats were represented as disordered polypeptides anchored to a fixed cylinder, although the pattern was not discussed^2,122^. Thus, entropic confinement is indeed a plausible rationalization for how the thin connections arise from the interactions among the system components, validating the predicted model morphology beyond its mere similarity to the cryo-ET maps.

## The densities of actively and passively diffusing molecules at the central transporter are distinct but overlapping

The integrative model indicates that the NTR:cargo density is concentrated some 5-15 nm away from the central axis of the NPC, while the density of the FG repeats is overlapping but it is shifted further away from the central axis by some additional 5-10 nm (**Fig. 3b**), with two high-density “lobes” along the channel axis forming at the cytoplasmic and nuclear sides of the central transporter, respectively (**Fig. 3g**, red). The lobes are consistent with high-density regions in the *in vitro* cryo-ET map (**Fig. 3f**, gray). We thus interpret the high-density lobes in the cryo-ET map to result from the high-occupancy of the NTR:cargo complexes, in agreement with the large fraction of the NTR and cargo molecules found in a mass spectrometry analysis of NPCs^17^. The distribution of both NTR:cargo complexes and FG repeats is nonetheless broad, with lower densities observed well into the cytoplasm and nucleus (**Fig. 3g**), also matching the cryo-ET map^17^.

The peak density of passively diffusing molecules is shifted even further toward the central axis compared with that of the NTR:cargo complexes (**Fig. 3h,i**), in agreement with imaging of the radial preference for passive diffusion^123,124^. As expected, the absolute density of the NTR:cargo complexes is much higher than that of passively diffusing molecules (**Fig. 3i, top**). In relative terms, the NTR:cargo complex density is peaking midway between the central axis and scaffold walls (**Fig. 3i**). The partial overlap between the NTR:cargo and FG densities (**Fig. 3g,i**) arises from a balance between their attractive interactions and repulsive excluded volume effect. In other words, smaller molecules tend toward the pore’s central axis while NTR:cargo complexes partially overlap with the FG repeats, driven by a balance between FG-binding interactions and excluded volume effect. Given the anchoring of the FG repeats to the NPC scaffold, such partial overlap is also consistent with the peak density of cargo:NTR complexes shifting further away from the central axis by some 10 nm in the dilated state in recent super-resolution 3D imaging of translocating cargo molecules^27^.

## The model reproduces size-dependent fluxes for passive diffusion

A useful model of transport must recapitulate one of the NPC’s most basic functions, that of forming a passive permeability barrier. We therefore validated predictions of flux for passively diffusing molecules of different sizes against experiment. In the model, the rate of diffusion is measured along the reaction coordinate corresponding to the NPC central axis, bounded by the two sides of the central channel. This rate corresponds to the NPC permeability (*p*), an intrinsic property of the NPC for a given diffusing molecule^18,20,77^; *p* is estimated from simulation by counting the number of diffusing molecules that pass through the NPC reaching the opposite channel opening per unit of time per unit of the difference in the concentrations of the diffusing molecules in the cytoplasmic and nuclear compartments (Δ*c*); the flux through a single NPC is *p* Δ*c*, according to the Fick’s first law. The permeability of the NPC in the model decreased as a cubic root of the molecular weight of the diffusing molecule (**Fig. 4a**) (or, equivalently, as an exponent of its molecular radius with a decay constant of -0.37 Å^-1^) (**Extended data Fig. 4a**). This result is in agreement with both our *in vivo* FRAP measurements of transport rates and previous simulations of a minimal pore model consisting of a cylindrical pore lined with generic flexible polymers (**Fig. 4a**)^20^, reminiscent of artificial NPC mimics previously used to study nucleocytoplasmic transport^60,83,103^. It also closely fits experimental results for passively diffusing molecules of varying sizes, shapes, and domain compositions^19,20^. These agreements indicate that a flexible polymer anchored to the scaffold walls is sufficient to a first order to reproduce the observed size-dependent barrier to passive diffusion of the NPC^28,29^, without additional considerations. Both the model and *in vivo* experimental data indicate that passive diffusion diminishes continuously with increasing molecular weight, in agreement with our previous studies of passive diffusion^20^, which revealed no well-defined size threshold (*eg*, 30-60 kDa),^20^.

We also found that the size-dependence of passive diffusion in yeast is conserved in other organisms and cell types, after controlling for differences in cell volumes (**Fig. 4a**; **Extended Data Fig. 4b; Supplementary Results**). Thus, our NPC model can accurately quantify passive NPC transport rates across multiple species.

## An exponential relationship between passive diffusion fluxes and densities

We assessed whether the changes in the flux of passively diffusing molecules as a function of their size correspond to changes in their density inside the NPC. The model shows that the density of passively diffusing molecules is indeed lower inside than outside the NPC, with the decrease being more pronounced for larger, low-flux molecules (**Fig. 4b-e**). The dependence of the flux on the density in our model is exponential, similar to the Eyring-Polanyi equation embodying the transition state theory (TST)^125,126^, as follows. In TST, an elementary chemical reaction proceeds from reactants to products through a transition state. The Eyring-Polanyi equation predicts that the reaction rate decreases exponentially as a function of the Gibbs free energy difference between the ground and transition states (free energy barrier). In the NPC, the reaction coordinate is the central axis of the channel, the reactants are molecules on one side of the NPC, the products are the same molecules on the other side of the NPC, the transition state corresponds to populating the central transporter of the NPC, and the transport rate decreases exponentially as a function of the free energy difference between one side of the NPC and the central transporter.

Our spatiotemporal model allows us to assess whether the Eyring-Polanyi equation holds for the unabridged NPC geometry, FG Nup anchoring patterns, and FG Nup composition. The free energy barrier was estimated *via* the Boltzmann inversion of the observed steady-state density of diffusing molecules. The resulting 2D and 1D free energy maps show that the free energy barrier for passive diffusion increases linearly with molecular radius, by 1 *k*_B_*T* for every 0.5 nm in radius (**Extended Data Fig. 4c**)^20,123^; and sub-linearly with molecular weight (**Fig. 4b-e; Extended Data Fig. 4c, right panel**), by 1 *k*_B_*T* for every 34 kDa in molecular weight of less than 150 kDa, using a first-order approximation (**Extended Data Fig. 4c, right panel**). In a remarkable similarity to TST, permeability to passive diffusion indeed depends exponentially on the free energy barrier of transport (**Fig. 4f**), fitting the Eyring-Polanyi equation with *R*^2^=0.999. This empirical result is consistent with the slow exponential decay of NPC permeability as a function of the radius of the diffusing molecule (**Extended Data Fig. 2a**) and its cubic decay as a function of the molecular weight of the diffusing molecule (**Fig. 4a**), observed in both simulation and experiment. Thus, the kinetics of passive diffusion across the NPC may be modeled similarly to that of an elementary chemical reaction. Finally, the direct correspondence between the free energy barrier and the kinetics of passive transport indicates that the steady-state densities of passively diffusing molecules highlight spatial pathways for passive diffusion.

## The density of transporting NTR:cargo complexes decreases slowly with molecular weight, in contrast to passive diffusion

We next investigated how NTRs mediate rapid facilitated diffusion (*ie*, selective transport) of cargo molecules, including large cargo molecules that cannot diffuse through the NPC passively. We began by simulating NTRs with a fixed valency of 4 interaction sites for FG repeats on each NTR, corresponding to the relatively weak (sub-millimolar) affinity of NTRs, such as the NTF2 dimer with four such sites (**Fig. 1f**). We assessed how the densities of the NTR:cargo complexes change as their molecular weight increases from 14 kDa to 133 kDa (**Fig. 5**).

Compared with passively diffusing molecules, the densities of the NTR:cargo complexes inside the central channel decreased far more slowly as a function of their molecular weight (**Fig. 5a,b**). Moreover, despite the weak (millimolar) affinities of individual interaction sites (**Fig. 1c,d,e**), the concentrations of the NTR:cargo complexes were much higher than those of passively diffusing molecules of the same molecular weight (**Fig. 5b**). The high densities of the NTR:cargo complexes are explained by the avidity (higher affinity) of NTRs with multiple interaction sites (**Fig. 1d,e**; **Fig. 2b**)^127^. These densities are consistent with the bright NPC-associated staining of many NTR:cargo complexes seen in actual cells^75^, the relatively large collective mass of NTR:cargo complexes (>26 MDa) in the yeast NPC^17^, and the 20-50 mM concentration of individual FG motifs in the central channel^17,35,47^, which exceeds the measured dissociation constant for NTRs (**Fig. 1e**)^102^.

## The high density of large NTR:cargo complexes reflects rapid transport fluxes

Compared to passively diffusing molecules, the higher densities of NTR:cargo complexes in the central transporter imply that the free energy barrier for their facilitated diffusion is lower (**Fig. 5c,d**). In fact, the free energy for facilitated diffusion has a local minimum in the central channel on either side of the maximum at its equator (*Z*=0 nm). This free energy barrier was only modestly higher for large than for small NTR:cargo complexes (**Fig. 5d**). As expected, the much higher densities and lower free energy barriers of the NTR:cargo complexes compared to passively diffusing molecules were indeed manifested in higher permeability for facilitated than passive diffusion. Nonetheless, NTRs with larger cargo molecules were distributed farther from the NPC “equator” at *Z*=0 (**Fig. 3e**) and closer to the central axis of the NPC compared to NTRs with smaller cargo molecules (**Fig. 3f**), again in agreement with previous observations^123^. This size-dependent difference is rationalized by the higher density of FG repeats away from the central axis, which exerts a stronger repulsive force on larger cargo molecules, pushing them toward the less dense regions.

## The model reproduces the speed and weak size-dependence of facilitated diffusion

While passive diffusion rates decrease rapidly with molecular weight (∼MW*^-^*^3^), facilitated diffusion of NTR:cargo complexes has a significantly weaker dependence on molecular weight (∼MW*^-^*^2/3^) than passively diffusing macromolecules (**Fig. 5g**), in agreement with *in vivo* measurements^71^. NPC selectivity, defined as permeability to facilitated diffusion at a given molecular weight divided by permeability to passive diffusion at the same molecular weight, increases from 30-fold at 14 kDa to 1000-fold at 133 kDa. As a result, the facilitated diffusion rate of a 133 kDa NTR:cargo complex is predicted to be 50% faster than that of a 14 kDa macromolecule, whereas its passive diffusion rate is 1000-fold slower, in agreement with experiment^20^. Moreover, the model reproduces experimental results showing that even small cargo molecules (∼20 kDa) are transported significantly faster *via* facilitated diffusion than passive diffusion^18^, thus rationalizing why even relatively small cargo molecules may require an NTR for their import into the nucleus (*eg*, Ran requires NTF2, while histones and ribosomal proteins require cognate karyopherins)^18,112^. In conclusion, our spatiotemporal model recapitulates the speed and weak size-dependence of facilitated transport^18^.

## Slide-and-exchange increases the rate of facilitated diffusion by an order of magnitude

We used our model to investigate how interactions between NTR:cargo complexes and FG repeats impact the kinetics of facilitated diffusion, all else being equal. We have already established that slide-and-exchange of FG repeats on NTF2^43^ and karyopherins (**Fig. 2a**) results in a rapid molecular exchange (**Fig. 2b**), in agreement with experiment^40,41,43^. We hypothesized that slide-and-exchange contributes to a reduction of flow resistance (and thus, to an increase in the flow) of the NTR:cargo complexes through the NPC. Flow resistance can be quantified by dynamic viscosity, which is inversely proportional to the diffusion coefficient according to the Einstein-Stokes relation^128,129^. With sliding, the apparent diffusion coefficient of the NTR:cargo complexes within the central channel is indeed only 32-43% slower than in the cytoplasm, compared to a 44-66% reduction without sliding, in agreement with previous experimental estimates of NTF2’s diffusion coefficient within the central transporter^18^ (**Fig. 5h**). Thus, slide-and-exchange helps compensate for added molecular crowding in the NPC, by limiting the change in dynamic viscosity for diffusing NTR:cargo complexes when they enter or exit the central transporter. Correspondingly, the permeability of the NPC to NTR:cargo complexes was 1-2 orders of magnitude higher with sliding of NTR:cargo complexes than without sliding (**Fig. 5i**, red *vs* pink), everything else being equal, including NTR:FG repeat avidities (**Fig. 2b**, top-left). Furthermore, sliding contributed to weakening size-dependence by disproportionately increasing the permeability of the NPC to larger NTR:cargo complexes (**Fig. 5i**, pink *vs* blue). Thus, slide-and-exchange is likely to contribute significantly to both the speed and selectivity of nucleocytoplasmic transport.

## The number of FG interaction sites on NTRs modulates facilitated diffusion rates

We next characterize how the multivalency of NTR:cargo complexes influences facilitated diffusion kinetics, revealing a tradeoff between valency and cargo size. Various NTRs, including larger ones such as Kap95^130^ and CRM1^52^ as well as smaller ones such as NTF2^51^, have multiple interaction sites for FG repeats^130,131^. In addition, some cargo molecules interact with multiple NTRs simultaneously, resulting in NTR:cargo complexes of high valency^53^. This valency impacts the avidity and exchange rates^132–134^ between FG repeats and NTRs^102,135^, but its effect on the kinetics of facilitated diffusion has not yet been comprehensively quantified, although several studies have indicated that adding even a small number of NTRs to a cargo molecule can significantly increase the transport rate^74,106^.

We compared trajectories of NTR:cargo complexes with identical molecular weights, but with the number of FG interaction sites varying from zero to eight (**Fig. 6**). As expected, their central transporter densities increase across the entire range; the density of complexes with even a single interaction site is already six-fold higher than that of complexes with no interaction sites (*ie*, equivalent to passive diffusion). For four sites or more, the central channel contains more NTRs than the cytoplasmic and nuclear peripheries (**Fig. 6a,b**); correspondingly, the free energy within the central transporter is lower than in the NPC periphery (**Fig. 6c,d**).

The gradual decrease in the free energy barrier due to the addition of FG interaction sites on an NTR results in an exponential increase in the permeability of the NPC for facilitated diffusion (**Fig. 6e**). Even a single interaction site is sufficient for a modest but notable three-fold increase in the NPC permeability (**Fig. 6e**), while six interaction sites increase the permeability by almost three orders of magnitude. However, as the number of sites rises further from six to eight, permeability begins to decrease, reflecting a tradeoff between the decrease in the free energy barrier of passive diffusion and entrapment caused by overly strong avidity within the central transporter, resulting in decreased diffusion. In conclusion, our reductionist model quantitatively reproduces and thus explains the changes in the rate of facilitated diffusion as a function of the number of FG interaction sites on the NTR.

## A tradeoff between the NTR valency and cargo size results in efficient facilitated diffusion of very large cargoes

We performed simulations in which we varied both the number of FG binding sites on the NTR surface from 0 to 16 and the size of the NTR:cargo complex from 16 kDa to 1.4 MDa. In total, 70 combinations of cargo size and valency were simulated for 80 μs each. Each simulation was repeated 100 times, for a total of 7,000 trajectories covering 500 seconds of facilitated diffusion.

For smaller NTR:cargo complexes (<400 kDa), the permeability increases exponentially as a function of NTR valency, from 0 to 6 FG interaction sites (**Fig. 7a**). For larger complexes (>400 kDa), significant facilitated diffusion is not observed for 0 to 4 sites, while it does occur for 6 or more interaction sites. However, permeabilities of the smaller complexes begin to decrease markedly for more than 6 sites, dropping to zero for 12 or more sites. In contrast, for larger complexes, the permeability decreases only slightly, or even increases, with additional interaction sites. As expected, for 0 interaction sites (*ie,* for passive diffusion), permeability sharply decreases with molecular weight, following the inverse cubic power law (above) (**Fig. 7b**). As the number of sites increases to 6, the relatively high permeability decreases more gradually with molecular weight. For more than 6 sites, the diffusion does not depend on the cargo size, indicating that high valency benefits larger cargos more than smaller cargos. As a result, the significant decrease in the permeability due to molecular weight (**Fig. 4a**) can be counteracted by adding a relatively small number of interaction sites (**Fig. 7a,b**). This observation is in agreement with experiment^74,106^ and resolves the apparent contradiction between the ability of the NPC to translocate larger MDa cargo molecules while significantly slowing the passive diffusion of small <50 kDa macromolecules.

Our model indicates that several particles the size of pre-ribosomal subunits can occupy the central transporter simultaneously (**Fig. 7c**), despite the voids that they introduce within the thicket of FG repeats (**Fig. 7d**). This result is validated by the recent cryo-EM structures of native pre-60S ribosomal particles within the NPC, showing multiple particles occupying the central transporter of the NPC with a relatively small number of accompanying NTRs^104^. Our model predicts that the multivalent interactions of the NTRs compensate for the entropic restriction of the FG repeats by these large particles. An additional strong validation of our model is provided by a comparison of the radial distribution of large particles in our simulations (diameter of 15 nm) and the recently measured radial distribution of the pre-60S particles (diameter of 21 nm) in the central channel^104^ (**Fig. 7e**). Our model correctly predicts that transport pathways are broadly distributed with a peak approximately 10 nm from the central axis of the NPC.

## Active Ran-dependent transport of large cargos is rationalized by coupling the spatiotemporal NPC transport model with a kinetic cell model

We next assessed our model of NTR-mediated facilitated diffusion of large cargoes in the context of experimental measurements in cells^74,10674^. In cells, NPC transport is energy-dependent and directional due to the concentration gradient of the RanGTP protein across the NE (active transport)^136,137^. This gradient provides the energy for directional transport through asymmetric dissociation of imported and exported complexes on the nuclear and cytoplasmic sides of the NPC, respectively^2,138,139^. In contrast, in our spatiotemporal model, transport is neither energy-dependent nor directional, due to the absence of RanGTP. Therefore, the experiments quantify active Ran-dependent transport rates of NTR:cargo complexes under non-equilibrium conditions, whereas our spatiotemporal model quantifies non-active facilitated diffusion rates under equilibrium conditions.

Both the experimental measurements and simulations (**Figs. 6 and 7**) indicate that a relatively small number of nuclear localization sequences (NLSs) are sufficient to allow translocation of very large (> MDa) cargo molecules. This agreement is consistent with the finding that RanGTP hydrolysis is not directly coupled to any molecular rearrangement of the NPC^2^; indeed, reversing the RanGTP gradient leads to a reversal of transport direction^137^. Thus, Ran is not required for rationalizing how a large cargo molecule permeates the NPC.

However, our simulations (**Figs. 6e and 7a,b**) initially did not reproduce the monotonic increase in the transport rate with additional FG interaction sites^74^ (**Fig. 8a**). We investigated this discrepancy by considering Ran-dependent transport in an entire cell and Ran-independent transport through a single NPC in tandem (**Fig. 8b**). To do so, we used our recent Bayesian metamodeling approach^99^. In this approach, a complex biological system is represented by a set of coupled models of different aspects and/or parts of the system (*eg*, trajectories of protein molecules and a network of protein-protein interactions). The models are then updated with respect to each other to obtain a more accurate, precise, complete, and explanatory depiction of the system (a metamodel). Here, a metamodel of Ran-dependent active transport in the cell couples two models. The first one is our spatiotemporal model of Ran-independent facilitated diffusion through a single NPC. The second model is our recent kinetic model of Ran-dependent transport in the cell^71^. The kinetic model aims to capture the broadly accepted features of Ran-dependent transport^2,46,69,70,140^ using a system of ordinary differential equations whose rate constants were fit to empirical measurements^71^. It describes facilitated diffusion of NTR:cargo complexes, passive diffusion of unbound cargo molecules, the Ran-dependent release of cargo in the nucleus, and the energy-dependent cycle that replenishes the concentration gradient of RanGTP across the NE. The parameters include rate constants for docking of the NTR:cargo complexes to the NPC and the facilitated diffusion rate of the NTR:cargo complexes between the cytoplasmic and nuclear sides of a single NPC (**Fig. 8a**, third column). These kinetic model parameters are coupled with the corresponding spatiotemporal model parameters, allowing us to refine their values using the metamodeling approach^99^. The result is a metamodel of nucleocytoplasmic transport at both single NPC and cellular levels.

Remarkably, the metamodel (**Fig. 8b-d**) resolves the above-mentioned discrepancy between the monotonic increase in measured Ran-dependent transport rates^74^ (**Fig. 8a**) and the saturation of simulated Ran-independent permeabilities as a function of valency (**Figs. 6e and 7a,b**). The metamodel correctly predicts that the initial flux and nuclear concentration of cargo increase significantly and monotonically when the docking affinity of NTR:cargo complexes to the FG repeats increases, even if the NPC permeability remains constant (**Fig. 8c; Extended Data Fig. 5a**). To verify that this effect is due to the RanGTP concentration gradient, we used a negative control simulation in which RanGTP could not release NTR:cargo complexes. As expected, increasing the docking affinity beyond a threshold indeed results in a decrease in the rate of equilibration across the NE (**Fig. 8d; Extended Data Fig. 5b**). Moreover, given that the NPC-bound NTRs could no longer dissociate from the cargo *via* binding to RanGTP, NTR:cargo complexes accumulated within the NPC (**Extended Data Fig. 5b**, bottom row), in agreement with experimental disruption of RanGTP-mediated cargo release^75^. In conclusion, both the spatiotemporal model of a single NPC and the kinetic model of the cell are validated by the experimental observations (**Fig. 8a** *vs* **Fig. 8c**). Thus, they provide a rationalization of how higher avidity for the NPC increases transport rates under non-equilibrium (Ran-dependent) conditions, but not under equilibrium conditions (**Fig. 8c** *vs* **Fig. 8d**).

## FG:FG interactions are not obligatory for rapid and selective transport

We also characterized how perturbations of the composition and geometry of the NPC influence the size-dependence and rates of facilitated and passive diffusion, and consequently, transport selectivity. We began by assessing the impact of interactions between pairs of FG repeats, or FG:FG interactions. Some phenomenological models of transport suggest that transport selectivity arises mostly from the interactions between pairs of FG motifs, based on experiments with saturated FG repeat hydrogels that by definition consist of highly cohesive FG repeats^35^ as well as more recent experiments with phase-separated FG repeats that also require significant FG repeat cohesion^84,85^. In contrast, other phenomenological models suggest instead that selectivity arises mostly from the entropic cost of restricting the FG repeat conformations^28–30^.

To address whether or not FG:FG repeat interactions are obligatory for fast and selective transport, we repeated coarse-grained Brownian dynamics simulations with either reduced or increased interactions between all types of FG motifs. When FG:FG interactions are eliminated entirely, the permeability of the NPC for both facilitated and passive diffusion increases in comparison to the wildtype NPC (**Fig. 9a**, right panel, yellow *vs* green). However, even without any FG:FG interactions, passive diffusion remains size-dependent. As a result, the permeability for facilitated diffusion remains orders of magnitude higher than for passive diffusion (**Fig. 9a-b**, yellow), albeit the transport selectivity of the wildtype is even stronger (**Fig. 9b**, green). Thus, cohesive interactions between pairs of FG motifs are not obligatory for rapid and selective transport in our model.

## Weak FG:FG repeat interactions fine tune the speed and selectivity of transport

The NPC permeability decreases with an increase of the interactions between the FG motifs (**Fig. 9a**). However, beyond weak FG:FG interactions, it decreases more precipitously for facilitated than for passive diffusion (**Fig. 9a**, green to cyan), reducing the selectivity relative to the native NPC (**Fig. 9b**, cyan). Thus, according to our model, overly strong FG:FG interactions may reduce transport selectivity altogether. In contrast, a low level of transient FG:FG interactions in the native NPC leading to short-lived and “fuzzy” FG:FG interactions, on the order of picosecond to nanosecond, is consistent with the empirically-observed fast exchange of FG repeats and NTRs^40,41,43^ (**Figs. 2b and 5i**). Thus, weak FG:FG interactions may improve selectivity, without being obligatory for it. Such interactions are consistent with the radius of gyration of the FG repeats (mainly GLFG repeats^32^) of some FG Nups, which in our model are fitted to SAXS, SANS, and FPLC experimental data and atomistic molecular dynamic simulations (**Extended Data Fig. 2a**). The weak FG:FG interactions are also consistent with the previously discussed increase of the end-to-end distance of the GLFG-rich Nup100 *in situ* relative to buffer in both FRET experiments^73^ and our model (**Extended Data Fig. 2b**), implying that FG:FG interactions are sufficiently weak to be overcome in the crowded environment of the central channel. Weak FG:FG interactions are also consistent with the suggested mechanism allowing the NPC to avoid clogging^49^, while still accounting for the reported higher-density regions in central channel^60^.

To test our predictions regarding the influence of FG:FG interactions on transport, we compared them to previous experiments and simulations of transport in artificial nanopores grafted with FG repeat domains of either wildtype Nsp1 or Nsp1-S^103^; Nsp1-S is identical to Nsp1 except for the hydrophobic FG residues that are mutated to the hydrophilic SG residues. These mutations are expected to have two major consequences: First, by reducing hydrophobicity, they reduce the interactions among the FG repeats, in particular for the relatively compact N-terminal regions of Nsp1^32^. Second, by eliminating the FG motifs, they disrupt their FG-mediated interactions with NTRs^103^. The simulation of the wildtype Nsp1 or Nsp1-S^103^ nanopores relied on the pore geometry and estimated grafting density from the original study^103^, while retaining the representation and parameters for the FG repeats, passively diffusing molecules, and NTR:cargo complexes of our current simulations. The resulting model reproduced the increase in passive permeability of the Nsp1-S *vs* wildtype Nsp1 nanopore^103^ (**Extended Data Fig. 6**, blue *vs* red). Our model also predicts that passive diffusion remains size-dependent even for the Nsp1-S nanopore, with the permeability increasing by an order of magnitude as the molecular weight is doubled.

## FG:FG interactions and NTRs contribute to robustness of transport

In addition to tuning the speed and selectivity, FG:FG interactions may contribute to the robustness of transport by, for example, limiting undesired leakage through the NPC. NTRs were also suggested to contribute to the robustness of transport. For instance, the NTR-centric model of transport (originally, Kap-centric)^76,141,142^ posits that the large population of NTRs in the central transporter^17,75^ directly contributes to size-dependence of passive diffusion. In agreement, in our model, the rise in permeability to passive diffusion in Nsp1-S mutant nanopore relative to the wildtype nanopore is reversed by re-introduction of NTR binding (but not FG:FG interactions) to Nsp1-S (**Extended Data Fig. 6,** green). This reversal indicates that both FG:FG interactions and the NTRs in the central channel confer robustness on the size-dependent free energy barrier.

## The model reproduces robustness of transport to FG Nup deletions

Additional mechanisms may contribute to the robustness of rapid and selective transport. Such mechanisms are further exemplified by the viability of mutant strains with FG Nup deletions^143^, which manifest relatively modest changes in passive and facilitated diffusion^20^. To study this robustness, we simulated transport through the NPC without Nsp1, eliminating 32% of the FG repeat mass. The simulations show that these NPC mutants retain both size-dependence and selectivity of transport (**Fig. 9c**). This prediction is validated directly by *in vivo* experimental measurements of passive and facilitated diffusion for the nsp1^ΔFG^ mutant strain^20^. It is also consistent with the size-dependence and selectivity of transport through NPC mimics with a single type of FG repeat^60,83,103^. The robustness of transport to changes in the FG Nup composition may be key to the NPC ability to evolve gradually and adapt to diverse needs of different types of eukaryotic cells^144^.

A characterization of the different FG repeat flavors and the resulting heterogeneity of their potential interactions and contributions to the nucleocytoplasmic transport^145^ remains beyond the scope of the current work. For example, different FG repeat flavors have varying *in vitro* affinities for different FG repeats and NTRs, which may influence both the type and amount of NTRs recruited to the NPC and so affect the overall permeability of the NPC as well as the efficiency of particular transport pathways^3^.

## NPC dilation increases NPC permeability

We assessed whether the robustness of transport to variations in FG repeat identities, copy numbers, anchoring positions, and FG:FG interactions extends to variation of the NPC diameter. While early evidence for this variation^146,147^ has been originally dismissed as spurious and without functional implications^147^, additional evidence for the functional importance of the variable NPC diameter has been mounting^56,71,78–80,148^. Thus, it was hypothesized that the dilation of the NPC plays a functional role in adaptations to internal and external stimuli, including mechanical stress, hyperosmotic shock, energy depletion^71,80,81^, and development^149^. Interestingly, *in vivo* measurements of both passive and facilitated diffusion decreased markedly under cellular conditions coinciding with a constricted NPC^80,82^, although implications for the structure-function relationship of the NPC remain uncertain due to the multitude of other concomitant phenomena^80^. It has been demonstrated using both experiment and simulation that artificial nanopores with increased diameter lead to faster translocation rates of Kap95 (facilitated diffusion) as well as BSA (passive diffusion), accompanied by an up to 8-fold increase in selectivity^60,83^.

To assess the dependence of transport on the diameter of the central channel, we relied on our minimal pore model with only 32 chains of FG repeats^20^, in which the parameters describing the FG Nups and their interactions with the NTR:cargo complexes were updated to the current values. The pore radius was varied from 13 to 25 nm in 1.5 nm increments. For each pore radius, 100-200 independent trajectories of 20 μs each were computed for both passive and facilitated diffusion, while keeping all other model parameters fixed. The use of the minimal pore model to study the dependence of the NPC permeability on its diameter is justified because the simplified pore model approximately reproduces both transport selectivity and rates of the more detailed model (**Fig. 9d-e**; *cf*, **Figs. 4a and 5g**).

In the simplified pore model, the permeability to facilitated diffusion increases incrementally but substantially with the NPC radius, growing by almost 10-fold as the radius increases from 13 to 25 nm (**Fig. 9d**). Passive diffusion increases even more significantly. Its size dependence is maintained, but passive permeability increases more significantly for larger than smaller molecules, decreasing the downward slope of the permeability *vs* molecular weight curve as the pore radius increases (**Fig. 9e**). We conclude that the dilation of NPCs *in vivo* may indeed explain the increase in the NPC permeability under some conditions, such as energy depletion, osmotic shock, and mechanical stress^71,80,81^.

## Discussion: the transport system has evolved for robustness

After validating and dissecting our model, we are now in a position to discuss the mechanism of transport and the design features that lead to its speed and selectivity as well as their robustness to perturbations in the transport system and its environmental conditions, such as the properties of transported cargo:NTR complexes, the NPC geometry, and the NPC composition.

In our model, nucleocytoplasmic transport arises only from the composition of the system and interactions between its components; thus, the model avoids making circular phenomenological assumptions about key emergent properties of the transport. Nevertheless, the model reproduces these properties after its parameters are fit to available experimental data and theoretical information. As a result, we can explain how the emergent properties of transport result from ten key features of molecular interactions between the system components (**Fig. 10**). Next, we discuss each one of these key design features in turn.

**Fig. 10.**
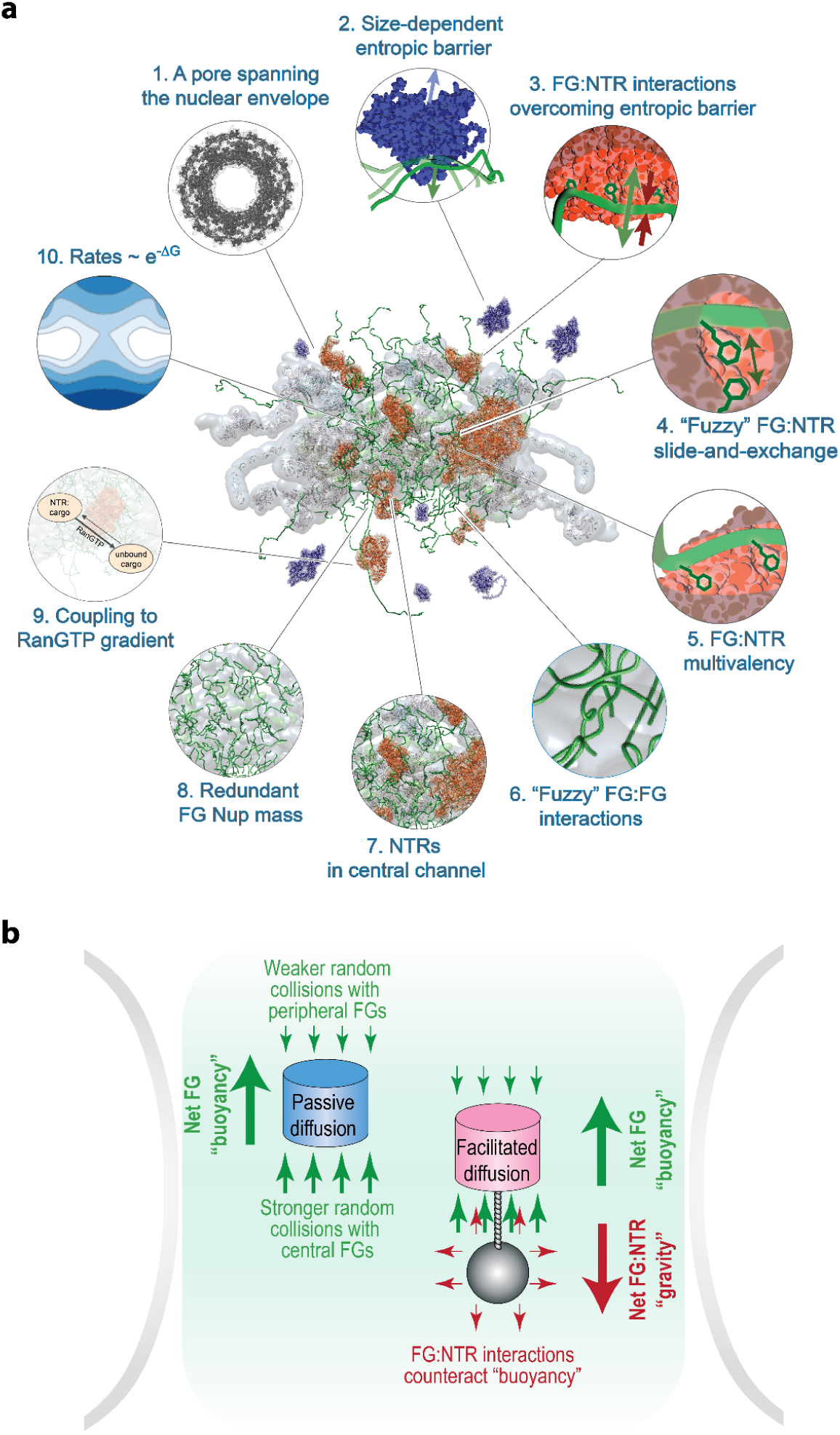
Key design features of the NPC. **a**, A snapshot of macromolecules diffusing through the pore is shown in the center. The small panels encircling the snapshot indicate the molecular design features of the transport system. Panels 1-3 indicate the key features of the virtual gating mechanism that confer transport speed and selectivity^28,29^. The remaining panels 4-10 indicate features that fine tune the speed and selectivity as well as provide their robustness. **b**, The parallel between the molecular origins of nucleocytoplasmic transport and buoyancy.

The first three features correspond to the virtual gating model (*ie*, Brownian affinity gating model) (**Fig. 10a**)^28,29,68^: a pore spanning the NE (**Fig. 1a**), flexible FG repeats forming a size-dependent entropic barrier to passive diffusion through the pore (**Fig. 4**), and transient interactions between the flexible FG repeats and NTR:cargo complexes to reduce the free energy barrier of facilitated diffusion (**Figs. 5-7**). In contrast to the phenomenological virtual gating model^3^, the current model does provide both a reductionist explanation of transport and its quantitative molecular depiction. As a result, we can suggest another perspective on virtual gating, by noting its parallels with buoyancy ^150^ (**Fig. 10b)**. In this parallel, FG Nups generate a net repulsive force that scales with the volume of displaced FG repeats, driving passively-diffusing molecules outward in a size-dependent manner. Microscopically, this net repulsive force is explained by more frequent random collisions with central than peripheral FG repeats, analogous to the pressure difference between the bottom and top sides of submerged objects that drives buoyancy. NTRs counterbalance this net force through their dynamic interactions with the FG repeats, selectively enabling them to “sink” into the FG milieu, analogous to how weight counters buoyancy.

In addition to the three design features explaining quantitatively how virtual gating results from molecular interactions, our model reveals additional seven features of transport design (**Fig. 10a**) that work in unison to confer robustness as follows.

The fourth design feature is the “fuzzy” interactions between FG repeats and the NTR:cargo complexes as described by the anisotropic slide-and-exchange model^43,92^; “fuzziness” denotes a highly-dynamic (transient) interacting state with an ensemble of rapidly interconverting conformers^92,151^ (**Figs. 2 and 5**). This design feature increases the exchange rate of interacting FG repeats and NTRs without reducing the avidity^40,41^ (**Fig. 2**), thereby enhancing both the selectivity and rate of facilitated diffusion by over an order of magnitude compared to less “fuzzy” isotropic interactions (**Fig. 5i**, pink). A “fuzzy” interaction between FG repeats and NTR:cargo complexes may also contribute to the resilience of the native NPC to clogging by slow exchanging NTR:cargo complexes.

The fifth design feature is the FG:NTR multivalency, with one NTR molecule simultaneously interacting with one or more FG motifs from one or more FG Nups (**Fig. 1c-f**; **Fig. 7a-b**). Our model shows how this feature enhances both the selectivity and rate of facilitated diffusion for multivalent NTR:cargo complexes by several orders of magnitude compared with monovalent NTR:cargo complexes (**Fig. 7a-b**), due to multiple relatively weak interaction sites increasing the avidity without compromising the exchange rate (**Fig. 2**). The ability to increase the avidity without compromising molecular exchange, conferred by both “fuzziness” and multivalency of the FG repeat:NTR interactions, is critical to the robust transport of a wide range of molecular cargos, including large pre-ribosomal subunits (**Fig. 7**) and capsid-like particles (**Fig. 8**). As an aside, even NTR:cargo interactions can be multivalent (not studied here), in the sense that one cargo molecule can bind multiple NTRs ^74,104^.

The sixth design feature is potential “fuzzy” interactions between pairs of FG repeats (**Fig. 9a-b; Extended Data Fig. 6**).^151^ This feature impacts selectivity by reducing the rate of passive diffusion more strongly than for facilitated diffusion (**Fig. 9a-b**, green *vs* yellow). Nonetheless, overly strong interactions between pairs of FG repeats are detrimental to selectivity (**Fig. 9b**). Indeed, the model of the wildtype NPC is in agreement with experimental data on size-dependent passive diffusion (**Fig. 4a; Extended Data Fig. 3a**) without recourse to or dependence on stronger interactions assumed in a model of a saturated hydrogel^35^; in fact, our model predicts that transport would maintain substantial selectivity even without any FG:FG interactions (**Fig. 9b**, green).

The seventh design feature is the presence of NTRs in the central channel^76,141,142^ (**Extended Data Fig. 6**, green). This feature greatly enhances selectivity by reducing the rates of passive diffusion, contributing to the robustness of the size-dependent barrier.

The eighth design feature is redundant FG Nup mass (**Fig. 9c**). This feature enhances the robustness to perturbations in FG Nup composition, such as evolutionary deletions and mutations of FG Nups, because the transport system is already selective for a low copy number of FG Nups.

The ninth design feature is a consequential result of the well-established coupling of facilitated diffusion to the concentration gradient of RanGTP^152^ (**Fig. 8; Extended Data Fig. 5**). This feature enhances the transport rates for large NTR:cargo complexes even when the permeability saturates as a function of valency at the single NPC level (**Fig. 8**). Thus, it enhances the robustness of transport, as it prevents permeability from becoming a bottleneck for transport of very large cargo molecules, such as pre-ribosomal subunits and viral particles. We demonstrate this feature by the coupling of our spatiotemporal model of transport through a single NPC to a kinetic model of transport in the cell^71^, allowing us to quantitatively explain the empirical relationship between transport rates of large cargo and valency^74^. Such coupling, made possible *via* our recent Bayesian metamodeling framework^99,153^, can be used in future to investigate the suggested role of GTP availability in modulating nucleocytoplasmic transport^154^.

The tenth design feature is the exponential coupling of transport kinetics and thermodynamics in analogy to the TST^20,125,126^ and Arrhenius kinetics^155^ (**Fig. 4c-f**; **Fig. 5c-d**; **Fig. 6c-e; Extended Data Fig. 4c**); the TST predicts that reaction rates decrease exponentially with the magnitude of an energy barrier for passive diffusion (**Fig. 4f)**, but increase exponentially with a decrease in the energy barrier as a result of FG repeat:NTR interactions during facilitated diffusion (**Fig. 5-6**). This feature thus enhances transport selectivity, and helps rationalize the NPC’s ability to facilitate the transport of megadalton cargo molecules as large as pre-ribosomal subunits through the transient energetic contributions from only a few NTRs (**Fig. 7-8**), while inhibiting the passive diffusion of molecules as small as a few dozen kDa (**Fig. 4**).

In conclusion, the model reveals how key emergent properties of NPC-mediated transport, including rate, size-dependence of the rate, and selectivity, arise from the system components and their interactions, in a robust fashion with regard to environmental fluctuations and evolutionary pressures. Consideration of this robustness may allow us to rationally modulate and control the transport system and artificial mimics inspired by it.

## Relation to other types of biophysical systems

The NPC is a unique cellular apparatus that acts as the gateway to the nucleus. Its overall architecture and function have been remarkably conserved across all eukaryotes^56^ since it originated presumably in the last eukaryotic common ancestor 2.7 billion years ago^144^ (**Fig. 4a; Extended Data Fig. 4a-b**). The NPC must selectively transport cargoes of diverse chemistry, shape, and size. The design of the NPC has therefore evolved to perform these roles robustly with regard to the environmental noise and evolutionary perturbations. We next contrast the NPC with several other types of biophysical systems, including polymer brushes^30,73^, biomolecular condensates^156^, and saturated hydrogels^35^.

Polymer brushes are commonly defined as polymer chains densely tethered to another polymer chain or surface^30,73^. Similarly to polymer brushes, the end-to-end distance of the FG repeats in our model increases upon grafting in the central channel (**Extended Data Fig. 2b**), a phenomenon proposed as a possible basis for an entropically-driven selectivity barrier^30,73^ involved in virtual gating, as discussed above. It remains a major factor in explaining the selective transport mechanism in our model.

Biomolecular condensates in cells are loosely defined as micron-scale compartments that lack surrounding membranes, but concentrate biomolecules including proteins and nucleic acids^156^. This rather broad definition includes a diverse group of biophysical systems in different states^3,157,158^. Several transport models propose that the central transporter shares unusual viscoelastic properties with gels, hydrogels, and liquid-liquid phase separations^3,35,62,159,160^. We next consider transport system properties shared with condensates. In the NPC, the scaffold constrains the FG repeats into the relatively narrow space of the central channel through their specific anchoring to the scaffold; thus, the central transporter resembles the concentrated biomolecules in a condensate, but crucially does not replicate the free diffusion of these biomolecules necessary for a true condensate, as the FG repeats are anchored to a static NPC scaffold, with these FG repeats recruiting NTRs and cargo molecules to the NPC^17,75,76,141,142,157^ (**Fig. 3**). Moreover, our model suggests that the physical interactions among FG repeats and NTRs inside the central channel are highly dynamic, resembling superficially some aspects of a liquid-like state of condensates, but also replacing homo-oligomeric FG repeat interactions with FG:NTR interactions; our model instead can accommodate, but does not depend upon, a modest level of “fuzzy” intermolecular interactions between some FG repeats (**Fig. 9a-b**; **Extended Data Fig. 6**). Even with attractive FG:FG interactions, the FG repeats remain highly dynamic and rapidly exchanging with NTRs^40,41^, courtesy of the slide-and-exchange mechanism^43^ (**Figs. 2 and 4i**).

Saturated hydrogels are loosely defined as cross-linked networks of hydrophilic polymer materials that quickly absorb and retain water^35^. They may be considered a specific case of a biomolecular condensate^161^. Similarly to condensates, they cover a diverse group of materials. In the context of the NPC, FG repeats were originally suggested to form a sieve-like hydrogel with a mesh size of approximately 30 kDa for unhindered passive diffusion, providing a firm barrier toward inert molecules^35^. However, this model is inconsistent with the gradual decrease in permeability to passive diffusion in various cell types *in vivo*, including for molecules larger than 30-60 kDa^19,20,82^ (**Fig. 4a; Extended Data Fig. 4a**). The hydrogel model also requires sufficiently strong FG:FG interactions to form the sieve, apparently inconsistent with the highly dynamic nature of FG repeats observed in recent *in situ* experiments^73^ and our model (**Extended Data Fig. 2b**) as well as the ability of the central transporter to efficiently translocate megadalton cargo with the aid of relatively few NTRs^74^ (**Fig. 6-7**).

## Resolving the paradox of transporting megadalton cargos

Considering that the NPC inhibits passive diffusion of relatively small molecules beyond a few tens of kDa^18–20,77^, it may seem paradoxical that it facilitates efficient transport of large pre-ribosomal subunits and mRNA molecules with remarkable efficiency while relying on a relatively small number of NTRs^74,106^. Recently, simulations have shed light on the insertion of the HIV-1 capsid into the NPC^162^ and how it could lead to the cracking of the NPC structure^163^. However, these simulations did not focus on the transport kinetics. In contrast, our spatiotemporal model does quantitatively predict the transport rates of large (MDa) cargo (**Figs. 7-8**). Moreover, it also reveals the NPC features that facilitate this transport without compromising the passive permeability barrier, as discussed above (**Fig. 10**).

## Limitations of the study

Two levels of uncertainty limit our current study. First, the uncertainty of characterizing the behavior of the NPC model using finite simulations. Second, the extent to which our model applies *in vivo*. We attempted to overcome both of these uncertainties *via* validating the simulations by comparison with experimental data (**Figs. 3-4**). The accuracy, precision, completeness, and explanatory power of the model will increase with more experimental data, better modeling methods, and more computing time. For example, the model will benefit from more accurate experimental characterization of the interactions of specific types of NTR:cargo complexes with the different types of FG repeats; and more accurate characterization of the different contributions of individual interaction sites^111^.

## Conclusions

Our reductionist spatiotemporal model of NPC-mediated transport was trained and validated by a wide range of empirical observations on the biochemistry, morphology, and function of the NPC. The model enabled us to dissect key aspects of the transport mechanism and quantify their relative contributions to its functional integrity. These predictions and rationalizations were made possible by relying on (i) integrative modeling to incorporate multiple types of experimental data and physical principles, (ii) a relatively coarse-grained representation of transport to increase model parsimony and computational efficiency, (iii) Bayesian metamodeling to couple transport through a single NPC with the cell-wide RanGTP cycle, and (iv) a significant computational effort to comprehensively map key transport parameters. The model provides a rigorous quantitative starting point for more detailed characterizations of how to rationally modulate and control the transport system or artificial mimics^60,83^ inspired by it. In addition, the present study illustrates how our integrative experimental and computational approach can produce insightful models for other complex biomolecular processes that are refractory to single methods, in turn providing necessary input for realistic and interpretable modeling of an entire cell^99,164–166^.

## Methods

Experimental and computational methods are described in detail in Supplementary Methods.

## Funding

Modeling activities were supported by the National Institutes of Health (NIH) grants R01 GM117212 (D.C.), P41 GM109824 and R01 GM112108 (M.P.R. and A.S.), and R01 GM083960 (A.S.). Our early Brownian dynamics simulations were performed on the Google Exacycle cloud computer from 2013 to 2014; we acknowledge Google salary support for D.R. from 2013 to 2014. Molecular dynamics simulations were also performed on the Anton2 special-purpose supercomputer provided by the National Resource for Biomedical Supercomputing (NRBSC) courtesy of D.E. Shaw Research, the Pittsburgh Supercomputing Center (PSC), and the Biomedical Technology Research Center for Multiscale Modeling of Biological Systems (MMBioS) through grant P41GM103712-S1 from the NIH. SAXS data at NSLS was collected on the LiX beamline, part of the Center for BioMolecular Structure (CBMS), supported by NIH grants (P30GM133893, S10OD012331), and by the Department of Energy (DOE) Office of Biological and Environmental Research (KP1605010). Work performed at the CBMS was supported by the DOE Office of Science, Office of Basic Energy Sciences Program (DE-SC0012704). *In situ* cryo–ET of NPC was supported by NIH grants DP2 GM123494 and U54-AI170856 and NSF grant MRI DBI 1920374 (E.V.). E.V. is an investigator of the Howard Hughes Medical Institute. We acknowledge the use of the UC San Diego cryo-EM facility, which was built and equipped with funds from the University of California, San Diego and an initial gift from Agouron Institute. D.S. was supported by a Damon Runyon Postdoctoral Fellowship (DRG-2364-19) and the K99 Pathway to Independence Award from the NIH (K99AG080112). R.E. and B.R. were supported by a Minerva center grant on Cell Intelligence, Israeli Science Foundation grant 385/24, and a Hebrew University Center for Interdisciplinary Data Science Research grant.

## Acknowledgements

We acknowledge Dr. Frank Alber for initiating our Brownian dynamics simulations of nucleocytoplasmic transport in 2005 as well as Dr. Lin Yang and staff at the LiX NSLS beamline for advice and assistance. We are also grateful to other colleagues for helpful discussions. This work is dedicated to the memory of Anton Zilman.

## Ethics information

Nothing to declare.

## Competing interests

All authors declare no competing interests.

## Extended Data Figures and Tables

**Extended Data Figure 1.**
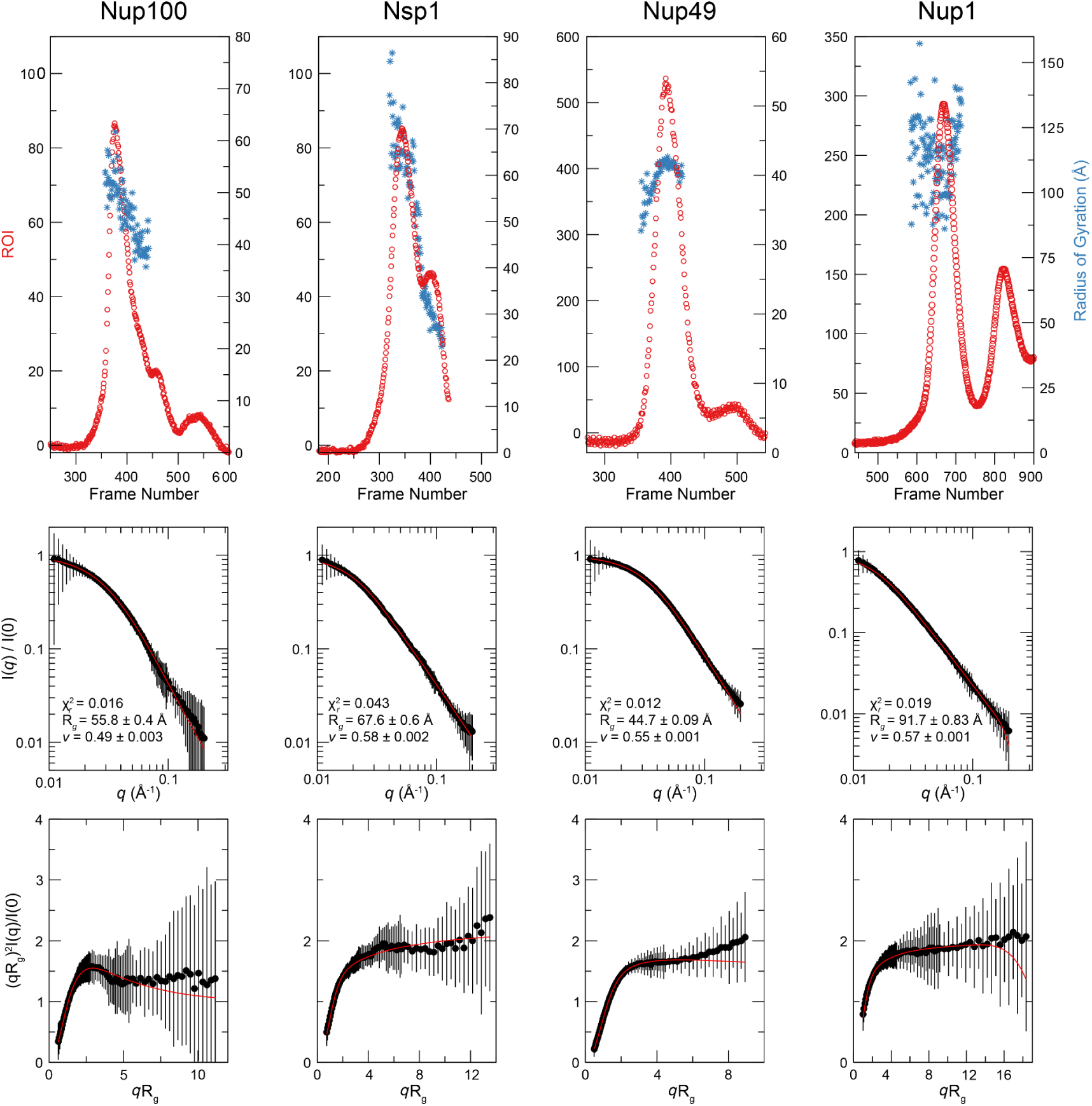
Size exclusion chromatography and SAXS data for quantifying radius of gyration distributions for entire FG repeat domains. These data were used as part of the input information for model construction (**Fig. 1b**; Table 1; Supplementary Tables 1-10). Top row: Distribution of radius of gyration derived from the scattering in the center frames (blue) superimposed on the normalized optical density (red) from the size exclusion runs. Center row: SAXS curves of intensity *vs q* (in units of reciprocal Å) at the peak value of optical density. Bottom row: Normalized (or “dimensionless”) Kratky plot (*sR*_g_)2*I*(*s*)/*I*_(0)_ *vs sR*_g_, characteristic of flexible macromolecules^169^. The horizontal or upward turning left part of each curve demonstrates a low degree of compactness, resulting from an intrinsic lack of rigid structure. The sequence of each FG domain of the four FG Nups is tabulated in Supplementary Results and Methods.

**Extended Data Figure 2.**
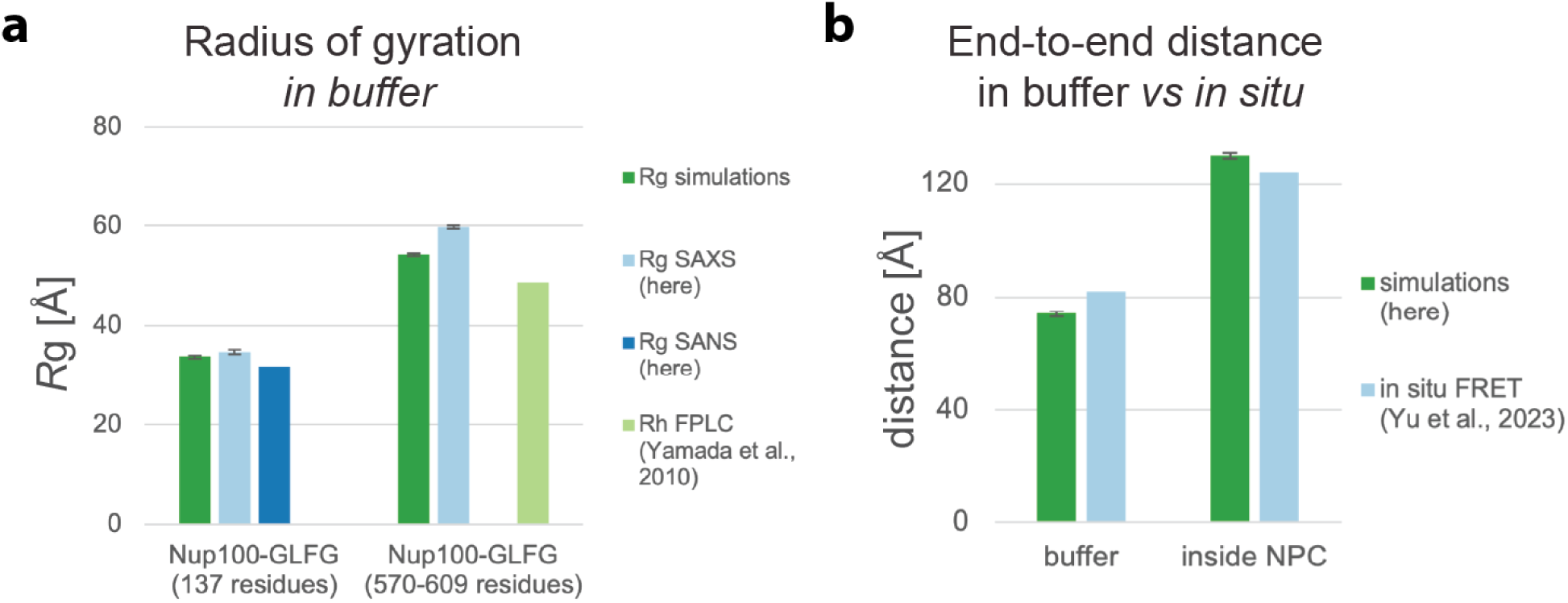
Training and validation of Nup100-GLFG behavior in a buffer and the NPC. **a**, Comparison of empirical and simulated training data on short *vs* long Nup100-GLFG segments in dilute buffer. In the simulations, the short and long segments of Nup100 consist of 140 and 580 residues, respectively, with a single chain simulated in a bounding box of 200×200×200 nm. For the short segment, the SAXS and SANS data were collected for Nup100 residues 318-444. For the long segment, the SAXS data were collected for Nup100 residues 1-602, and the FPLC data were collected for residues 2-610 (**Supplementary Data Tables 2, 10**). **b**, Comparison of simulated and empirical validation data^73^ on end-to-end distance for a 260-residue Nup100 segment in buffer and inside the NPC.

**Extended Data Figure 3.**
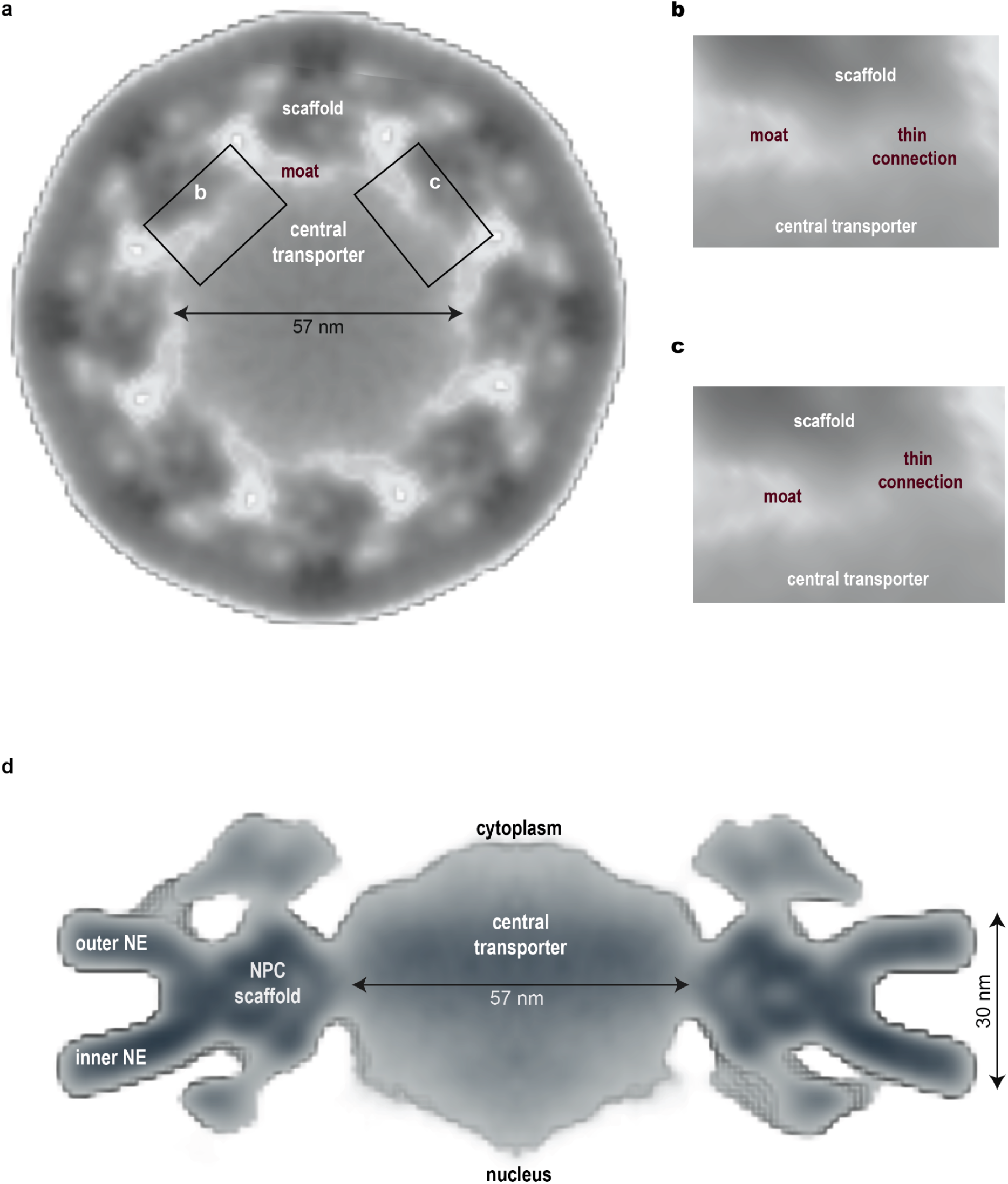
Current *in situ* cryo-ET map of the central transporter. **a**, A top view of the density in the central channel at the midpoint of the NE from the *in situ* cryo-ET map. **b**, and **c**, Zoomed-in views of the moat region indicated in **a. d**, A side view of the NPC scaffold and the central transporter, embedded in the NE.

**Extended Data Figure 4.**
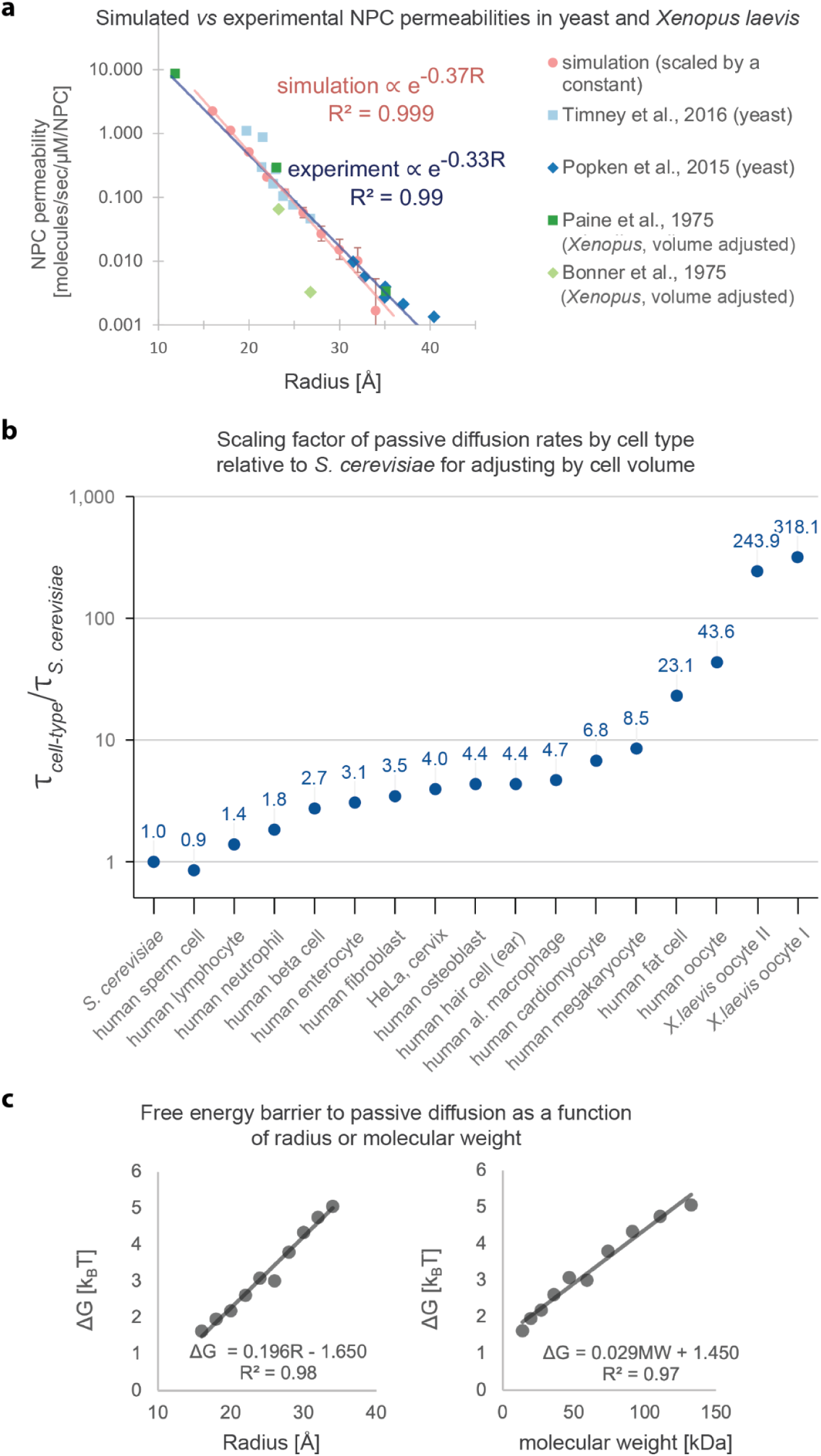
Additional analyses of simulated and measured passive diffusion. **a**, Comparison of simulated (pink) and experimental permeabilities from *in vivo* measurements (**Fig. 3a**), but as a function of estimated molecular radius instead of molecular weight, shown on a log-linear scale. The pink and blue lines are exponential curves fitted to the simulated and experimental measurements in yeast, respectively. **b**, Computed scaling factors for transport time tau in various cell types compared with that for *S. cerevisiae*, based on differences in the volumes of the nuclear and cytoplasmic compartments. Transport times are slower in larger cells due to the longer diffusion times within larger compartments, even when accounting for the larger number of NPCs. The scaling factors for human and *X. laevis* cells are likely to be overestimated due to the volume occupied by the many fat granules in these cell types, although they are not expected to significantly change the analysis. **c**, Gibbs free energy for passively diffusing molecules of different radii (left) or molecular weight (right).

**Extended Data Figure 5.**
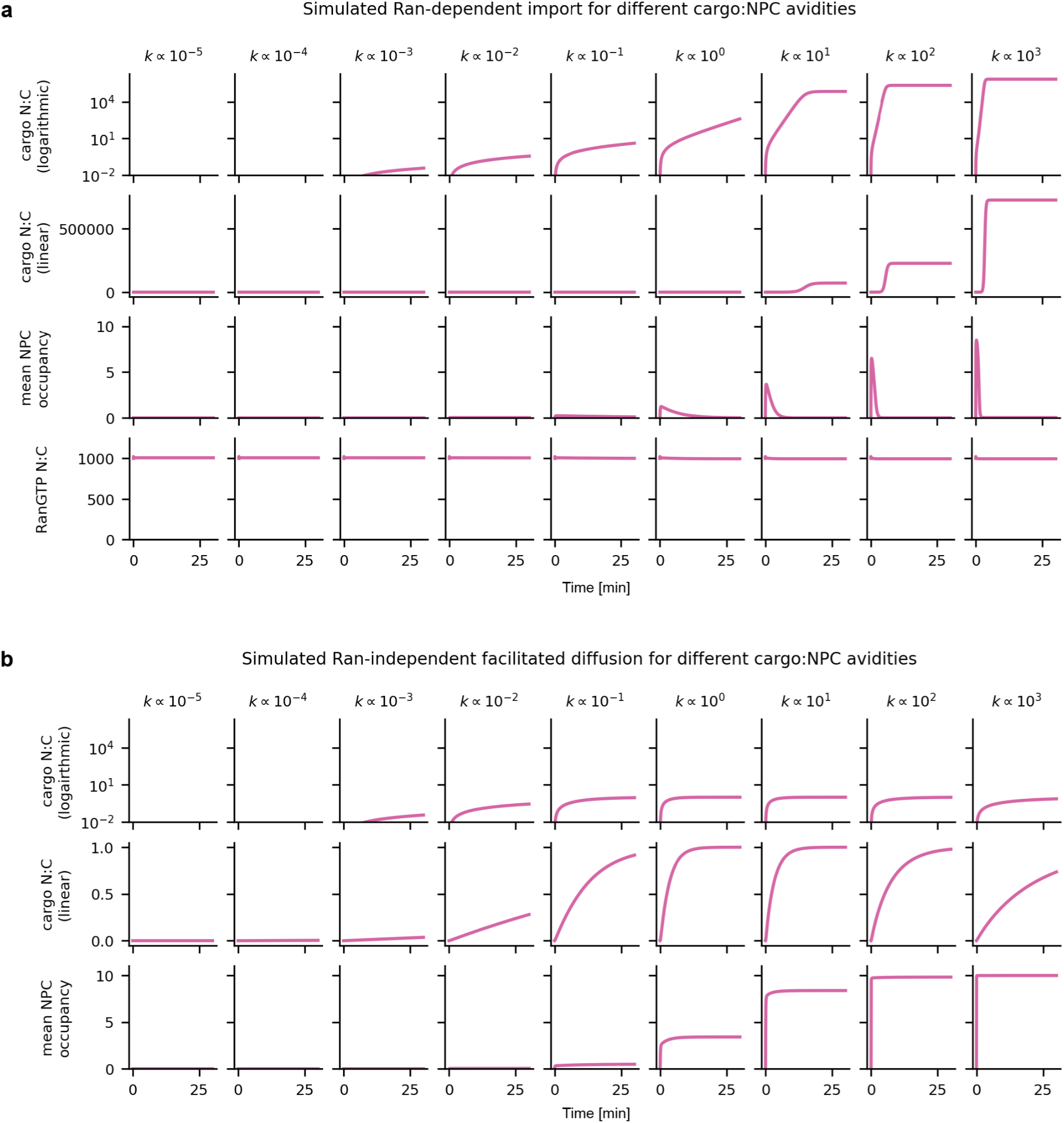
Ran-dependent transport. Simulations of transport or facilitated diffusion for very large cargo using a metamodel that couples our previous kinetic model of transport in the cell^71^ with the current spatiotemporal model of transport through a single NPC. a, Simulations of Ran-dependent transport for varying magnitudes of avidity of the NTR:cargo complexes to the NPC (left to right), showing cargo nuclear-to-cytoplasmic (N:C) ratio on a logarithmic (first row) and linear scales (second row), number of large NTR:cargo complexes occupying each NPC on average (third row); and nuclear-to-cytoplasmic ratio of RanGTP (fourth row). b, Same as the first three rows of (a) for Ran-independent facilitated diffusion. In these simulations, RanGTP does not dissociate NTR:cargo complexes. Thus, they dissociate at the same rate on both the nuclear and cytoplasmic sides, until equilibration of their nuclear-to-cytoplasmic ratio.

**Extended Data Figure 6.**
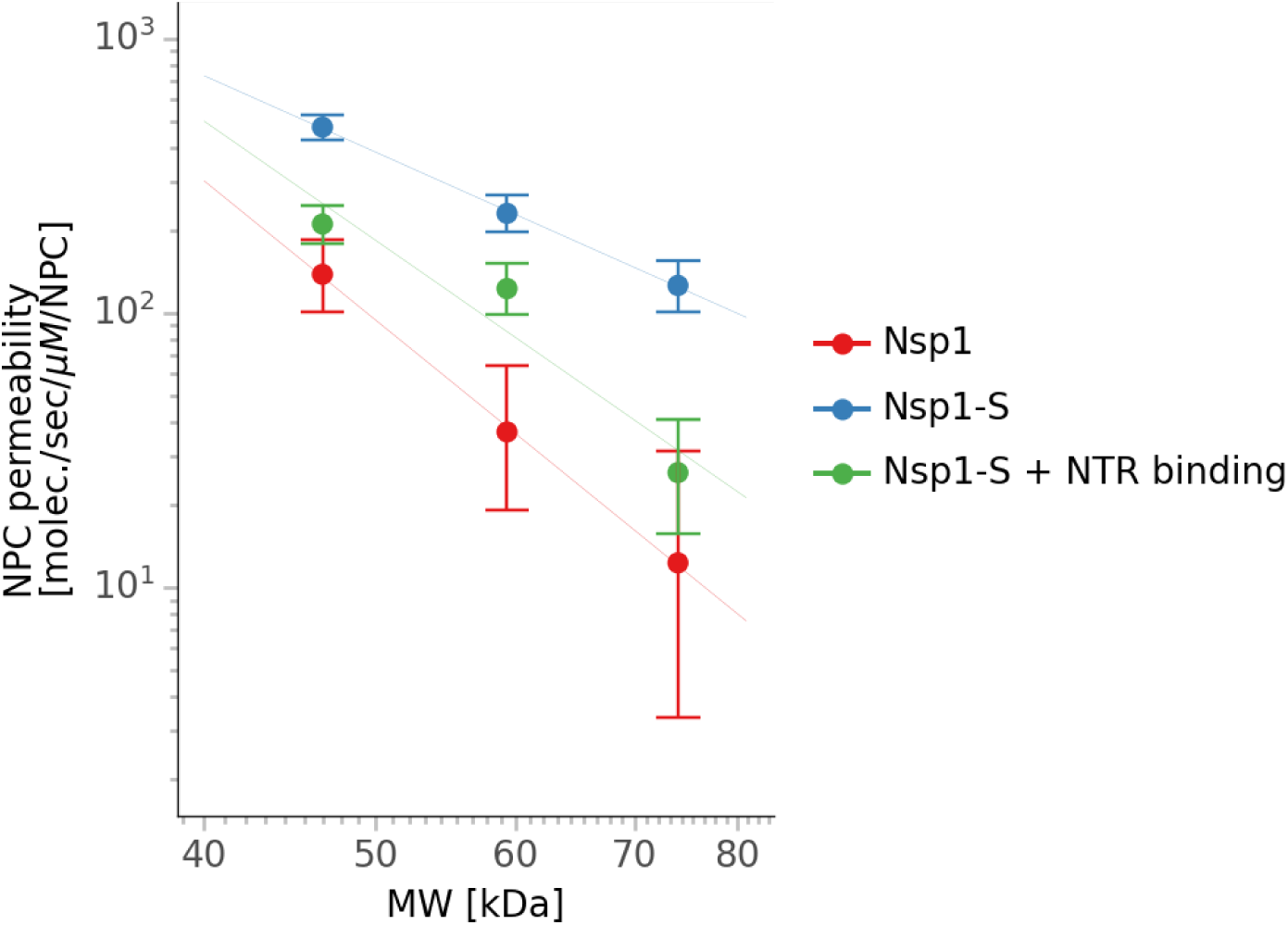
Simulated passive diffusion rates through artificial nanopores^103^. NPC permeability as a function of molecular weight for a cylindrical pore with a radius of 24 nm, lined with 104 chains of Nsp1 (red), Nsp1-S (FG mutated to SG, implying loss of both FG:FG interactions and NTR interactions; blue), and Nsp1-S with restored NTR interactions, but without FG:FG interactions (green).

## Supplementary Videos

**Supplementary Video 1**

https://drive.google.com/open?id=12qSS4Y0xKijj49tyHayOE1JWG29AAXvL&usp=drive_fs

A sample Brownian Dynamics simulation trajectory from our model of transport through the NPC. The scaffold of the NPC is shown in yellow surface representation. FG repeat domains of different FG Nups are anchored to the scaffold (tube representation in various hues). Inert (passively-diffusing) macromolecules that do not interact with the FG repeats are represented as blue spheres of different radii, corresponding to different molecular weights. Note the size dependency of passive diffusion. NTR:cargo complexes that interact with the FG repeats are represented as red spheres of different radii.

**Supplementary Video 2**

https://drive.google.com/file/d/1FzVrpmqUOA2kWnNZ1PZlEUuJHF5dAUgP/view?usp=sharing

A 12 μs Anton 2 simulation of the interaction between Kap95 (gray cartoon representation) in the presence of the Nsp1-derived FSFG6 construct^43^ with six consecutive FSFG repeats (pink representation; phenylalanine residues in pink space-fill representation). The surface representation on Kap95 indicates cumulative contact areas, colored by the fraction of time in contact with FSFG motifs of FSFG6 up until the current point in the simulation, from ∼0% in blue (short transient contacts; for example, probing a local surface non-specifically) to 100% in red (longer-lived contacts in grooves between external helices).

**Supplementary Video 3**

https://drive.google.com/open?id=12zDlvzgAVmDr773OPKcjYAwVdVN2tZG4&usp=drive_fs

Sliding of an FSFG motif (residues 88-91 of FSFG6) about its interaction site with Kap95, as visualized by tracking its center of mass over a 1.5 μs simulation on Anton 2. The simulation itself included two FSFG6 constructs, but only a single FSFG motif from the first chain is shown for clarity. The arrows indicate the principal components computed by applying principal component analysis to the set of FSFG center of mass coordinates, similarly to an analysis of FSFG:NTF2 interactions in Figure 4 of ref.^43^

## Notes

https://docs.google.com/document/d/1pyDS-FGiYeUFrGMy4x7YMr96gezvcS0XQMlTLkf6AVg/edit#heading=h.kacenoxbrhrt

https://drive.google.com/open?id=12qSS4Y0xKijj49tyHayOE1JWG29AAXvL&usp=drive_fs

https://drive.google.com/file/d/1FzVrpmqUOA2kWnNZ1PZlEUuJHF5dAUgP/view?usp=sharing

https://drive.google.com/open?id=12zDlvzgAVmDr773OPKcjYAwVdVN2tZG4&usp=drive_fs

https://github.com/integrativemodeling/npctransport

### Competing Interest Statement

The authors have declared no competing interest.

### Summary of Updates

Substantial additions to the analysis, discussion and the supplementary material

