## Supplementary Material for "Integrative mapping reveals molecular features underlying the mechanism of nucleocytoplasmic transport"

*** Corresponding Authors**

### Table of Contents


**1. Supplementary Results** **3**
1.1 NPC permeability to passive diffusion is conserved across multiple species 3
1.2 Effect of sliding on exchange rate of FG repeats on NTRs 3
**2. Supplementary Methods** **4**
2.1 Integrative spatiotemporal modeling of transport through the NPC 4
Step 1: Gather input information 4
Step 2: Represent nucleocytoplasmic transport 4
Step 3: Compute and filter transport trajectory sets 5
Step 4: Validate the model 5
2.2 Representation of model components, interactions, and dynamics 6
Model components 6
Interactions among model components 7
I. Excluded volume term 7
II. FG polymer term 7
III. FG cohesiveness term 8
IV. FG:NTR interaction term 9
V. Non-specific interaction term 10
Dynamics of the model 10
2.3 Simulations of freely-diffusing FG repeats and NTR:cargo complexes 11
Parameterizing FG repeats to fit empirical radius of gyration and end-to-end distance 11
In silico titrations of FG repeats with a varying number of FG motifs 12
Parameterization and validation of per-site dissociation constant KD,mono 12
Avidities of NTR:cargo complexes for NTRs with varying numbers of FG interaction sites 13
2.4 In situ cryo-electron tomography of S. cerevisiae NPC 14
Tilt-series processing and alignment 14
Generation of subtomograms containing NPCs 14
Model creation, refinement, and classification 15
2.5 Anton2 all-atom simulations 15
System preparation 15
Anton simulations 16
**Supplementary Tables** **17**
Supplementary Table S1. Radius of gyration. 17
Supplementary Table S2. Data on maximal dimension Dmax or QCM-D film thickness. 19
Supplementary Table S3. Persistence length for FSFG constructs from SANS. 21
Supplementary Table S4A. $K_D$ values from NMR and ITC. 22
Supplementary Table S4B. Joint analysis of $K_D$ measurements. 23
Supplementary Table S5. Measured and simulated diffusion coefficients in buffer. 24
Supplementary Table S6. Polymer relaxation time and shape parameters from MD simulations. 26
Supplementary Table S7. Details on SANS measurements of *R*g and maximal dimension *D*max. 27
Supplementary Table S8. Ensemble modeling data of Rg and Dmax based on SAN1. 28
Supplementary Table S9. Constructs used in LH and SS-RH SAXS. 29
Supplementary Table S10. Parameters for FG Nups following model training in the coarse-grained model of transport. 32
Supplementary Table S11. Sequences of FG Nup domains used in Extended Data Fig. 1. 34
**References for Supplementary Material** **36**

### 1. Supplementary Results

#### 1.1 NPC permeability to passive diffusion is conserved across multiple species

We examined whether or not the size-dependence of passive diffusion in yeast is conserved in other organisms and cell types. For example, a *Xenopus* oocyte is a highly specialized cell with a diameter that is 200-300 times larger than that of the yeast cell[1–3], resulting in a seven orders of magnitude larger volume. Moreover, the half-life for the decrease in the difference between the concentrations of the passively diffusing molecules in the cytoplasm and nucleus (*ie*, the equilibration time) is two orders of magnitude longer in a *Xenopus* oocyte than in yeast[2,4,5]. Nevertheless, this observation is rationalized simply by Fick's first law[3], based on the volumes of the cytoplasm and nucleus[2,4] as well as parsimoniously assuming the same NPC surface density in both organisms; in particular, the yeast simulations accurately predict the transport rates for dextrans and ovalbumin in *Xenopus* oocytes (**Fig. 3a**). Thus, the permeability of a single NPC to passively diffusing molecules appears to be conserved across species. A similar calculation for other cell types indicates that *in vivo* equilibration times in most cells are slower but still on the same order of magnitude as in yeast (**Extended Data Fig. 2b**). Thus, our NPC model may accurately quantify NPC transport rates across multiple species, not only yeast. This generality increases our confidence in the nucleocytoplasmic transport mechanism suggested by the model, because it is unlikely that the model conserves the NPC permeability across different species by chance.

#### 1.2 Effect of sliding on exchange rate of FG repeats on NTRs

We assessed two alternative forms of the interaction between FG repeats and NTRs in our coarse-grained model, resulting in a sliding and non-sliding interaction mode, respectively (**Fig. 1g**). For the sliding mode, FG repeats interact with NTRs anisotropically, as observed in the atomistic molecular dynamics simulations[6]. For the non-sliding mode, the interaction is isotropic, depending only on the distance, but not on the relative orientation of the FG motif and its interaction site. In both modes, interaction parameters were optimized to reproduce the empirical interaction affinity for a single FG motif interacting with a single NTR **(Fig. 1g**, left). As predicted by the atomistic molecular dynamics simulations, sliding of competing FG motifs also facilitates faster exchange in the coarse-grained Brownian dynamics simulations; the simulated on- and off-rates (**Fig. 1g**, middle and right, respectively) are an order of magnitude higher for the sliding than non-sliding mode. The accurate reproduction of the rapid exchange between FG repeats and NTRs[7,8] in the context of multivalent interactions[9] justifies using the sliding mode in our coarse-grained model.

#### 2. Supplementary Methods

##### 2.1 Integrative spatiotemporal modeling of transport through the NPC

We used integrative, data-driven coarse-grained Brownian dynamics simulations (**Fig. 1a; Supplementary Video 1**). The integration of multiple types of information allowed us to coarse-grain both the representation of the system and its trajectories, resulting in increased accuracy and efficiency of modeling. The modeling process consisted of the following four steps (**Methods**). First, we gathered varied experimental and theoretical information on the structure and dynamics of the transport system and its various components (**Fig. 1b, Table 1**). Second, we created a coarse-grained Brownian dynamics representation of the transport system (**Fig. 1a**). Third, we computed a set of transport trajectories for each of the many combinations of parameter values (*eg*, interactions and diffusion constants), followed by systematically filtering these sets based on their fit to input information (*eg*, **Fig. 1c,d**, blue). Finally, we validated each filtered set of trajectories by quantifying how well it reproduces experimental data not used in its construction or filtering (*eg*, **Fig. 1c-e**). We outline each one of these 4 steps next, followed by Sections with additional details about modeling.

###### Step 1: Gather input information

In Step 1, input information was derived from multiple experimental and computational methods, providing extensive characterizations of the structure, dynamics, and thermodynamics of the entire yeast NPC (**Fig. 1b**; **Table 1**). These characterizations included published datasets[6,9–12] as well as small-angle X-ray scattering (SAXS) profiles of FG Nups and atomistic molecular dynamics simulations of Kap95 in the presence of FSFG and GLFG repeats. We also relied on additional data for model validation, including published datasets[3,13–17] as well as an *in situ* cryo-electron tomography (cryo-ET) map of the central transporter and atomic force microscopy (AFM) images[18]. The scaffold model, including the localization of the FG Nup anchor domains, was provided by the complete integrative structure model of the yeast NPC[12]. This model is the starting point for the representation of transport (Step 2), computing and filtering of simulation trajectories (Step 3), and validation of the transport model (Step 4). To address the apparent inconsistencies between some datasets, such as those on the FG:FG interactions[10,19–21], we modeled transport for a range of parameter values (Steps 2-4) instead of including these datasets into model construction.

###### Step 2: Represent nucleocytoplasmic transport

The NPC and its environment are a dynamic system of interacting components (**Fig. 1a**). Model components include the NE, the scaffold of the yeast NPC embedded in the NE, the attachment sites of FG repeats to the scaffold, the dynamics of these FG repeats, NTR:cargo complexes with multiple interaction sites for FG repeats, and passively diffusing individual macromolecules that do not interact specifically with FG repeats. Each nucleoporin domain was represented by up to a few spherical beads, totalling 20,272 beads per NPC for the disordered FG repeats alone. Interactions among these components and with their environment were parameterized based on input information. To efficiently model the transport trajectories, we relied on the Brownian dynamics scheme[22]. Brownian dynamics simulations have been previously shown to provide useful dynamic depictions of protein-protein interactions[23], including in the crowded environment of the cell[24] and the NPC[3,12,25].

##### Step 3: Compute and filter transport trajectory sets

We proceeded in two stages to reduce computational cost (**Methods**). First, many combinations of uncertain model design features (*eg*, model granularity and interaction parameter values) were filtered to retain only those model instances that produced trajectories consistent with the experimental data, resulting in an improved fit of the model to a subset of the data. Second, extensive production trajectories were generated using the tuned model.

Brownian dynamics simulations were implemented in our open-source *Integrative Modeling Platform* software (www.integrativemodeling.org)[3,12]. A starting configuration is propagated in ~$10^8$ small steps, reflecting the interactions between the components and small random forces, resulting in a ~0.1 ms trajectory. Because transport is a relatively rapid but rare stochastic event[26,27], thousands of independent trajectories starting from random configurations were computed for each set of conditions, leading to an aggregate simulation time of several seconds.

To compute both the filtering and production trajectories, we had to use extensive computational resources. The filtering stage produced millions of relatively short trajectories (5-10 μs) on Google Exacycle, benefiting from frequent overnight runs on several hundred thousand CPUs, and on thousands of CPUs on our own Linux high-performance computing cluster at UCSF. The subsequent production trajectories required a few million CPU core hours on our cluster.

##### Step 4: Validate the model

The model is validated in four ways, as follows. First, we estimate model uncertainty (precision) corresponding to the variation among a set of independent trajectories. These uncertainties are quantified by the confidence intervals for model properties, such as density maps and transport rates, estimated from the trajectories[28]. Thus, we first verify that the cumulative simulation time resulted in sufficiently precise estimates to justify our conclusions. Second, we ensure that the model, including its representation and parameter values, is consistent with information used and not used to construct it. This goal is achieved by using some input information for defining model representation and filtering transport trajectories, with the remainder of the information used for validating the filtered trajectories. Third, we compensate for the remaining uncertainty about model representation and parameter values by interpreting the trajectories computed for a range of uncertain parameter values. Finally, we use our model to rationalize available experimental and theoretical information, to make new testable predictions, and to provide explanations about the modeled system. In seeking mechanistic insight into the nucleocytoplasmic transport, we benefit from the reductionist nature of the model that maps the system behavior in terms of its components and interactions between them.

#### 2.2 Representation of model components, interactions, and dynamics

The model of nucleocytoplasmic transport was implemented in our open source *Integrative Modeling Platform* (IMP) software (http://integrativemodeling.org)[29–31] using a previously published protocol for Brownian dynamics simulations[3,12]. A detailed manual, input files, and scripts for the simulations are deposited in a GitHub repository (https://github.com/integrativemodeling/npctransport) and at http://integrativemodeling.org/systems/npctransport.

##### Model components

The simulated components include the static ring-shaped scaffold of the NPC, the NE, disordered and flexible FG repeat domains of FG Nups, NTR:cargo complexes, and passively diffusing macromolecules, all enclosed within a bounding box of 2,000 x 2,000 x 2,000 Å. The configuration of these components is fully specified by a configuration vector $X$ that includes their spatial coordinates, their orientation vectors, and the values of some auxiliary variables, as detailed in the following Section “Interactions among model components”.

The integrative model of the NPC scaffold[12] was coarse-grained to increase the computational efficiency of the simulations while maintaining a relatively high accuracy of representation of the FG Nup anchor points: the static scaffold surface was approximated by a collection of spherical particles using the IMP `get_simplified_from_volume()` method, with an approximation error of $min(0.3x(max(2.0, d) - 1)^{1.2}, 5.0)$ nm between the surface of the original representation and the surface of the new coarse-grained representation, where $d$ is the minimal distance between the center of an approximated sphere and any FG Nup anchor point; this approximation implies a deviation from the original NPC scaffold of less than 0.3 nm near the anchor residues of the FG repeats. The NE was represented as a 30 nm slab (based on, for example, the *in situ* cryo-ET map in **Extended Data Fig. 3**) with a toroid pore (outer radius $R$ = 5.4 nm; inner radius $r$ = 39 nm). The disordered domain of each FG Nup (FG repeat domain), typically consisting of 8-53 FG repeats and 250-1000 residues (**Table 2**), was represented as a flexible string of beads (spherical particles), each one of which has a radius of 8 Å and encompasses 20 residues (**Table 2**)[3,12], except for Nup42 that was omitted from the model because of the uncertainty regarding its anchor point as well as a relatively small size and copy number. An FG repeat domain was divided into either GLFG-like, FSFG-like, or non-FG segments, and anchored to the C-terminus of its globular anchor domain (**Table 2**). The number of beads was calculated for each segment separately and rounded up. The FG repeat domain of Nup2, which was not included in the original integrative structure of the NPC[12], was anchored to the same anchor domain as Nup60 based on the known interaction between the two anchor domains[32,33]. Each bead also has a single specific interaction site on its surface, representing an FG motif that may interact with other FG motifs and with NTRs.

Each NTR:cargo complex was represented as a single spherical particle of radius $r$ with $n$ specific interaction sites, randomly distributed on its surface. Each passively diffusing molecule was modeled as a single spherical particle of radius $r$. For conversion between $r$ and molecular weight, we assumed that each sphere has a uniform protein density of 1.38 g/cm$^3$ [34], resulting in $MW = \frac{4}{3}\pi r^3 \rho$ kDa. Alternatively, the radius can be interpreted as the Stokes radius of the molecule, resulting in $MW = (r/6.6)^3$ kDa [35], leading to nearly identical values of the molecular weight as a function of $r$.

**Interactions among model components**

Interactions among the model components were quantified using a coarse-grained potential energy function $U(X)$, where $X$ is the configuration vector:

$$U(X) = U_{\text{excluded}} + U_{\text{FG-polymer}} + U_{\text{FG-cohesiveness}} + U_{\text{FG:NTR}} + U_{\text{non-specific}} \ \text{kcal/mol}$$

where $X$ was omitted on the right side of the equation for simplicity. We now explain each one of these terms in turn.

*I. Excluded volume term*

$$U_{\text{excluded}} = -k_{\text{ex}} \cdot \left( \sum_{\{i,j|\hat{d}_{i,j}<0\}} \hat{d}_{i,j} + \sum_{\{i|\hat{d}_{i,NE}<0\}} \hat{d}_{i,NE} + \sum_{\{i|\hat{d}_{i,BB}<0\}} \hat{d}_{i,BB} \right) \text{kcal/mol}$$

$U_{\text{excluded}}$ is a linear excluded-volume potential that penalizes overlaps between model components. $\hat{d}_{i,j}$ denotes the minimal distance between the surfaces of the $i^{\text{th}}$ and $j^{\text{th}}$ particles; with radii $r_i$ and $r_j$, respectively; it is equal to $d_{i,j} - r_i - r_j$, taking a negative value when the two particles intersect and zero when they touch. Similarly, $\hat{d}_{i,BB}$ denotes the minimal distance between the surface of the $i^{\text{th}}$ particle and the simulation bounding box. $\hat{d}_{i,NE}$ denotes the minimal distance between the surface of the $i^{\text{th}}$ particle and the NE. $k_{\text{ex}}$ is a force constant of 10 kcal/mol/Å, allowing a soft overlap of 1-2 Å between the coarse-grained representations of the model components.

*II. FG polymer term*

$$U_{\text{FG-bond}} = \sum_{FG \in FGs} \sum_{i=0}^{n_{FG}-1} 0.5 \left[ k_{b1} \left( d_{i,i+1} - d'_{i,i+1} \right)^2 + k_{b2} \left( d'_{i,i+1} - d'^{rest}_{i,i+1} \right)^2 \right] \text{kcal/mol}$$

$U_{\text{FG-bond}}$ is a harmonic bonded interaction potential accounting for the spring-like nature of flexible polymers in general[36] and disordered FG repeat domains in particular[6–8,37–40]. $d_{i,i+1}$ denotes the distance between the centers of the $i^{\text{th}}$ and $(i+1)^{\text{th}}$ beads in a single FG repeat domain

$FG \in FGs$ that has $n_{FG}$ beads. For each pair of consecutive beads, the potential includes the sum of two harmonic terms. The first term couples $d_{i,i+1}$ to $d'_{i,i+1}$, a slow-diffusing auxiliary variable representing the momentary rest distance. The second term couples $d'_{i,i+1}$ to $d'^{rest}_{i,i+1}$, the equilibrium rest length between consecutive repeats. The indirect coupling of $d_{i,i+1}$ to $d'^{rest}_{i,i+1}$ *via* $d'_{i,i+1}$ was used to include information about the empirical relaxation time for the end-to-end-distance between consecutive repeats, analogously to the coupling of fast-moving variables to auxiliary slow-moving variables in temperature-accelerated molecular dynamics simulations[41,42]. The value of $d'^{rest}_{i,i+1}$ was typically set to 30.4 Å[6], but it can fluctuate due to model parameter changes when fitting to the data. This value is an input to the simulations; it may differ from the output (apparent) resting distance in the simulations due to, for example, cohesive interactions within or between FG repeats or the constricted volume within the central channel. The value of the first coupling coefficient $k_{b1}$ was set to guarantee a small standard deviation $\sigma_1$ = 2.0 Å of the difference between $d_{i,i+1}$ and $d'_{i,i+1}$. The value of $k_{b1}$ was computed by equating the normal and Boltzmann distributions of the spring length, $e^{-0.5(d_{i,i+1}-\hat{d}_{i,i+1})^2/\sigma^2} = e^{-0.5k_{b1}(d_{i,i+1}-\hat{d}_{i,i+1})^2/k_BT}$, where $k_BT$ is the Boltzmann constant of 0.0019872041 kcal/mol/K [43]) and $T$ is the simulation temperature. From this equality, we inferred that $k_{b1} = k_BT \cdot \sigma_1^{-2}$ $\sigma_1 = \sqrt{k_BT \cdot k_{b1}^{-1}}$. Distance-dependent contributions to configurational entropy can be neglected because $d_{i,i+1} - d'_{i,i+1} << d_{i,i+1}$. Similarly, the value of the second coupling coefficient $k_{b2}$ was typically set to 0.0075 kcal/mol/Å$^2$, implying a standard deviation of $\sqrt{k_BT \cdot k_2^{-1}}$ = 8.9 Å for the distance between two consecutive beads in the FG repeat domains. The diffusion coefficient $D_{i,i+1}$ of the auxiliary variable $\hat{d}_{i,i+1}$ was set to $k_BT \cdot \tau^{-1} \cdot k_{b2}^{-1}$ Å$^2$/fs, where $\tau$ is the estimated relaxation time of the distance between two consecutive FG repeats. It was typically set to 50 ns for all FG repeats unless stated otherwise (**Table 1**), based on the relaxation time of a polymer chain of $\zeta \cdot k_{b2}^{-1}$ [36], where $\zeta$ is the hydrodynamic friction coefficient of the Brownian dynamics, which is equal to $k_bT \cdot D_{i,i+1}^{-1}$ (Section “Dynamics of the model” below).

*III. FG cohesiveness term*

$U_{\text{FG-cohesiveness}} = \sum_{i,j} L(d_{i_0,j_0})$ kcal/mol

$U_{\text{FG-cohesiveness}}$ is an isotropic non-bonded linear interaction potential representing cohesive interactions between pairs of interaction sites on FG repeats. $L(d_{i_0,j_0})$ is a truncated linear restraint on $d_{i_0,j_0}$, the distance between the interaction sites on the $i^{th}$ and $j^{th}$ FG beads. It is equal to $0.5(k_{\psi_i} + k_{\psi_j})d_{i_0,j_0}$ when $d_{i_0,j_0} < 0.5(R_{\psi_i} + R_{\psi_j})$ or zero otherwise, where $\psi_i$ and $\psi_j$ are the flavors of the $i^{th}$ and $j^{th}$ FG beads, respectively; $k_{\psi_i}$ and $k_{\psi_j}$ are the force coefficients for the corresponding flavors, parameterized as follows: $k_{GLFG-like} = 1.55$ kcal/mol/Å; $k_{FSFG-like} = 1.50$ kcal/mol/Å; $k_{disordered} = 1.50$ kcal/mol/Å; and $R_{\psi_i}$ and $R_{\psi_j}$ are the maximal range for cohesive interaction of FG repeats for the corresponding flavors, parameterized to 6 Å for all FG flavors.

*IV. FG:NTR interaction term*

$$U_{\text{FG:NTR}} = 0.5k_{NTR} \cdot \sum_{i_m,j_n} \varphi_{i_m,j_n} H(\hat{d}_{i,j}) \text{ kcal/mol}$$

$U_{\text{FG:NTR}}$ is an anisotropic non-bonded interaction potential between interaction sites on NTRs and FG repeat domains that takes into account both the distance between the FG repeats and the NTR:cargo complex and the relative orientation of their interaction sites with respect to one another. The anisotropy reflects the observed sliding of FG motifs on the surface of NTRs in all-atom molecular dynamics simulations that reproduce experimental R1/R2 relaxation data from NMR and are consistent with the elongated chemical shifts associated with FG interaction sites on NTRs[6]. $k_{NTR}$ is the force coefficient for this interaction term, set to 5.98 kcal/mol/Å$^2$ in all simulations unless stated otherwise. $\varphi_{i_m,j_n}$ is a unitless anisotropic attenuation factor that is equal to 1.0 when the interaction sites of the $m^{th}$ interaction site on the $i^{th}$ NTR:cargo complex and the $n^{th}$ interaction site on the $j^{th}$ FG bead are facing each other and 0.0 when they are rotated beyond a certain angle. This threshold angle is equal to $max(0, \frac{cos(\Theta_{i_m,j}) - cos[max\Theta_1)]}{1 - cos[max\Theta_1]}) \cdot max(0, \frac{cos(\Theta_{2,i,j_n}) - cos[max\Theta_2]}{1 - cos[max\Theta_2]})$, where $\Theta_{i_m,j}$ is the angle formed between the $m^{th}$ interaction site of the $i^{th}$ NTR:cargo complex, the center of the $i^{th}$ NTR:cargo complex, and the center of the $j^{th}$ FG bead; and $\Theta_{i,j_n}$ is the angle formed between the center of the $i^{th}$ NTR:cargo complex, the center of the $j^{th}$ FG bead, and the $n^{th}$ interaction site of the $j^{th}$ FG bead. max$\Theta_1$ and max$\Theta_2$ are both set to $\frac{\Pi}{4}$. $H(\hat{d}_{i,j})$ is a truncated harmonic restraint on $\hat{d}_{i,j}$, the minimal distance between the surfaces of the NTR:cargo complex and the FG bead. It has a maximal range of $R_{NTR}$. When $\hat{d}_{i,j}$ is lower than $\frac{1}{2}R_{NTR}$, it is equal to $\hat{d}_{i,j}^{\,2} - \frac{1}{2}R_{NTR}^{\,2}$; in the range $0.5R_{NTR} < \hat{d}_{i,j} < R_{NTR}$, it is equal to $(R_{NTR} - \hat{d}_{i,j})^2$. $R_{NTR}$ was set to 5.5 Å unless specified otherwise.

*V. Non-specific interaction term*

$$U_{\text{non-specific}} = \sum_{i \in FGs, j} L(\hat{d}_{i,j}) \text{ kcal/mol}$$

$U_{\text{non-specific}}$ is an isotropic non-bonded linear interaction potential representing non-specific interactions between FG repeats and any diffusing particles, including NTR:cargo complexes, passively diffusing molecules, and other FG repeats. $L(\hat{d}_{i,j})$ is a truncated linear restraint on $d_{i,j}$, the minimal distance between the surfaces of the $i^{\text{th}}$ FG bead and the $j^{\text{th}}$ diffusing particle. It has a maximal range of $R_{ns}$. If the second particle is an FG bead, it is equal to $\sqrt{k_{ns[\Psi_i]} k_{ns[\psi_j]}} d_{i,j}$ when $d_{i,j} < R_{ns}$ or zero otherwise, where $k_{ns[\psi_i]}$ and $k_{ns[\psi_j]}$ are the non-specific force coefficients for flavors $\psi_i$ and $\psi_j$ of the corresponding beads. If the second particle is not an FG bead, it is equal to $k_{ns[\Psi_i]} d_{i,j}$ when $d_{i,j} < R_{ns}$ or zero otherwise, where $k_{ns[\psi_i]}$ is the non-specific force coefficients for flavors $\psi_i$. Unless stated otherwise, $k_{ns[FSFG-like]}$ was set to 0.01 kcal/Å, $k_{ns[GLFG-like]}$ was set to 0.08 kcal/mol/Å, $k_{ns[Nup159]}$ was set to 0.07 kcal/mol/Å, and $k_{ns[disordered]}$ was set to 0.01 kcal/mol/Å. $R_{ns}$ was set to 5.0 Å for all FG flavors.

#### Dynamics of the model

Brownian dynamics simulations[22] were implemented in IMP as described previously[3,12]. All simulations were conducted at 297.15 K. The force vector $\vec{f}(X_i)$ is acting on the $i^{\text{th}}$ particle with coordinates $X_i$. It is equal in magnitude and opposite in direction to the gradient $\nabla U(X_i)$. The coordinates of all non-static particles (FG repeat domains, NTR:cargo complexes, and passively diffusing molecules) were updated at each time step using the following discrete integration equation:

$$X_i(t + \Delta t) = X_i(t) + \frac{D}{k_B T} \Delta t \vec{f}(X_i) + \sqrt{2D\Delta t} R$$

where $t$ denotes time, $\Delta t$ is the integration time step, $D$ is a translational diffusion coefficient assigned to each bead in units of Å$^2$/fs, $k_B T$ is as described above, and $R$ is a standard normal random variable with the mean of 0 and the standard deviation of 1. The term $\frac{D}{k_B T}$ is denoted $\zeta$, the hydrodynamic friction coefficient for the diffusing particles. Torques between interaction sites on FG repeats and NTR:cargo complexes were integrated similarly to the integration of translational forces, but using a rotational diffusion coefficient $D_{rot}$ instead of $D$, specified in units of rad$^2$/fs. The sum of the torques was multiplied by $\frac{1}{k_B T} D_{rot} \Delta t$ , followed by adding it to a random rotation about a uniformly sampled rotation axis; the magnitude of the rotation is a normal random variable with mean 0 and standard deviation of $\sqrt{6D_{rot}\Delta t}$, approximating an independent rotation around three rotational degrees of freedom of magnitude $\sqrt{2D_{rot}\Delta t}$ each.

To approximate the differences between diffusion rates of molecules of different sizes, we defined the Stokes radius of each bead to equal its radius $r$. We then assigned each bead a translational diffusion coefficient in units of Å$^2$/fs using the Stokes-Einstein equation, $D = \frac{k_B T}{6\pi\eta r}$ , where η=0.92 mPa·s is the dynamic viscosity of water at 297.15 K. Each bead was also assigned a random rotational diffusion coefficient $D_{rot} = \frac{k_B t}{8\pi\eta r^3}$, in units of rad$^2$/fs, using the Einstein-Stokes-Debye equation, and multiplying η by a constant factor of 3.33 to account for increased viscosity in the crowded molecular environment[44].

#### 2.3 Simulations of freely-diffusing FG repeats and NTR:cargo complexes

To fit the parameters of the FG repeats and their interactions with NTR:cargo complexes to biochemical measurements, we simulated freely-diffusing FG repeats and NTR:cargo complexes as follows.

##### Parameterizing FG repeats to fit empirical radius of gyration and end-to-end distance

A dilute solution with a single FG repeat molecule at a concentration of 208 nM was simulated in a cubic bounding box of edge length *d*, specified in units of angstroms *(d=2,000* Å by default*)*. The representation of the FG motifs are as described above, using 6 beads, representing 120 residue repeats, and control simulations using 30 beads, representing 600 residue repeats. We mapped how the $U_{\text{FG-cohesiveness}}$ term in the model's potential energy function (Section "Interactions among model components" above) influences the radius of gyration and end-to-end distance of the FG repeats. Specifically, we repeated the simulations for all value combinations of the force coefficient $k_{\psi_i}$ and the range coefficient $R_{\psi_i}$, assuming $k_{\Psi_i} = k_{\psi_j}$ and $R_{\psi_i} = R_{\psi_j}$ (value range of 0.01-3.0 kcal/mol/Å for $k_{\psi_i}$ in 0.01 kcal/mol/Å intervals, and 4.0-10.0 Å for $R_{\psi_i}$ in 0.1 Å intervals). For each parameter combination, the dynamics of the entire system was simulated for 1,000 μs in 100 independent runs. The first 10 μs of each simulation were used as equilibration time and were discarded from the subsequent statistical analysis. From the simulation results, we quantified the mean and standard deviation of the radius of gyration and end-to-end-distance. Following fitting of the parameters to the empirical data on observed end-to-end distance and radius of gyration (**Supplementary Tables S1-2**), the parameters were set as follows: $k_{GLFG-like} = 1.55$ kcal/mol/Å; $k_{FSFG-like} = 1.50$ kcal/mol/Å; $k_{disordered} = 1.50$ kcal/mol/Å; a uniform value for $R_{\psi_i}$ and $R_{\psi_j}$ of 6 Å for all FG flavors.

#### *In silico* titrations of FG repeats with a varying number of FG motifs

In the *in silico* titration experiments, FG repeat molecules and NTRs were simulated with approximately 529,200 different combinations of FG repeat concentration, NTR:cargo concentration, interaction valencies, and interaction parameters. Each FG repeat molecule was represented using a flexible string of six beads, using the FSFG-like repeat parameters described above, , and corresponding to an approximately 120 residue construct with six FG repeats. As in the experimental construct library, we repeated the simulations with FG repeat molecules containing 1-6 consecutive FG motifs, which interact specifically with NTRs, and 0-5 “inactive” SG motifs, which do not interact specifically with NTRs[9]. NTRs were modeled as described above, with dimensions similar to those of NTF2 (radius of 20.0 Å; PDB 1GYB), with 1, 2, or 4 interaction patches distributed randomly on the sphere’s surface, as described above. The $K_{D,\mathrm{mono}}$ of an individual NTR site with an FG motif was set to 1.25, 2.5, 5.0, 10.0, 20.0, or 40.0 mM, encompassing a previously estimated range[9]. The parametrization of $K_{D,\mathrm{mono}}$ is described below. The concentration of FG repeats was set to 70 different values spaced equally on a logarithmic scale, in the range of 0.1 µM - 250 mM. The concentration of NTR was set in the same range. The concentration of FG repeat molecules and NTRs in the various simulations was mapped in the range of 0.01 µM - 250 mM. The minimal number of FG repeat molecules and NTRs was set to 32 and the maximal number was 10,000, adjusting both the number of molecules of each type and the box size to obtain the desired concentrations as closely as possible. For each of the 529,200 simulations, the simulation time was sufficient to estimate the interaction $K_D$ with precision better than 20%; $K_D$ was estimated from the fraction of bound FG repeat molecules and NTRs, based on the equation $K_D$=[FG][NTR]/[FG:NTR].

#### Parameterization and validation of per-site dissociation constant $K_{D,\mathrm{mono}}$

$K_{D,\mathrm{mono}}$ is the hypothetical interaction dissociation constant between a single FG motif and a single NTR interaction site, when they interact in a monovalent context. $K_{D,\mathrm{mono}}$ cannot be measured directly, as NTRs contain multiple interaction sites for FG motifs. However, we can fit the value of $K_{D,\mathrm{mono}}$, to reproduce measured avidities and titration curves for a subset of the experimental NMR and ITC titration data, followed by their validation by titration data not used for the fitting[9]. Specifically, based on NMR and ITC titrations, the $K_D$ for the interaction between NTF2 and the FSFG1 construct (six repeats with a single FSFG motif and five mutated SSSG motifs) has been estimated to be in the range of 2.2 - 4.5 mM[9]. We fitted $K_{D,\mathrm{mono}}$, to these data, including to the raw titration curves, by repeating the *in silico* titrations for different values of $K_{D,\mathrm{mono}}$ (1.25-40 mM). A good fit was obtained for a $K_{D,\mathrm{mono}}$ value of 40 mM (**Fig. 1c-e,** $FG_1$). The avidities between FG repeats and NTRs are expected to be substantially lower (stronger) than $K_{D,\mathrm{mono}}$, due to site multiplicity on both the FG repeats and NTRs (Section “Avidities of NTRs with varying numbers of FG interaction sites” below). The parameter fitting was followed by validation using titration data not used for training (**Fig. 1c-e**, $FG_2$-$FG_6$).

**Mapping the model parameters to the per-site dissociation constant $K_{D,mono}$ for sliding (anisotropic, default) interaction potential**

As $K_{D,mono}$ is not a direct input to the simulation, we first computed a mapping between a subset of the model parameters and $K_{D,\text{mono}}$. Specifically, we performed auxiliary simulations with a single interaction site and a single FG repeat with a single FG motif (a single bead), for different values of $k_{\text{NTR}}$ and $R_{NTR}$, the input force constant, and range parameters for the interaction between pairs of interaction sites on FGs and NTRs. We then computed $K_{D,\text{mono}}$ for each value of $k_{FG\text{-}NTR}$ from the fraction of bound and unbound FG and NTR molecules over time using the equation $K_{D,\text{mono}}$=[FG][NTR]/[FG:NTR]. Finally, we used non-linear regression to relate $k_{\text{FG-NTR}}$ and $K_{D,\text{mono}}$ *via* a log-quadratic relation, allowing us to compute the input value of $k_{NTR}$ and $R_{NTR}$ that results in per-site affinity of $K_{D,\ mono}$. Specifically, when fixing all other interaction parameters as in **Supplementary Table S10** and varying $k_{NTR}$ and $R_{NTR}$, we obtain the following log-quadratic relation between the input model parameters $k_{NTR}$ and $R_{NTR}$ and the apparent value of $K_{D,\ mono}$:

$$\log(K_{D,mono}) = Q_0 + Q_1 R_{NTR} + Q_2 k_{NTR} + Q_3 R_{NTR} k_{NTR} + Q_4 {R_{NTR}}^2 + Q_5 {k_{NTR}}^2$$

Where the coefficients are as follows:

$Q_0 =- 1.32828572$; $Q_1 = 0.555793607$; $Q_2 = 0.336939985$; $Q_3 =- 0.142452838$; $Q_4 =- 0.0642247161$; $Q_5 =- 0.000576784272$

**Mapping the model parameters to the per-site dissociation constant $K_{D,mono}$ for non-sliding (isotropic) interaction potential**

We also repeated the same process to estimate the log-quadratic relation between the input model parameters $k_{NTR}$, $R_{NTR}$, and the apparent value of $K_{D,\ mono}$ for a non-sliding interaction potential, where $k_{NTR}$ is a constant repulsion force specified in units of kcal/mol/Å within the range $R_{NTR}$:

$Q_0 = 7.465229634$; $Q_1 =- 1.368456938$; $Q_2 =- 0.6409987219$; $Q_3 =- 0.3562974146$; $Q_4 = 0.07905547031$; $Q_5 = 0.01909698241$

**Avidities of NTR:cargo complexes for NTRs with varying numbers of FG interaction sites**

To determine avidities of NTR:cargo complexes as a function of NTR valency, that is, the number of FG motif binding sites on the NTR, we simulated FG repeat constructs with six repeats (six beads) and 145 kDa NTR:cargo complexes (radius of 35 Å, approximating the NTR:cargo complex for karyopherins such as Kap95[45]) using a range of valencies (1-12 interaction sites). We then estimated the avidities of the NTR:cargo complexes to the FG repeat constructs as a function of the valency.

For NTRs with 1 to 8 interaction sites, we simulated the system using the FG repeat concentration of 36 μM (585 chains) and NTR:cargo complex concentration of 5 μM (81 chains), within a simulation box of 3,000 Å x 3,000 Å x 3,000 Å. For each number of NTR interaction sites, the simulation was 40 μs long. The chain avidity was determined from the fraction of bound FG repeats and NTRs during the last 20 μs of the simulation, using the equation $K_D$=[FG][NTR]/[FG:NTR].

For NTRs with 8 to12 interaction sites, we simulated the system using FG repeat concentration of 1 μM (602 chains) and NTR:cargo complex concentration of 0.150 μM (90 chains) in a simulation box of 10,000 Å x 10,000 Å x 10,000 Å. For each number of NTR interaction sites, the simulation was 1000 μs long, following a 10 μs warm-up; this longer simulation time was needed due to the lower concentration and thus lower collision rate. The avidity was determined by estimating the off rate per ns, and the on rate per μM, and then using the equation $K_D=k_{off}/k_{on}$. 95% Poisson confidence intervals were estimated for the observed off and on events[46], resulting in the 95% confidence intervals of the ratio of $k_{off}$ and $k_{on}$. The avidity for NTRs with 8 interaction sites was similar to that observed in a higher concentration above; thus, we used the former value.

#### 2.4 *In situ* cryo-electron tomography of *S. cerevisiae* NPC

*Saccharomyces cerevisiae* W303 cells, collected during their logarithmic growth phase, were deposited onto electron microscopy grids and rapidly vitrified by plunge-freezing into liquid ethane-propane mixture. In cryogenic conditions, lamellae were generated from the clusters of yeast cells using cryogenic-focused ion beam (cryo-FIB) milling in an Aquilos Dual-Beam system (Thermo Fisher Scientific)[47]. The tilt-series data was collected on a Titan Krios G3 electron microscope (Thermo Fisher Scientific) operated at 300 kV, with a post-column energy filter and K2 Summit direct detector (Gatan) used for counting and dose fractionation modes. Parameters for the tilt-series were: Tilt range of ± 45-60°, pixel size of 3.45 Å, tilt increments of 3° (4-5° for some), and a defocus range of -2 to -11 μm. SerialEM was used for data collection[48]. Additional data from EMPIAR-10466 were integrated into the dataset.

###### Tilt-series processing and alignment

The tilt image frames underwent motion correction *via* WARP[49], applying a motion model of a 3x3 grid. The corrected frames were then collated into tilt-series stacks in WARP. These motion-corrected series were subsequently aligned using AreTomo[50]. Post alignment, the stacks were reintegrated into WARP for CTF estimation, determination of defocus handedness, and final reconstruction. All tilt series CTF estimations and defocus handedness were manually checked and adjusted if necessary.

###### Generation of subtomograms containing NPCs

From the aligned tilt-series, tomograms were reconstructed in WARP[49]. The NPC particles in these tomograms were manually picked and assigned initial orientations normal to the NE. The subtomograms of the individual NPC particles, totaling ~4000, were generated in WARP at pixel size of 10 Å.

##### Model creation, refinement, and classification

A select group of NPC particles were used to generate an initial $C_8$ symmetrized model. This initial model was used for initial refinement, with $C_8$ symmetry, using localized searches around the initial orientation (*ie*, initial Euler angles). After the refinement, atypical particles were removed using 3D classification and manual inspection. The acceptable particles underwent further refinement with the $C_8$ symmetry to generate the final *in situ* map. These analyses were performed in Relion 3[51].

#### 2.5 Anton2 all-atom simulations

The simulation boxes for the FSFG and GLFG constructs simulated in this study in the presence of Kap95 or without NTR are:

| FG Nup | Receptor protein | PDB-ID (receptor) | Simulation time |
|---|---|---|---|
| Nsp1-$FSFG_{125}$ | Kap95 | 5OWU:A 1-861 | 20.0 µs |
| Nsp1-$FSFG_{125}$ (*restrained to Kap95 based on 5OWU interaction site)* | Kap95 | 5OWU:A 1-861 | 2.8 µs |
| 2 x Nsp1-$FSFG_{125}$ | Kap95 | 5OWU:A 1-861 | 14.2 µs |
| Nup100-$GFLG_{132}$ | Kap95 | 5OWU:A 1-861 | 3.5 µs |
| 2 x Nup100-$GFLG_{132}$ | Kap95 | 5OWU:A 1-861 | 25.0 µs |
| Nsp1-$FSFG_{125}$ | Maltose Binding Protein (MBP) *(control)* | 1OMP:A 1-370 | 14.0 µs |

**Nsp1-FGFG125 sequence:**

MDNKTTNTTPSFSFGAKSDENKAGATSKPAFSFGAKPEEKKDDNSSKPAFSFGAKSNEDKQD GTAKPAFSFGAKPAEKNNNETSKPAFSFGAKSDEKKDGDASKPAFSFGAKPDENKASATSKPA

**Nup100-GLFG132 sequence:**

MSLFGKANTFSNSASGGLFGQNNQQQGSGLFGQNSQTSGSSGLFGQNNQKQPNTFTQSNT GIGLFGQNNNQQQQSTGLFGAKPAGTTGSLFGGNSSTQPNSLFGTTNVPTSNTQSQQGNSLF GATKLTSNLE

##### System preparation

We constructed an initial model of the disordered Nsp1-$FSFG_{125}$ construct from Nsp1 and the $GLFG_{132}$ construct from Nup100 in complex with Kap95, and used this model as a template for other systems, as follows.

*Bound Kap95:Nsp1-FSFG₁₂₅ system.* Initial coordinates for the NTR structure were taken from chain A of the 5OWU PDB entry[52], where Kap95 was initially in complex with Nup1 (chain B). In the restrained simulation, the binding site for the FG motif of Nup1 (residues 1008-1009) was used to initially position one of the FSFG repeats of Nsp1-*FSFG*$_{125}$, otherwise the FG repeats were initially positioned away from Kap95. The construct was minimized using the AMBER99SB force field [53] in an implicit-solvent box using a modified generalized Born solvation model [54], with strong (500 kcal mol$^{-1}$ Å$^{-2}$) positional restraints on the atomic coordinates of Kap95 and the bound FG repeat constructs.

*Equilibration.* Missing hydrogen atoms were added using the *reduce* tool in AMBER. Crystal water molecules from the 5OWU PDB structure were added, if the water oxygen atom was over 1.5 Å from any protein atom. $Na^+$ counterions were added to neutralize the protein charge, and the protein was aligned by its principal axes using the *alignAxes* command in the `tleap` AMBER tool, and solvated in an orthorhombic box with anisometric solvent buffering. Solvent was added and parameterized based on the TIP4PD water model[55]. Simulations were performed using AMBER99SB-ILDN [56]. Periodic boundary conditions were imposed. For equilibration simulations in AMBER, short-range electrostatic interactions and van der Waals interactions were included with a cutoff of 8 Å. Long-range electrostatic interactions were treated using the periodic mesh Ewald algorithm [57]. Each box was minimized for 250 steepest descent steps, followed by 250 conjugate gradient descent steps. This relaxation was followed by a second minimization with a 10 kcal mol$^{-1}$ Å$^{-2}$ positional restraint on all protein atoms, using steepest descent minimization for 250 steps and conjugate gradient minimization for 750 steps. Each box was then heated to 300 K over 20 ps using Langevin dynamics with a collision frequency of 1 ps$^{-1}$, followed by equilibration in the NPT ensemble for 200 ps, using a pressure of 1 atm and a relaxation time of 2 ps. Subsequently, the system was equilibrated for another 10 ns with AMBER before switching to Anton [58,59]. The time step for the AMBER simulations was 2 fs.

##### Anton simulations

Long simulations were run on the Anton2 supercomputer at the Pittsburgh Supercomputing Center [60]. These simulations also used a 2 fs time step. The Gaussian-split Ewald algorithm [61] was used to compute electrostatic interactions. Parameters for the simulation were informed by the system-optimized parameters suggested by the Anton software. The van der Waals interactions and the direct part of the electrostatic interaction calculations were calculated for all atoms within a cutoff of at least 11 Å for all boxes. The Multigrator thermostat/barostat [62] was used to keep the simulation temperature at 300 K and the pressure at 1 atm.

#### Supplementary Tables

**Supplementary Table S1.** Radius of gyration.
Mean radius of gyration from small-angle X-ray scattering (SAXS; here; also see **Extended Data Figure S1**), small-angle neutron scattering (SANS)[11] (**Supp. Table S7 in ref.**[9]), atomistic molecular dynamics simulations on Anton supercomputer (MD; here and in ref.[6] for FSFGs), FPLC[10], and a theoretical estimate for different Flory coefficients[36].

| **FG type** | **nres** | **# repeats** | **MW [kDa]** | **$R_g$ SAXS [Å]** | **$R_g$ SANS [Å]** | **$R_g$ MD [Å]** | **$R_s$ Yamada et al. 2010 FPLC [Å]** | **Theoretical $R_g$ of 120 residues at Flory coefficient 0.4/0.5/0.6 [Å]** |
|---|---|---|---|---|---|---|---|---|
| FSFG-K free (Nsp1) | 133 | 6 | 14 | 37.75 ± 0.47 | - | 31.6 ± 3.7 | - | 36.2 / 35.9 / 35.5 (SAXS)<br>31.1 / 31.0 / 30.8 (MD) |
| FG-124 | 137 | 9 | 14 | 34.83 ± 0.20 | - | - | - | 32.2 / 32.6 / 33.0 |
| FSFG2 free | 60 | 2 | 5 | 25.57 ± 0.56 | - | - | - | 33.7 / 36.2 / 38.8 |
| FSFG6:NTF2 | 125 (+8) | 6 | 13 | 41.75 ± 0.35<br>600 μM:600 μM | - | - | - | 39.3 / 39.7 / 40.1 |
| FSFG2:NTF2 | 54 | 2 | 13 | 24.42 ± 0.05<br>800 μM:800 μM | - | 21.7 ± 4.7 | - | 33.6 / 36.4 / 39.4 (SAXS)29.9 / 32.3 / 35.0 (MD) |
| GLFG (M-Nup100(318-444)SNLEHHHHH) | 138 | 8 | 13 | 34.73 ± 0.2 | 31.75 +/- 1.07 | - | - | 32.8 / 32.4 / 31.9 (SAXS)<br>30.0 / 29.6 / 29.2 (SANS) |
| Nsp1 30-591 | 571 | 32 | 58 | 75.11 ± 0.27 | - | - | - | 40.2 / 34.4 / 29.5 |
| Nup100 1-570 | 570 | 44 | 58 | 59.68 ± 0.29 | - | - | - | 32.0 / 27.4 / 23.4 |
| Nsp1n 1-172 | 172 | 12 | 18 | - | - | - | 27.1 ± 0.0 | 23.5 / 22.6 / 21.8 |
| Nup116m 165-715 | 551 | 42 | 55 | - | - | - | 46.5 ± 0.0 | 25.3 / 21.7 / 18.6 |
| Nup100n 2-610 | 609 | 43 | 63 | - | - | - | 48.7 ± 0.4 | 25.4 / 21.6 / 18.4 |
| Nup49 1-215 | 215 | 16 | 21 | - | - | - | 26.9 ± 0.0 | 21.3 / 20.1 / 19.0 |
| Nup49 1-269 (RH/JFM/IN design) | 277 | 17 | 27 | 46.07 ± 0.22 (debye analysis) | - | - | - | 33.4 / 30.8 / 28.4 |

| FG type | nres | # repeats | MW [kDa] | $R_g$ SAXS [Å] | $R_g$ SANS [Å] | $R_g$ MD [Å] | $R_s$ Yamada et al. 2010 FPLC [Å] | Theoretical $R_g$ of 120 residues at Flory coefficient 0.4/0.5/0.6 [Å] |
|---|---|---|---|---|---|---|---|---|
| Nup42 1-212 | 212 | 18 | 21 | - | - | - | 28.4 ± 0.5 | 22.6 / 21.4 / 20.2 |
| Nup57 1-255 | 255 | 16 | 26 | - | - | - | 31.1 ± 1.0 | 23.0 / 21.3 / 19.8 |
| Nup145N 1-242 | 242 | 13 | 26 | - | - | - | 28.2 ± 0.2 | 21.3 / 19.9 / 18.5 |
| Nup1c 798-1076 | 279 | 10 | 29 | - | - | - | 32.4 ± 0.4 | 23.1 / 21.2 / 19.5 |
| Nup159 441-881 | 441 | 26 | 46 | - | - | - | 55.4 ± 0.2 | 32.9 / 28.9 / 25.4 |
| Nup60 389-539 | 151 | 4 | 17 | - | - | - | 31.3 ± 0.2 | 28.6 / 27.9 / 27.3 |
| Nup1m 220-797 | 578 | 20 | 64 | - | - | - | 67.9 ± 0.2 | 36.2 / 30.9 / 26.4 |
| Nup2 186-561 | 376 | 16 | 41 | - | - | - | 59.8 ±0.3 | 37.9 / 33.8 / 30.1 |
| Nsp1m 173-603 | 431 | 23 | 46 | - | - | - | 65.3 ± 0.1 | 39.2 / 34.5 / 30.3 |
| Nup145Ns 243-433 | 191 | 1 | 23 | - | - | - | 29.8 ± 0.0 | 24.7 / 23.6 / 22.5 |
| Nup100s 611-800 | 190 | 0 | 23 | - | - | - | 36.6 ± 0.3 | 30.5 / 29.1 / 27.8 |
| Nup116s 765-960 | 196 | 0 | 24 | - | - | - | 39.1 ± 0.2 | 32.1 / 30.6 / 29.1 |
| Nup116 348-458 | 111 | 10 | 13 | - | - | - | 20.4 ± 0.1 | 21.0 / 21.2 / 21.4 |
| Nup116 charged 348-458 (mutation) | 111 | 10 | 13 | - | - | - | 27.2 ± 0.1 | 28.1 / 28.3 / 28.5 |
| Nsp1 377-471 | 95 | 6 | 12 | - | - | - | 26.8 ± 0.7 | 29.4 / 30.1 / 30.8 |
| Nsp1 F>S (mutation) 377-471 | 95 | 0 | 11 | - | - | - | 28.3 ± 0.2 | 31.1 / 31.8 / 32.6 |
| Nup1FG(352-1076)-LEHHHH Ryo+Sam construct | 734 | 17 | 76 | 102.29 ± 18.15 Å (from radial distribution) | - | - | - | 49.6 / 41.35 / 34.5 |

**Supplementary Table S2.** Data on maximal dimension $D_{max}$ or QCM-D film thickness.
Maximal dimension $D_{max}$ in Å units from SAXS (here), SANS[11], maximal end-to-end distance in atomistic molecular dynamics simulations on the Anton supercomputer (here; except FSFG-K in ref.[6]), or film thickness from quartz crystal microbalance with diffusion dissipation (QCM-D)[21].

| | **SAXS [Å]** | **SANS [Å]** | **MD** | **QCM-D film thickness [Å] (NTR concentration)** |
|---|---|---|---|---|
| FSFG-K (Nsp1) | 125.4<br>(8 mg/mL) | - | - | - |
| Regular FSFG repeats (315 AA, 16 FxFG motifs); 6.1 pmol/cm$^2$ grafting density; at different NTF2 concentrations | - | - | - | 100-160 (0 µM)<br><br>150-200 (10 µM) |
| FG-N | 122.8 (8 mg/mL) | - | - | - |
| FSFG (2mer) | 82.2 (800 µM) | - | - | - |
| GLFG | 132.2 | 107<br>(325 µM in 92% D2O) | - | - |
| Nup100FG | 186 | - | - | - |
| Nsp1FG | 241 | - | - | - |
| Nsp1 (full length, 615 AA, 19 FxFG motifs, 14 other FG motifs); 5.1 pmol/cm$^2$ grafting density; at different Impß concentrations | - | - | - | 240-340 (0 µM) (<br>280-380 Å (10 µM) |
| Kap95-FSFG6x2 (chain A/B) | - | - | 107.1<br>105.3 | - |
| FSFG6 in Kap95-FSFG6 | - | - | 108.5 | - |
| MBP-FSFG6 | - | - | 111.5 | - |
| Kap95-GLFGnup100 (3500 ns) | - | - | 78.7 | - |

| | SAXS [Å] | SANS [Å] | MD | QCM-D film thickness [Å] (NTR concentration) |
|---|---|---|---|---|
| Kap95-GLFGnup100x2 (chain A/B) | - | - | 70.4<br>70.2 | - |
| NTF2-FSFG6 high salt (500 ns) | - | - | 98.1 | - |
| NTF2-GLFGnup100 high salt | - | - | 67.6 | - |
| FSFG6 free low salt (218 ns) | - | - | 107.0 | - |
| FSFG2 in NTF2:FSFG2 low salt (1466 ns) | - | - | 74.4 | - |
| Nup49 (RH design) | 163 | - | - | - |
| Nup1FG(352-1076)-LEHHHH Ryo+Sam construct | 354 | - | - | - |

**Supplementary Table S3.** Persistence length for FSFG constructs from SANS[11], with analysis following ref.[63].

| | **Kuhn length (2*persistence length) [Å]** | **Contour Length [Å]** |
|---|---|---|
| FSFG-K (Nsp1) 8 mg/mL | 19.06 ± 0.74 | 471.5 ± 16.1 |
| FSFG (2mer) 800 μM | 15 ± 2.77 | 284.8 ± 44.39 |
| GLFG | 12.99 | 577.7 |
| Nsp1FG | 26.7 ± 1.7 | 1299 ± 75.92 |

**Supplementary Table S4A.** $K_D$ values from nuclear magnetic resonance (NMR) and isothermal titration calorimetry (ITC) for NTF2[9]. All constructs except FGFG12 contain 6 repeats, as described in Hayama *et al.* 2018.

| Construct | NMR $K_D$ [mM] | ITC $K_D$ [mM] | Joint Analysis | n |
|---|---|---|---|---|
| FSFG1:NTF2 | 4.35 ± 0.20 | 2.73 ± 0.57 | 3.54 ± 0.79 | - |
| FSFG2:NTF2 | 2.03 ± 0.07 | 1.93 ± 0.38 | 1.98 ± 0.39 | - |
| FSFG3:NTF2 | 1.19 ± 0.04 | 1.02 ± 0.18 | 1.11 ± 0.19 | - |
| FSFG4:NTF2 | 0.86 ± 0.03 | 0.62 ± 0.08 | 0.74 ± 0.12 | 1 |
| FSFG5:NTF2 | 0.64 ± 0.02 | 0.52 ± 0.08 | 0.58 ± 0.09 | 1 |
| FSFG6:NTF2 | 0.53 ± 0.01 | 0.53 ± 0.07 | 0.53 ± 0.07 | 1 |
| FSFG12:NTF2 | 0.37 ± 0.01 | 0.42 ± 0.03 | 0.34 ± 0.04 | 2 |
| FSFSSS* | 2.13 ± 0.10 | - | - | - |
| FSSFSS* | 3.27 ± 0.23 | - | - | - |
| FSSSSF* | 3.49 ± 0.18 | - | - | - |

* six repeats with S indicating a repeat where FSFG was mutated to SSSG

**Supplementary Table S4B**. Joint analysis of $K_D$ measurements.
The analysis is based on both NMR and ITC for NTF2[9]

| **Construct** | **$K_D$ [mM]** | **SEM** | **95% CI min** | **95% CI max** |
|---|---|---|---|---|
| FSFG12:NTF2 | 0.4307 | 0.018 | 0.3946 | 0.4667 |
| FSFG1:NTF2 | 3.4264 | 0.2443 | 2.9438 | 3.9091 |
| FSFG2:NTF2 | 1.7284 | 0.09 | 1.5502 | 1.9065 |
| FSFG3:NTF2 | 1.1784 | 0.052 | 1.0754 | 1.2814 |
| FSFG4:NTF2 | 0.8351 | 0.0397 | 0.7555 | 0.9147 |
| FSFG4:NTF2* | 0.87945 | 0.039824 | - | - |
| FSFG5:NTF2* | 0.5563 | 0.0172 | 0.522 | 0.5905 |
| FSFG5:NTF2* | 0.59061 | 0.0162 | - | - |
| FSFG6:NTF2* | 0.3659 | 0.0191 | 0.3283 | 0.4035 |
| FSFG12:NTF2* | 0.43066 | 0.018042 | - | - |

* alternative estimate using extended model for concentration normalization

**Supplementary Table S5.** Measured and simulated diffusion coefficients in buffer.

| | Measured diffusion coefficient [$Å^2$/ns] | *D* from all-atom MD simulations on Anton [$Å^2$/ns] *without box size correction* | *D* from MD with box-size correction [$Å^2$/ns]<br>*With correction for bounding box side* length $L_B$<br>$D = 695.113/L_B$ $Å^2$/ns |
|---|---|---|---|
| **FSFG-K** | 8.07 +/- 0.033<br>Infinite dilution<br>(Rs=29.6 Å, or 26.2 Å based on infinite dilution) | 1.29 (lag time 30 ns, FSFG6_FREE_TIP4PD, low salt) | 8.43 ($L_B$=97.35 Å; $R_S$=28.05 Å) |
| **FSFG2 - free (2mer)** | - | - | - |
| **FG-N** | 8.75 +/- .263<br>Infinite dilution (Rh=24.2 Å) | - | - |
| **NTF2 - free** | 8.80 (Rh= 27.3 Å) single point measured | - | - |
| **Kap95 – free (5mg/mL) (obtained 11/18/2013)** | 5.085 (Rh = 47 Å) | - | - |
| **FSFG-K in presence of Kap95, 150 mM salt** | 3.153 (Rh = 76.3 Å)<br>FSFG:Kap95 1:1 molar ratio | 1.179 +- 0.040 ($R_S$ = 199 Å) at lag=206.64 ns | 5.895 ($L_B$=147.38 Å; $R_S$=40.1 Å; Rg=33.2 Å) |
| **FSFG-Kx2:Kap95 (chain A/B)** | - | 0.934 +- 0.024<br>0.836 +- 0.036 | 5.51 +- -0.024 (La=151.90 Å; Rs=42.9 Å; Rg=33.0 Å)<br>5.412 +- 0.024 (La=151.90 Å; Rs=43.7 Å; Rg=31.9 Å) |
| **FSFG-K in presence of NTF2** | 6.72 (Rh = 35.9 Å)<br>FSFG:NTF2 1:2 molar ratio | - | - |
| **FSFG6 free from MBP:FSFG6 simulation (frame 1300-4700, 0.96 ns per frame)** | - | 1.275 ($R_S$=186 Å) | 6.942 ($L_B$=122.65 Å; $R_S$=34.2 Å) |
| **FSFG6 free from Kap95:FSFG6** | - | 2.220 ($R_S$=107 Å) at lag=50.4 n | 6.936 ($L_B$=147.38 Å; $R_S$=34.24 Å; |

| | Measured diffusion coefficient [Å$^2$/ns] | *D* from all-atom MD simulations on Anton [Å$^2$/ns] *without box size correction* | *D* from MD with box-size correction [Å$^2$/ns]<br>*With correction for bounding box side* length $\boldsymbol{L_B}$<br>**D =** *695.113/*$L_B$ Å$^2$/ns |
|---|---|---|---|
| **simulation (frame 625-825, 5.04 ns per frame) 150 mM salt** | | | Rg=31.987) |
| **GLFG-Nup100 free from Kap95:GLFG simulations (frame 0-50, 0.96 ns per frame)** | - | 2.60 ($R_S$=91.36 Å) at lag=19.2 ns | 7.455 ($L_B$=143.16 Å; $R_S$=31.86 Å) |

**Supplementary Table S6.** Polymer chain relaxation time and shape parameters from MD simulations.
Relaxation time tau, center-of-mass (c.o.m.) distance, $R_g$, Cα(N)-Cα(C) distance, end-to-end distance and Flory coefficient for various constructs and simulation conditions, estimated from atomistic molecular dynamics simulations on the Anton[6] and Anton2 (here) supercomputers.

| | Tau over adjacent pairs of 20-res beads [ns] (95% interval) | Tau end-to-end [ns] over 6 x 20 residue beads | 20-20 mean c.o.m. length [Å] (std-dev) | $R_g$ of 20 residues | $R_g$ of 40 residues | $R_g$ of 120 residues | $R_g$ of all residues [std-dev] | Maximal Cα(N)-Cα(C) distance [Å] | Mean Cα(N)-Cα(C) distance [Å] (std-dev) | Mean end-to-end distance with 20 res beads [Å] (std-dev) | Flory coefficient from $R_g$ vs. length in 10 res intervals (95% intervals) |
|---|---|---|---|---|---|---|---|---|---|---|---|
| Kap95-FSFG 6x2 (chain A/B) | 223.2<br>(132-350)<br>324<br>(182-461) | 670.6<br>385.4 | 27.39 (8.89)<br>27.75 (9.42) | 12.7<br>13.0 | 18.2<br>18.0 | 32.1<br>32.8 | 31.2 [6.2]<br>33.4 [5.5]<br>(125 res) | - | - | - | 0.53 (+-0.02)<br>0.57 (+-0.03) |
| Kap95-FSFG 6 | 297.4<br>(165-675) | 226.8 | 25.69 (9.10)<br>Skewness<br>-0.01 | 12.4 | 17.8 | 33.4 | 33.2 [6.8]<br>(125 res) | 175.7 | 81.1 (65.2) | 71.8 (24.5) | 0.58 (+-0.02) |
| MBP-FSFG6 | 122.4<br>(83-166) | 93.6 | 27.87 (9.33)<br>Skewness<br>-0.05 | 12.8 | 18.2 | 35.7 | 36.4 [7.1]<br>(125 res) | 203.2 | 83.8 (69.0) | 77.2 (26.4) | 0.60 (+-0.02) |
| Kap95-GLF Gnup100 (3500 ns) | 142.6<br>(100.3-269.3) | 195.4 | 17.78 (5.79)<br>Skewness<br>0.32 | 11.3 | 15.5 | 21.3 | 22.2 [1.8]<br>(132 res) | 83.5 | 54.2 (83.5) | 36.5 (5.5) | 0.4 (+-0.05) |
| Kap95-GLF Gnup100x2 (chain A/B) | 227.0<br>(132.0-406.6)<br>301.0<br>(184.8-807.8) | 279.8<br>216.5 | 16.74 (4.97)<br><br>17.68 (7.60)<br>Skewness<br>0.39/0.46 | 9.7<br><br>11.1 | 13.9<br><br>13.9 | 19.3<br><br>18.3 | 19.7 [1.4]<br><br>19.3 [1.7]<br>(132 res) | 58.7<br><br>71.2 | 41.7 (34.0)<br><br>31.6 (26.4) | 35.4 (3.0)<br><br>20.7 (7.4) | 0.42 (+-0.05)<br>0.33 (+-0.05) |
| NTF2-FSFG 6 high salt (500 ns) | 26.4<br>(22.1-47.5) | 15.8 | 21.13 (8.16) | 12.4 | 16.1 | 28.1 | 28.3<br>(125 res) | - | - | - | 0.50 (+-0.04) |
| NTF2-GLFG nup100 high salt | 100.3<br>(58.1-132.0) | 89.8 | 17.47 (5.95) | 10.0 | 14.4 | 19.6 | 19.1<br>(132 res) | - | - | - | 0.43 (+-0.08) |
| FSFG6 free low salt (218 ns) | 26.4<br>(21.3-37.0) | 37.0 | 24.61 (9.55) | 12.4 | 16.4 | 30.0 | 31.5<br>(125 res) | - | 87.6 (70.5) | 72.3 | 0.51 (+-0.05) |
| NTF2:FSFG 2 low salt, large box (1466 ns) | 55.4 | - | 24.19 (9.01) | 13.0 | 17.8 | - | 21.5<br>(54 res) | - | - | - | 0.52 (+-0.05) |

**Supplementary Table S7**. Details on SANS measurements of $R_g$ and maximal dimension $D_{max}$ [11]; see also **Supplementary Table S2**.

| Sample | Debye analysis | P(r) | |
|---|---|---|---|
| | $R_g$ [Å] | $R_g$ [Å] | $D_{max}$ [Å] |
| [$^2$H]-FSFG-K [0.6 mM] (8mg/ml) | 36.2 ± 0.3 | 35.8 ± 1.1 | 127.0 |
| [$^2$H]-FSFG-K [0.6 mM] Kap95 [0.25 mM] | 39.6 ± 0.8 | 42.3 ± 2.7 | 147.0 |
| [$^2$H]-FSFG-K [0.6 mM] Kap95 [0.5 mM] | 41.8 ± 1.2 | 44.2 ± 1.7 | 143.5 |
| [$^2$H]-FSFG-K [0.6 mM] NTF2 [0.3 mM] | 41.2 ± 0.3 | 39.8 ± 1.9 | 142.8 |
| [$^2$H]-FSFG-K [0.6 mM] NTF2 [0.6 mM] | 45.6 ± 0.5 | 42.2 ± 2.2 | 151.0 |
| [$^2$H]-FSFG-K [0.6 mM] NTF2 [1.2 mM] | 47.9 ± 0.6 | 46.4 ± 2.9 | 168.0 |

**Supplementary Table S8.** Ensemble modeling data of $R_g$ and $D_{max}$ based on SANS[11].

| Sample | $\chi^2$ | Selected ensemble $R_g$ [Å] | Selected ensemble $D_{max}$ [Å] | Selected ensemble end-to-end distance (Å) Cα(N)-Cα(C) |
|---|---|---|---|---|
| [$^2$H]-FSFG-K [0.6 mM] (8mg/ml) | 0.20 | 35.5 | 87.4 | 73.7 |
| [$^2$H]-FSFG-K[ 0.6 mM] Kap95 [0.25 mM] | 0.26 | 46.9 | 129.1 | 118.0 |
| [$^2$H]-FSFG-K [0.6 mM] Kap95 [0.5 mM] | 0.38 | 47.0 | 137.4 | 135.0 |
| [$^2$H]-FSFG-K [0.6 mM] NTF2 [0.3 mM] | 0.17 | 38.5 | 119.0 | 119.0 |
| [$^2$H]-FSFG-K [0.6 mM] NTF2 [0.6 mM] | 0.15 | 40.7 | 117.6 | 115.3 |
| [$^2$H]-FSFG-K [0.6 mM] NTF2 [1.2 mM] | 0.27 | 42.0 | 141.0 | 128.1 |
| FSFG-K [375 mM] + [$^2$H]-Kap95 [75 µM] | 0.99 | 46.6 | 120.9 | 108.3 |

**Supplementary Table S9.** Constructs used in LH and SS-RH SAXS.

| Abbreviation | Simple Structure | Sequence | #AA | MW [Da] |
|---|---|---|---|---|
| (FSFG-K-274-397)-6His | M-Nsp1(274-397)-LEHHHHHH | MDNKTTNTTPSFSFGAKSDENKAGATSKPAFSFG<br>AKPEEKKDDNSSKPAFSFGAKSNEDKQDGTAKPAFSFGAKPAEKNN<br>NETSKPAF<br>SFGAKSDEKKDGDASKPAFSFGAKPDENKASATSKPALEHHHHHH | 133 | 14,135 |
| (FG-N)-6His | MGT-Nsp1(48-172)-SHMHHHHHH | MGTSAPNNTNNANSSITPAFGSNNTGNTAFGNSNPTSNVFGSNNST<br>TNTFGSNSAGTSLFGSSSAQQTKSNGTAGGNTFGSSSLFNNSTNSN<br>TTKPAFGGLNFGGGNNTTPSSTGNANTSNNLFGATASHMHHHHHH | 137 | 13,649 |
| (FG-N)-(FSFG-K)-6His | MGT-Nsp1(48-172)-ASATSKPA-Nsp1(284-397)-SHHHHHH | MGTSAPNNTNNANSSITPAFGSNNTGNTAFGNSNPTSNVFGSNNST<br>TNTFGSNSAGTSLFGSSSAQQTKSNGTAGGNTFGSSSLFNNSTNSN<br>TTKPAFGGLNFGGGNNTTPSSTGNANTSNNLFGATAASATSKPAFS<br>FGAKSDENKAGATSKPAFSFGAKPEEKKDDNSSKPAFSFGAKSNED<br>KQDGTAKPAFSFGAKPAEKNNNETSKPAFSFGAKSDEKKDGDASKP<br>AFSFGAKPDENKASATSKPASHHHHHH | 257 | 25,955 |
| (FG-N)-(FSFG-K)-Tet-6His | MGT-Nsp1(48-172)-ASATSKPA-Nsp1(284-397)-SHMGEYFTLQIRGRERFEMFRELNEALELKDAQAHMHHHHHH | MGTSAPNNTNNANSSITPAFGSNNTGNTAFGNSNPTSNVFGSNNST<br>TNTFGSNSAGTSLFGSSSAQQTKSNGTAGGNTFGSSSLFNNSTNSN<br>TTKPAFGGLNFGGGNNTTPSSTGNANTSNNLFGATAASATSKPAFS<br>FGAKSDENKAGATSKPAFSFGAKPEEKKDDNSSKPAFSFGAKSNED<br>KQDGTAKPAFSFGAKPAEKNNNETSKPAFSFGAKSDEKKDGDASKP<br>AFSFGAKPDENKASATSKPASHMGEYFTLQIRGRERFEMFRELNEA<br>LELKDAQAHMHHHHHH | 292 | 30,235 |
| 2mer | M-Nsp1(346-397)-HHHHHH | MAKPAEKNNNETSKPAFSFGAKSDEKKDGDASKPAFSFGAKPDENK<br>ASATSKPAHHHHHH | 60 | 6,455 |

| Abbreviation | Simple Structure | Sequence | #AA | MW [Da] |
|---|---|---|---|---|
| Nsp1FG | M-Nsp1(30-591) | MSTGAGAFGTGQSTFGFNNSAPNNTNNANSSITPA<br>FGSNNTGNTAFGNSNPTSNVFGSNNSTTNT<br>FGSNSAGTSLFGSSSAQQTKSNGTAGGNT<br>FGSSSLFNNSTNSNTTKPAFGGLNFGGGNNTTPSSTGNANTSNNLF<br>GATANANKPAFSFGATTNDDKKTEPDKPAFSFNSSVGNKTDAQAPT<br>TGFSFGSQLGGDKTVNEAAKPSLSFGSGSAGANPAGASQPEPTTNE<br>PAKPALSFGTATSDNKTTNTTPSFSFGAKSDENKAGATSKPAFSFG<br>AKPEEKKDDNSSKPAFSFGAKSNEDKQDGTAKPAFSFGAKPAEKNN<br>NETSKPA<br>FSFGAKSDEKKDGDASKPAFSFGAKPDENKASATSKPA<br>FSFGAKPEEKKDDNSSKPAFSFGAKSNEDKQDGTAKPA<br>FSFGAKPAEKNNNETSKPAFSFGAKSDEKKDGDASKPAFSFGAKSD<br>EKKDSDSSKPAFSFGTKSNEKKDSGSSKPA<br>FSFGAKPDEKKNDEVSKPAFSFGAKANEKKESDESKSA<br>FSFGSKPTGKEEGDGAKAAISFGAKPEEQKSSDTSKPA<br>FTFGLEHHHHHH | 571 | 58,494 |
| GLFG | M-Nup100(318-444)SNLEHHHHHH | MSLFGKANTFSNSASGGLFGQNNQQQGSGLFGQNSQTSGSSGLFGQ<br>NNQKQPNTFTQSNTGIGLFGQNNNQQQQSTGLFGAKPAGTTGSLFG<br>GNSSTQP<br>NSLFGTTNVPTSNTQSQQGNSLFGATKLTSNLEHHHHHH | 138 | 14,348 |
| Nup100FG | pET21b-Nup100FG-(1-570)-CHHHHHH | MFGNNRPMFGGSNLSFGSNTSSFGGQQSQQPNSLFGNSNNNNNSTSN<br>NAQSGFGGFTSAAGSNSNSLFGN<br>NNTQNNGAFGQSMGATQNSPFGSLNSSNASNGNTFGGSSSMGSFGGNT<br>NNAFNNNSNSTNSPFGFNKPNT<br>GGTLFGSQNNNSAGTSSLFGGQSTSTTGTFGNTGSSFGTGLNGNGSNI<br>FGAGNNSQSNTTGSLFGNQQSS<br>AFGTNNQQGSLFGQQSQNTNNAFGNQNQLGGSSFGSKPVGSGSLFGQS<br>NNTLGNTTNNRNGLFGQMNSSN<br>QGSSNSGLFGQNSMNSSTQGVFGQNNNQMQINGNNNNSLFGKANTFSN<br>SASGGLFGQNNQQQGSGLFGQN<br>SQTSGSSGLFGQNNQKQPNTFTQSNTGIGLFGQNNNQQQQSTGLFGAK<br>PAGTTGSLFGGNSSTQPNSLFG<br>TTNVPTSNTQSQQGNSLFGATKLTNMPFGGNPTANQSGSGNSLFGTKP<br>ASTTGSLFGNNTASTTVPSTNG<br>LFGNNANNSTSTTNTGLFGAKPDSQSKPALGGGLFGNSNSNSSTIGQN<br>KPVFGGTTQNTGLFGATGTNSS<br>AVGSTGKLFG CHHHHHH | 577 | 57,953 |

| Abbreviation | Simple Structure | Sequence | #AA | MW [Da] |
|---|---|---|---|---|
| Nup49 | M-Nup49-(1-269<br>) | MFGLNKASSTPAGGLFGQASGASTGNANTGFSFGGTQTGQNTGPSTGG<br>LFGAKPAGSTGGLGASFGQQQQQSQTNAFGGSATTGGGLFGNKPNNTA<br>NTGGGLFGANSNSNSGSLFGSNNAQTSRGLFGNNNTNNINNSSSGMNN<br>ASAGLFGSKPAGGTSLFGNTSTSSAPAQNQGMFGAKPAGTSLFGNNAG<br>NTTTGGGLFGSKPTGATSLFGSSNNNNNNNNSNNIMSASGGLFGNQQQ<br>QLQQQPQMQCALQNLSQLPITPMTRISELLEHHHHHH | 277 | 27,421 |
| Nup1FG | M-Nup1-(352-10<br>76)-LEHHHHHH | MKATSSAGAVFKSSVEMGKTDKSTKTAEAPTLSFNFSQKANKTKAVDN<br>TVPSTTLFNFGGKSDTVTSASQPFKFGKTSEKSENHTESDAPPKSTAP<br>IFSFGKQEENGDEGDDENEPKRKRRLPVSEDTNTKPLFDFGKTGDQKE<br>TKKGESEKDASGKPSFVFGASDKQAEGTPLFTFGKKADVTSNIDSSAQ<br>FTFGKAATAKETHTKPSETPATIVKKPTFTFGQSTSENKISEGSAKPT<br>FSFSKSEEERKSSPISNEAAKPSFSFPGKPVDVQAPTDDKTLKPTFSF<br>TEPAQKDSSVVSEPKKPSFTFASSKTSQPKPLFSFGKSDAAKEPPGSN<br>TSFSFTKPPANETDKRPTPPSFTFGGSTTNNTTTTSTKPSFSFGAPES<br>MKSTASTAAANTEKLSNGFSFTKFNHNKEKSNSPTSFFDGSASSTPIP<br>VLGKPTDATGNTTSKSAFSFGTANTNGTNASANSTSFSFNAPATGNGT<br>TTTSNTSGTNIAGTFNVGKPDQSIASGNTNGAGSAFGFSSSGTAATGA<br>ASNQSSFNFGNNGAGGLNPFTSATSSTNANAGLFNKPPSTNAQNVNVP<br>SAFNFTGNNSTPGGGSVFNMNGNTNANTVFAGSNNQPHQSQTPSFNTN<br>SSFTPSTVPNINFSGLNGGITNTATNALRPSDIFGANAASGSNSNVTN<br>PSSIFGGAGGVPTTSFGQPQSAPNQMGMGTNNGMSMGGGVMANRKIAR<br>MRHSKRLEHHHHHH | 734 | 76,082 |

**Supplementary Table S10.** Parameters for FG Nups following model training in the coarse-grained model of transport.

| Nup | subregion | Residue range | Self-k [kcal/mol/Å] | self-range | Kap-k [kcal/mol/A$^2$] | Kap-range [Å] | Non-specific k | Non-specific range | Backbone-k ($k_2$) [kcal/mol/Å$^2$] | Backbone tau [ns] |
|---|---|---|---|---|---|---|---|---|---|---|
| Nsp1 | N | 1-180 | 1.47 | 6.00 | 2.64 | 5.5 | 0.08 | 5.00 | 0.0075 | 50 |
| | C | 181-550 | 1.32 | 6.00 | 2.64 | 5.5 | 0.01 | 5.00 | 0.0075 | 50 |
| | s | 551-636 | 1.28 | 6.00 | - | - | 0.01 | 5.00 | 0.0075 | 50 |
| Nup100 | | 2-610 | 1.47 | 6.00 | 2.64 | 5.5 | 0.08 | 5.00 | 0.0075 | 50 |
| | s | 551-800 | 1.28 | 6.00 | - | - | 0.01 | 5.00 | 0.0075 | 50 |
| Nup116 | | 1-751 | 1.47 | 6.00 | 2.64 | 5.5 | 0.08 | 5.00 | 0.0075 | 50 |
| | s | 751-950 | 1.28 | 6.00 | - | - | 0.01 | 5.00 | 0.0075 | 50 |
| Nup159 | | 442-881 | 1.32 | 6.00 | 2.64 | 5.5 | 0.07 | 5.00 | 0.0075 | 50 |
| | s | 882-1116 | 1.28 | 6.00 | - | - | 0.01 | 5.00 | 0.0075 | 50 |
| Nup49 | | 1-240 | 1.47 | 6.00 | 2.64 | 5.5 | 0.08 | 5.00 | 0.0075 | 50 |
| | s | 241-269 | 1.28 | 6.00 | - | - | 0.01 | 5.00 | 0.0075 | 50 |
| Nup57 | | 1-200 | 1.47 | 6.00 | 2.64 | 5.5 | 0.08 | 5.00 | 0.0075 | 50 |
| | s | 201-286 | 1.28 | 6.00 | - | - | 0.01 | 5.00 | 0.0075 | 50 |
| Nup145N | | 1-200 | 1.47 | 6.00 | 2.64 | 5.5 | 0.08 | 5.00 | 0.0075 | 50 |
| | s | 201-250 | 1.28 | 6.00 | - | - | 0.01 | 5.00 | 0.0075 | 50 |
| Nup1 | s | 201-325 | 1.28 | 6.00 | - | - | 0.01 | 5.00 | 0.0075 | 50 |
| | m | 326-797 | 1.32 | 6.00 | 5.98 | 5.5 | 0.01 | 5.00 | 0.0075 | 50 |

| Nup | subregion | Residue range | Self-k [kcal/mol/Å] | self-range | Kap-k [kcal/mol/A$^2$] | Kap-range [Å] | Non-specific k | Non-specific range | Backbone -k ($k_2$) [kcal/mol/Å$^2$] | Backbone tau [ns] |
|---|---|---|---|---|---|---|---|---|---|---|
| | c | 798-1076 | 1.47 | 6.00 | 5.98 | 5.5 | 0.08 | 5.00 | 0.0075 | 50 |
| Nup60 | | 399-539 | 1.32 | 6.00 | 5.98 | 5.5 | 0.01 | 5.00 | 0.0075 | 50 |
| | s | 301-398 | 1.28 | 6.00 | - | - | 0.01 | 5.00 | 0.0075 | 50 |

**Supplementary Table S11.** Sequences of FG Nup domains used in **Extended Data Fig. 1**

NUP100

```
>NUP100 YKL068W SGDID:S000001551
  1   50 MFGNNRPMFGGSNLSFGSNTSSFGGQQSQQPNSLFGNSNNNNNSTSNNAQ
 51  100 SGFGGFTSAAGSNSNSLFGNNNTQNNGAFGQSMGATQNSPFGSLNSSNAS
101  150 NGNTFGGSSSMGSFGGNTNNAFNNNSNSTNSPFGFNKPNTGGTLFGSQNN
151  200 NSAGTSSLFGGQSTSTTGTFGNTGSSFGTGLNGNGSNIFGAGNNSQSNTT
201  250 GSLFGNQQSSAFGTNNQQGSLFGQQSQNTNNAFGNQNQLGGSSFGSKPVG
251  300 SGSLFGQSNNTLGNTTNNRNGLFGQMNSSNQGSSNSGLFGQNSMNSSTQG
301  350 VFGQNNNQMQINGNNNNSLFGKANTFSNSASGGLFGQNNQQQGSGLFGQN
351  400 SQTSGSSGLFGQNNQKQPNTFTQSNTGIGLFGQNNNQQQQSTGLFGAKPA
401  450 GTTGSLFGGNSSTQPNSLFGTTNVPTSNTQSQQGNSLFGATKLTNMPFGG
451  500 NPTANQSGSGNSLFGTKPASTTGSLFGNNTASTTVPSTNGLFGNNANNST
501  550 STTNTGLFGAKPDSQSKPALGGGLFGNSNSNSSTIGQNKPVFGGTTQNTG
551  600 LFGATGTNSSAVGSTGKLFGQNNNTLNVGTQNVPPVNNTTQNALLGTTAV
601  650 PSLQQAPVTNEQLFSKISIPNSITNPVKATTSKVNADMKRNSSLTSAYRL
651  700 APKPLFAPSSNGDAKFQKWGKTLERSDRGSSTSNSITDPESSYLNSNDLL
701  750 FDPDRRYLKHLVIKNNKNLNVINHNDDEASKVKLVTFTTESASKDDQASS
751  800 SIAASKLTEKAHSPQTDLKDDHDESTPDPQSKSPNGSTSIPMIENEKISS
801  850 KVPGLLSNDVTFFKNNYYISPSIETLGNKSLIELRKINNLVIGHRNYGKV
851  900 EFLEPVDLLNTPLDTLCGDLVTFGPKSCSIYENCSIKPEKGEGINVRCRV
901  950 TLYSCFPIDKETRKPIKNITHPLLKRSIAKLKENPVYKFESYDPVTGTYS
951  960 YTIDHPVLTHHHHHH
```

NUP1 FG Domain

```
>NUP1 YOR098C SGDID:S000005624
  1   50 MSSNTSSVMSSPRVEKRSFSSTLKSFFTNPNKKRPSSKKVFSSNLSYANH
 51  100 LEESDVEDTLHVNKRKRVSGTSQHSDSLTQNNNNAPIIIYGTENTERPPL
101  150 LPILPIQRLRLLREKQRVRNMRELGLIQSTEFPSITSSVILGSQSKSDEG
151  200 GSYLCTSSTPSPIKNGSCTRQLAGKSGEDTNVGLPILKSLKNRSNRKRFH
201  250 SQSKGTVWSANFEYDLSEYDAIQKKDNKDKEGNAGGDQKTSENRNNIKSS
251  300 ISNGNLATGPNLTSEIEDLRADINSNRLSNPQKNLLLKGPASTVAKTAPI
301  350 QESFVPNSERSGTPTLKKNIEPKKDKESIVLPTVGFDFIKDNETPSKKTS
351  400 PKATSSAGAVFKSSVEMGKTDKSTKTAEAPTLSFNFSQKANKTKAVDNTV
401  450 PSTTLFNFGGKSDTVTSASQPFKFGKTSEKSENHTESDAPPKSTAPIFSF
451  500 GKQEENGDEGDDENEPKRKRRLPVSEDTNTKPLFDFGKTGDQKETKKGES
501  550 EKDASGKPSFVFGASDKQAEGTPLFTFGKKADVTSNIDSSAQFTFGKAAT
551  600 AKETHTKPSETPATIVKKPTFTFGQSTSENKISEGSAKPTFSFSKSEEER
601  650 KSSPISNEAAKPSFSFPGKPVDVQAPTDDKTLKPTFSFTEPAQKDSSVVS
651  700 EPKKPSFTFASSKTSQPKPLFSFGKSDAAKEPPGSNTSFSFTKPPANETD
701  750 KRPTPPSFTFGGSTTNNTTTTSTKPSFSFGAPESMKSTASTAAANTEKLS
```

```
 751  800 NGFSFTKFNHNKEKSNSPTSFFDGSASSTPIPVLGKPTDATGNTTSKSAF
 801  850 SFGTANTNGTNASANSTSFSFNAPATGNGTTTTSNTSGTNIAGTFNVGKP
 851  900 DQSIASGNTNGAGSAFGFSSSGTAATGAASNQSSFNFGNNGAGGLNPFTS
 901  950 ATSSTNANAGLFNKPPSTNAQNVNVPSAFNFTGNNSTPGGGSVFNMNGNT
 951 1000 NANTVFAGSNNQPHQSQTPSFNTNSSFTPSTVPNINFSGLNGGITNTATN
1001 1050 ALRPSDIFGANAASGSNSNVTNPSSIFGGAGGVPTTSFGQPQSAPNQMGM
1051 1076 GTNNGMSMGGGVMANRKIARMRHSKRHHHHHH
```

NSP49 FG domain

```
NUP49 YGL172W SGDID:S000003140
   1   50 MFGLNKASSTPAGGLFGQASGASTGNANTGFSFGGTQTGQNTGPSTGGLF
  51  100 GAKPAGSTGGLGASFGQQQQQSQTNAFGGSATTGGGLFGNKPNNTANTGG
 101  150 GLFGANSNSNSGSLFGSNNAQTSRGLFGNNNTNNINNSSSGMNNASAGLF
 151  200 GSKPAGGTSLFGNTSTSSAPAQNQGMFGAKPAGTSLFGNNAGNTTTGGGL
 201  250 FGSKPTGATSLFGSSNNNNNNNNSNNIMSASGGLFGNQQQQLQQQPQMQC
 251  277 ALQNLSQLPITPMTRISELLGHHHHHH
```

NSP1 FG

>NSP1 YJL041W SGDID:S000003577

```
  1  50 MNFNTPQQNKTPFSFGTANNNSNTTNQNSSTGAGAFGTGQSTFGFNNSAP
 51 100 NNTNNANSSITPAFGSNNTGNTAFGNSNPTSNVFGSNNSTTNTFGSNSAG
101 150 TSLFGSSSAQQTKSNGTAGGNTFGSSSLFNNSTNSNTTKPAFGGLNFGGG
151 200 NNTTPSSTGNANTSNNLFGATANANKPAFSFGATTNDDKKTEPDKPAFSF
201 250 NSSVGNKTDAQAPTTGFSFGSQLGGNKTVNEAAKPSLSFGSGSAGANPAG
251 300 ASQPEPTTNEPAKPALSFGTATSDNKTTNTTPSFSFGAKSDENKAGATSK
301 350 PAFSFGAKPEEKKDDNSSKPAFSFGAKSNEDKQDGTAKPAFSFGAKPAEK
351 400 NNNETSKPAFSFGAKSDEKKDGDASKPAFSFGAKPDENKASATSKPAFSF
401 450 GAKPEEKKDDNSSKPAFSFGAKSNEDKQDGTAKPAFSFGAKPAEKNNNET
451 500 SKPAFSFGAKSDEKKDGDASKPAFSFGAKSDEKKDSDSSKPAFSFGTKSN
501 550 EKKDSGSSKPAFSFGAKPDEKKNDEVSKPAFSFGAKANEKKESDESKSAF
551 600 SFGSKPTGKEEGDGAKAAISFGAKPEEQKSSDTSKPAFTFGAQKDNEKKT
601 650 EESSTGKSTADVKSSDSLKLNSKPVELKPVSLDNKTLDDLVTKWTNQLTE
651 700 SASHFEQYTKKINSWDQVLVKGGEQISQLYSDAVMAEHSQNKIDQSLQYI
701 750 ERQQDELENFLDNFETKTEALLSDVVSTSSGAAANNNDQKRQQAYKTAQT
751 800 LDENLNSLSSNLSSLIVEINNVSNTFNKTTNIDINNEDENIQLIKILNSH
801 824 FDALRSLDDNSTSLEKQINSIKKHHHHHH
```
